# Dynamic interplay between RPL3- and RPL3L-containing ribosomes modulates mitochondrial activity in the mammalian heart

**DOI:** 10.1101/2021.12.04.471171

**Authors:** Ivan Milenkovic, Helaine Graziele Santos Vieira, Morghan C Lucas, Jorge Ruiz-Orera, Giannino Patone, Scott Kesteven, Jianxin Wu, Michael Feneley, Guadalupe Espadas, Eduard Sabidó, Norbert Hubner, Sebastiaan van Heesch, Mirko Voelkers, Eva Maria Novoa

## Abstract

The existence of naturally occurring ribosome heterogeneity is now a well-acknowledged phenomenon. However, whether this heterogeneity leads to functionally diverse ‘specialized ribosomes’ is still a controversial topic. Here, we explore the biological function of RPL3L (uL3L), a ribosomal protein (RP) paralog of RPL3 (uL3) that is exclusively expressed in muscle and heart tissues, by generating a viable homozygous *Rpl3l* knockout mouse strain. We identify a rescue mechanism in which, upon RPL3L depletion, RPL3 becomes upregulated, yielding RPL3-containing ribosomes instead of RPL3L-containing ribosomes that are typically found in cardiomyocytes. Using both ribosome profiling (Ribo-Seq) and a novel orthogonal approach consisting of ribosome pulldown coupled to nanopore sequencing (Nano-TRAP), we find that RPL3L neither modulates translational efficiency nor ribosome affinity towards a specific subset of transcripts. By contrast, we show that depletion of RPL3L leads to increased ribosome-mitochondria interactions in cardiomyocytes, which is accompanied by a significant increase in ATP levels, potentially as a result of mitochondrial activity fine-tuning. Our results demonstrate that the existence of tissue-specific RP paralogs does not necessarily lead to enhanced translation of specific transcripts or modulation of translational output. Instead, we reveal a complex cellular scenario in which RPL3L modulates the expression of RPL3, which in turn affects ribosomal subcellular localization and, ultimately, mitochondrial activity.

## INTRODUCTION

A major challenge in biology is to comprehend how protein synthesis is regulated with high precision, both in the spatial and the temporal context. While many layers of gene expression regulation have been extensively studied over the years, ribosomes have been historically perceived as static and passive elements that do not partake in regulatory processes. Several works, however, have challenged this view (1–6), and have provided evidence that ribosomes with specialized functions exist, and can preferentially translate specific subsets of mRNAs (7). While naturally occurring ribosome heterogeneity is now a well-documented phenomenon (8–10), whether this heterogeneity leads to functionally diverse ‘specialized ribosomes’ is still a controversial topic (11–14).

Ribosomes are supramolecular ribonucleoprotein complexes responsible for protein synthesis in all known organisms. Eukaryotic ribosomes consist of two subunits: the small 40S subunit, which is made up of 32 ribosomal proteins (RPs) and the 18S ribosomal RNA (rRNA), and the large 60S subunit, comprised of 47 RPs and the 5S, 5.8S and 28S rRNA (15). While the structure and composition of ribosomes was thought to be largely invariant (16), several studies that examined the dynamics of RP composition and their relationship with ribosome function suggest the contrary (2, 7, 9, 11, 17–20). For example, ribosomal protein stoichiometry in yeast and mouse stem cells was shown to depend on the number of ribosomes bound per mRNA (9). Similarly, RPL38 was shown to be of paramount importance for the translation of Homeobox mRNAs in mice (2), while RPL10A-containing ribosomes were found to preferentially translate a subpool of mRNAs in mouse stem cells (7).

Eukaryotes contain many duplicated genes encoding RPs, which for a long time were thought to be functionally redundant (21, 22). However, a pioneering study in yeast revealed that depletion of RP paralogs does not lead to the same phenotypes, suggesting paralog-specific functions (3). In mammals, RP paralogs have been shown to be differentially expressed upon tumorigenesis (10) and some of them are exclusively expressed in restricted subsets of tissues (8, 10). For example, *Rpl3* is constitutively expressed in all tissues, whereas the expression of its paralog gene, *Rpl3l*, is restricted to heart and skeletal muscle tissues (23). Thus, while the existence of heterogeneous ribosomes in terms of RP composition is well documented (24, 25), the biological function of RP paralog genes with restricted tissue-specific expression, such as *Rpl3l*, is largely unknown.

The human *Rpl3l* gene was first identified in studies focusing on autosomal dominant polycystic kidney disease gene regions, in which the authors identified a gene sharing 77% identity with the mammalian *Rpl3* gene (26, 27). These studies found that *RPL3L* was exclusively expressed in heart and skeletal muscle, but no further indications were found as to its role in these tissues (27). Two decades later, it was found that RPL3L (uL3L) is downregulated upon hypertrophic stimuli, and it was proposed that RPL3L (uL3L) may function as a negative regulator of muscle growth (15). In support of this hypothesis, mutations in the human *Rpl3l* gene have been linked to several heart disorders, including atrial fibrillation (28, 29) and childhood-onset cardiomyopathy (30). However, why certain tissues express distinct RP proteins, and with what biological relevance, still remains unclear.

Here, we used a combination of wet lab and bioinformatics techniques to study the role of the RPL3(uL3)-RPL3L(uL3L) pair in protein translation in the mouse heart. We found that *Rpl3l* is expressed only postnatally, with its expression limited to specific cell types, in addition to being restricted to heart and muscle tissues. Specifically, we observed that in hearts, *Rpl3l* was exclusively expressed in cardiomyocytes, whereas *Rpl3* was mainly expressed in non-myocyte heart cell types. To explore the biological function of *Rpl3l*, we generated a viable homozygous *Rpl3l*^-/-^ mouse strain, and identified a rescue mechanism in which RPL3 becomes upregulated upon *Rpl3l* knockout, yielding RPL3-containing ribosomes instead of RPL3L-containing ribosomes that are typically found in mouse cardiomyocytes. Using both ribosome profiling (Ribo-seq) and ribosome pulldown coupled to nanopore sequencing (Nano-TRAP), we identified only a handful of differentially translated transcripts in *Rpl3l*^-/-^ hearts, suggesting that RPL3L does not play a major role in modulating translation efficiency or ribosome affinity towards preferentially translating a specific subset of transcripts. By contrast, when coupling ribosome pulldown to proteomics (Proteo-TRAP), we found that a very large amount of mitochondrial proteins were significantly enriched in RPL3-containing ribosome immunoprecipitates in cardiomyocytes. We then confirmed that RPL3, unlike RPL3L, is detected in the mitochondrial fraction of heart lysates, suggesting that the use of either of the two paralogs might be fine-tuning the ribosome-mitochondria interactions in cardiomyocytes. Moreover, we show that ATP levels, but not mitochondrial abundance, are significantly increased in *Rpl3l*^-/-^ cardiomyocytes.

Altogether, our work reveals that upon depletion of *Rpl3l*, RPL3 is upregulated, mimicking the RPL3/RPL3L interplay that occurs upon hypertrophic stimuli (15). This switch in RPL3/RPL3L expression patterns causes RPL3 to replace RPL3L in cardiac or skeletal muscle ribosomes, which in turn leads to higher proportion of ribosome-bound mitochondria and consequently increased ATP production.

## MATERIALS AND METHODS

### Generation of *Rpl3* and *Rpl3l* knockout mice

*Rpl3*^*+/-*^ *and Rpl3l*^*-/-*^ mice were produced by the Mouse Engineering Garvan/ABR (MEGA) Facility (Moss Vale and Sydney, Australia) using CRISPR/Cas9 gene targeting in C57BL/6J mouse embryos following established molecular and animal husbandry techniques (31). The single guide RNAs (sgRNAs) employed were based on the following target sequences (protospacer-associated motif = PAM italicised and underlined): *Rpl3* (Exon 6) TTCCAGGGGTAACCAGTCGTTGG and *Rpl3l* (Exon 5) GTGGGCCTTCTTCTGCCGAAAGG. To target *Rpl3 or Rpl3l*, 50 zygotes were electroporated in a solution containing the specific sgRNA (300 ng/μL) and Cas9 mRNA (600 ng/μL). Embryo preparation and electroporation was carried out according to the method of Qin et al (32), except that zygotes were incubated for 25 s (not 10 s) in acidic Tyrode’s solution and received three (not two) pulses. Microinjected or electroporated embryos were cultured overnight and those that underwent cleavage were introduced into pseudopregnant foster mothers. Pups were screened by PCR across the target site and Sanger sequencing of PCR products to detect modifications to the targeted gene. Founders carrying frameshift deletions were backcrossed to wild-type C57BL/6J mice and heterozygous progeny then inter-crossed. Viable homozygous mice were generated at normal Mendelian frequencies in the case of the *Rpl3l*^*Δ*^ lines whereas homozygous *Rpl3*^*Δ*^ mice all showed prenatal lethality.

### Validation of *Rpl3l*^-/-^ mice

The CRISPR/Cas9 deletion in the *Rpl3l* gene was validated: i) at the genomic level using Sanger sequencing, which confirmed that homozygous knockout mice had a 13 bp long deletion in the fifth exon (**Figure S1B**), and ii) at the mRNA level using sNuc-seq, which showed that reads mapping to *Rpl3l* had a 13 bp deletion in *Rpl3l*^-/-^ hearts (**Figure S1A)**. The lack of *Rpl3l* in *Rpl3l*^-/-^ mice was validated: i) at the mRNA level using RT-qPCR, which showed that *Rpl3l* was absent in the cytoplasm of striated muscles in *Rpl3l*^-/-^ mice, but present in WT mice (**Figure 2B)**; and ii) at the protein level using western blot (**Figure 2C**), immunofluorescence (**Figure 2D**) and proteomics (**Figure 5B** and **Table S9**), confirming the absence of RPL3L (uL3L) in *Rpl3l*^-/-^ mice.

**Figure 1.**
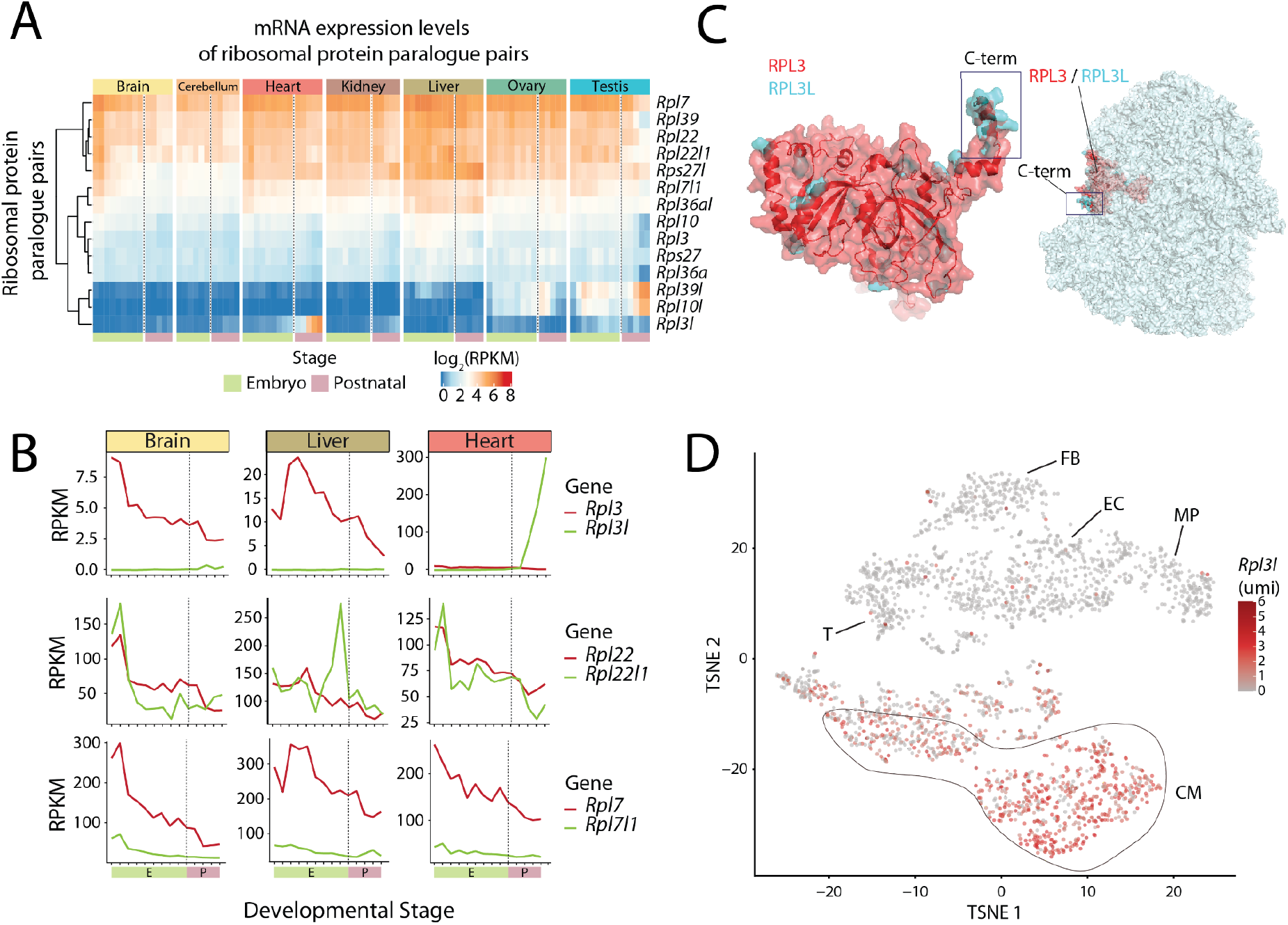
RPL3L (uL3L) is a vertebrate RP paralog with restricted tissue and developmental expression patterns, such as postnatal expression in mouse cardiomyocytes. **(A)** Heatmap of mRNA expression levels (log RPKM) of ribosomal proteins and their respective paralogs across embryonic (green: E10.5, E11.5, E12.5, E13.5, E14.5, E15.5, E16.5, E17.5, E18.5) and postnatal mice tissues (pink: P0, P3, P14, P28, P63). Processed data (RPKM) was obtained from Cardoso-Moreira et al. (42). See also Figure S7 for a heatmap containing all ribosomal proteins. **(B)** mRNA expression levels of ribosomal paralog pairs (RPKM) in 3 different tissues (brain, liver and heart) for *Rpl3*/*Rpl3l* (upper panel), *Rpl22*/*Rpl22l1* (middle panel) and *Rpl7* and *Rpl7l1* (bottom panel). The developmental stages, shown on the x-axis, have been colored depending on whether they correspond to embryonic (green) or postnatal (pink) stages. **(C)** Structural alignment of human RPL3 (red) and RPL3L (uL3L) (cyan), the C-terminus is highlighted (left) and location of RPL3 (uL3)/RPL3L (uL3L) within the ribosome (right). The C-terminus of RPL3 (uL3) and RPL3L (uL3L) is located at the surface of the ribosome, whereas the N-terminus of both proteins lies closer to the peptidyl transferase center (PTC). The ribosome structure has been obtained from the cryo-EM structure of the human 80S ribosome, corresponding to PDB code 6IP5 (61), which includes RPL3. The *H. sapiens* RPL3L (uL3L) structure was obtained from the ModBase (62) database and structurally superimposed to the RPL3 structure in the 80S ribosome. **(D)** T-distributed stochastic neighbor embedding (T-SNE) plot depicting *Rpl3l* expression across mouse heart cell types. Expression data has been extracted from publicly available single-cell RNAseq data from Ren et al. (63). Each dot represents a cell. Expression levels are shown as umi (unique molecular identifiers). Abbreviations: CM (cardiomyocytes), EC (endothelial cells), FB (fibroblasts), MP (macrophage), T (T cells).

**Figure 2.**
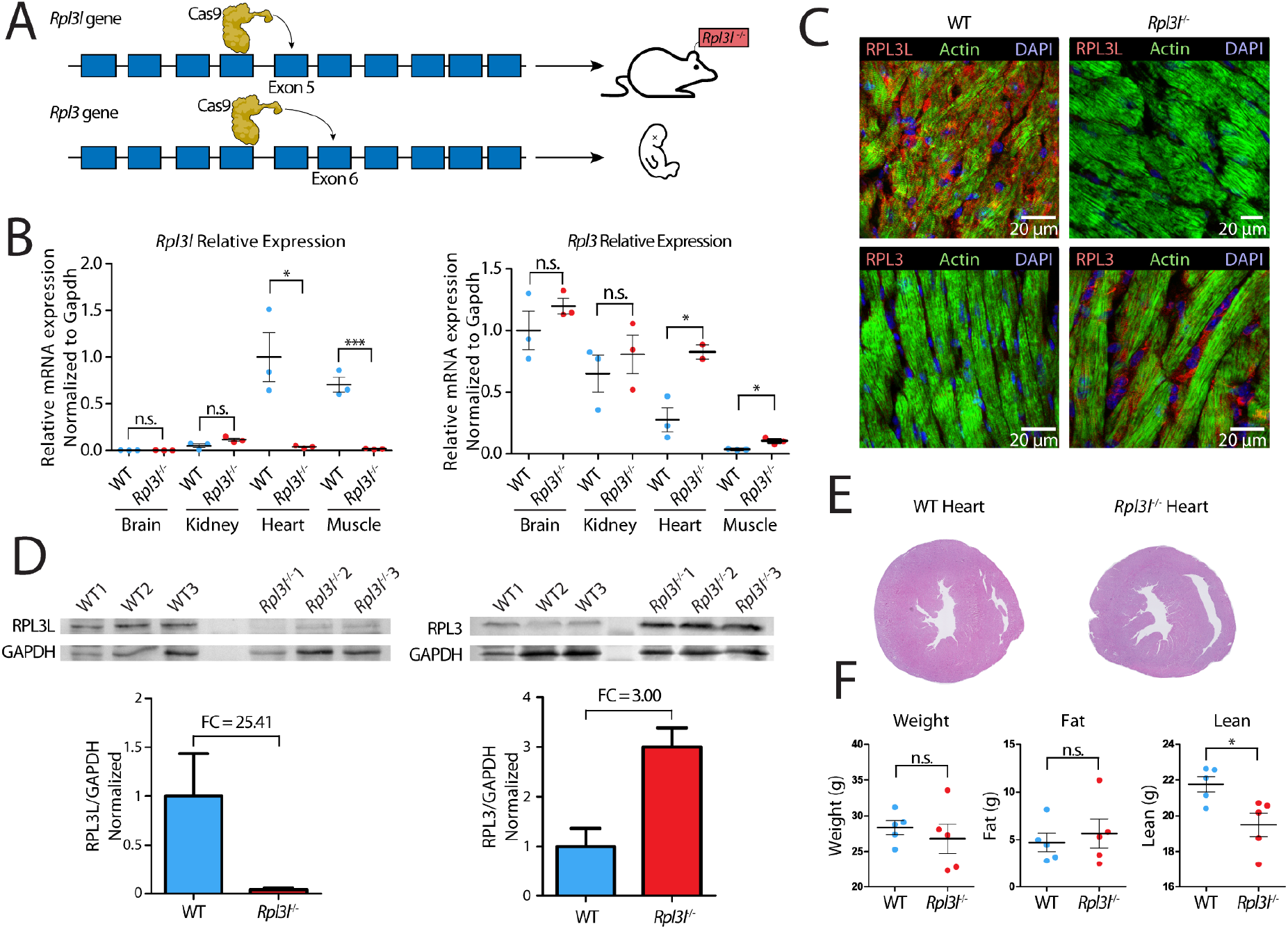
Phenotypic and molecular characterization of *Rpl3l*^-/-^ knockout mice. **(A)** Strategy for *Rpl3*^-/-^ and *Rpl3l*^-/-^ mice generation using the CRISPR-Cas9 system. *Rpl3l*^-/-^ mice were successfully generated by introducing a 13-bp deletion in exon 5. *Rpl3*^-/-^ mice have an embryonic lethal phenotype. See also Figure S1. **(B)** Relative expression levels of *Rpl3l* (left) and *Rpl3* (right) measured using RT-qPCR and normalized to *Gapdh. Rpl3* is ubiquitously expressed, while *Rpl3l* is heart- and muscle-specific. *Rpl3l*^-/-^ mice do not express *Rpl3l* in any of the tissues (n = 3). Statistical significance was assessed using the unpaired t-test (* for p < 0.05, ** for p < 0.01, *** for p < 0.001). **(C)** Immunofluorescence staining of RPL3 and RPL3L in both WT (left) and *Rpl3l*^-/-^ mice heart tissues (right). Nuclei have been stained with DAPI and are shown in blue, actin is depicted in green and RPL3L (uL3L) (top) and RPL3 (uL3) (bottom) in red. **(D)** Western blot analysis of RPL3L (uL3L) (left) and RPL3 (uL3) (right) in cardiomyocytes isolated from WT and *Rpl3l*^-/-^ hearts (n=3). In the bottom, barplots depicting the fold change of *Rpl3* and *Rpl3l* expression in cardiomyocytes is shown. RPL3L (uL3) and RPL3 (uL3) levels were normalised to GAPDH. See also Figure S4 for full blot images, and Figure S5 for western blot results using total heart samples from WT and *Rpl3l*^-/-^ mice. **(E)** Representative histological sections of WT and *Rpl3l*^-/-^ heart tissues stained with hematoxylin and eosin. A total of 10 mice were included in the histological analyses. See also Figure S9. **(F)** EchoMRI analyses of aged (55-week old) WT and *Rpl3l*^-/-^ mice, in which weight, fat and lean mass were measured for each animal (n = 5). Statistical significance was assessed using unpaired t-test (* for p < 0.05).

**Figure 3.**
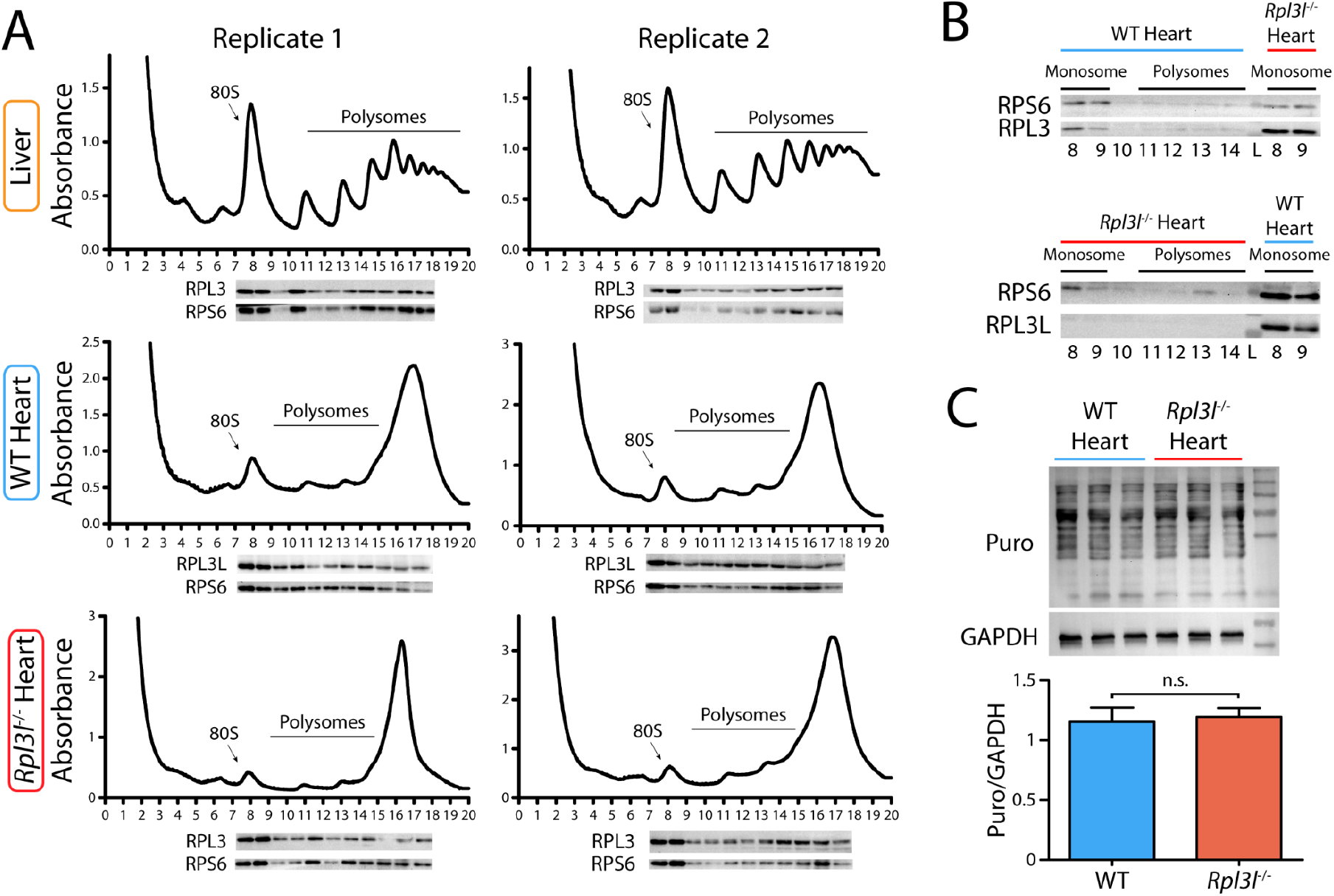
RPL3L and RPL3 are incorporated into translating ribosomes in WT and *Rpl3l*^-/-^ hearts, respectively. **(A)** Polysome profiles of liver (top), WT heart (middle) and *Rpl3l*^-/-^ heart (bottom) were performed in 10-50% sucrose gradients with corresponding western blot analyses, in two biological replicates. Two hearts were pooled together per replicate. Membranes were probed with anti-RPL3L (uL3L), anti-RPL3 (uL3) and anti-RPS6 (eS6) antibodies to show the incorporation of both paralogs in translating ribosomes. **(B)** Polysome profiles fractions of a WT heart (top) and a *Rpl3l*^-/-^ heart (bottom), analyzed with western blot and probed with anti-RPL3L (uL3L), anti-RPL3 (uL3) and anti-RPS6 (eS6). Fractions 8 and 9, corresponding to the monosome peak of the polysome profile, of a *Rpl3l*^-/-^ heart and WT heart, respectively, were used as positive controls. **(C)** Puromycin incorporation assay performed on WT and *Rpl3l*^-/-^ hearts in biological triplicates. GAPDH was used as loading control and for normalization in the densitometric analysis. Statistical significance was assessed using unpaired t-test. Puro stands for Puromycin.

**Figure 4.**
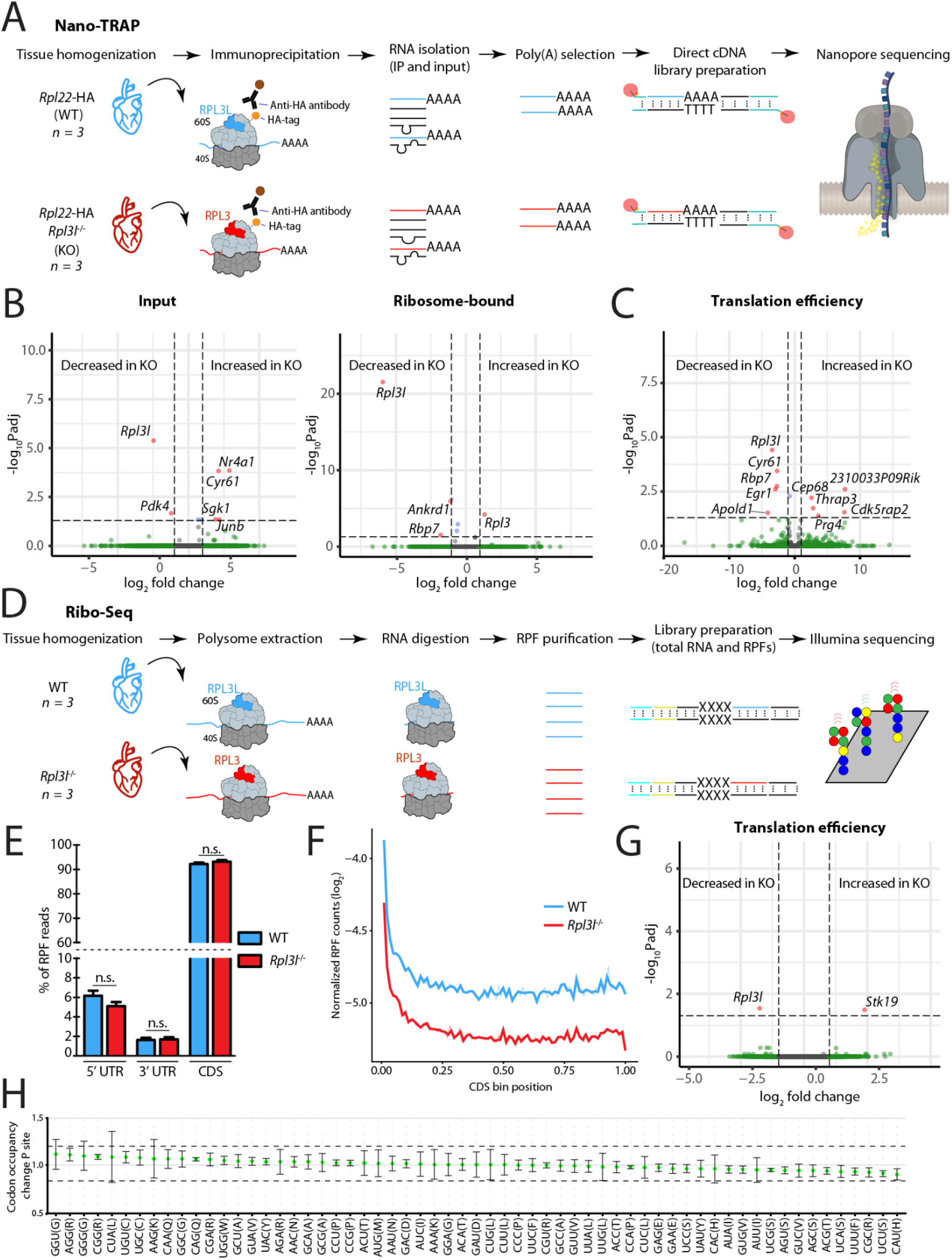
RPL3L usage does not lead to preferential translation or altered translation efficiency. **(A)** Schematic representation of the Nano-TRAP method. **(B)** Volcano plots representing differentially expressed input mRNA (left) and ribosome-bound mRNA (right) transcripts identified using Nano-TRAP, which correspond to those with fold change greater than 1 and FDR adjusted p-value lower than 0.05. Nano-TRAP results show minor differences in transcripts captured in RPL3L (uL3L)- and RPL3 (uL3)-bearing ribosomes. Each dot represents a gene, and they have been colored depending on: i) adjusted p-value < 0.05 and fold change > 1 (red), ii) only fold change > 1 (green) or only adjusted p-value < 0.05 (blue), iii) neither fold change > 1 nor adjusted p-value < 0.05 (gray). See also Table S4. **(C)** TE analysis using NanoTrap. Counts were normalised by the sum of counts for each sample. Every dot represents a gene. See also Table S4. **(D)** Schematic representation of the Ribo-Seq method. (**E**) Percentage of RPF reads mapping to 5’ UTR, 3’ UTR and CDS regions of identified genes. The values shown are means of three biological replicates. Error bars represent standard deviation. Statistical significance was assessed using the unpaired t-test (* for p < 0.05). (**F**) Metagene analysis of RPF reads from WT and *Rpl3l*^-/-^ hearts. (**G**) Analysis of differential translation efficiency (TE), calculated as the ratio between RPFs and mRNAs (see *Methods*), between WT and *Rpl3l*^-/-^ ribosomes. See also Table S6. **(H)** Codon occupancy change between WT and *Rpl3l*^-/-^ at the P-site.

**Figure 5.**
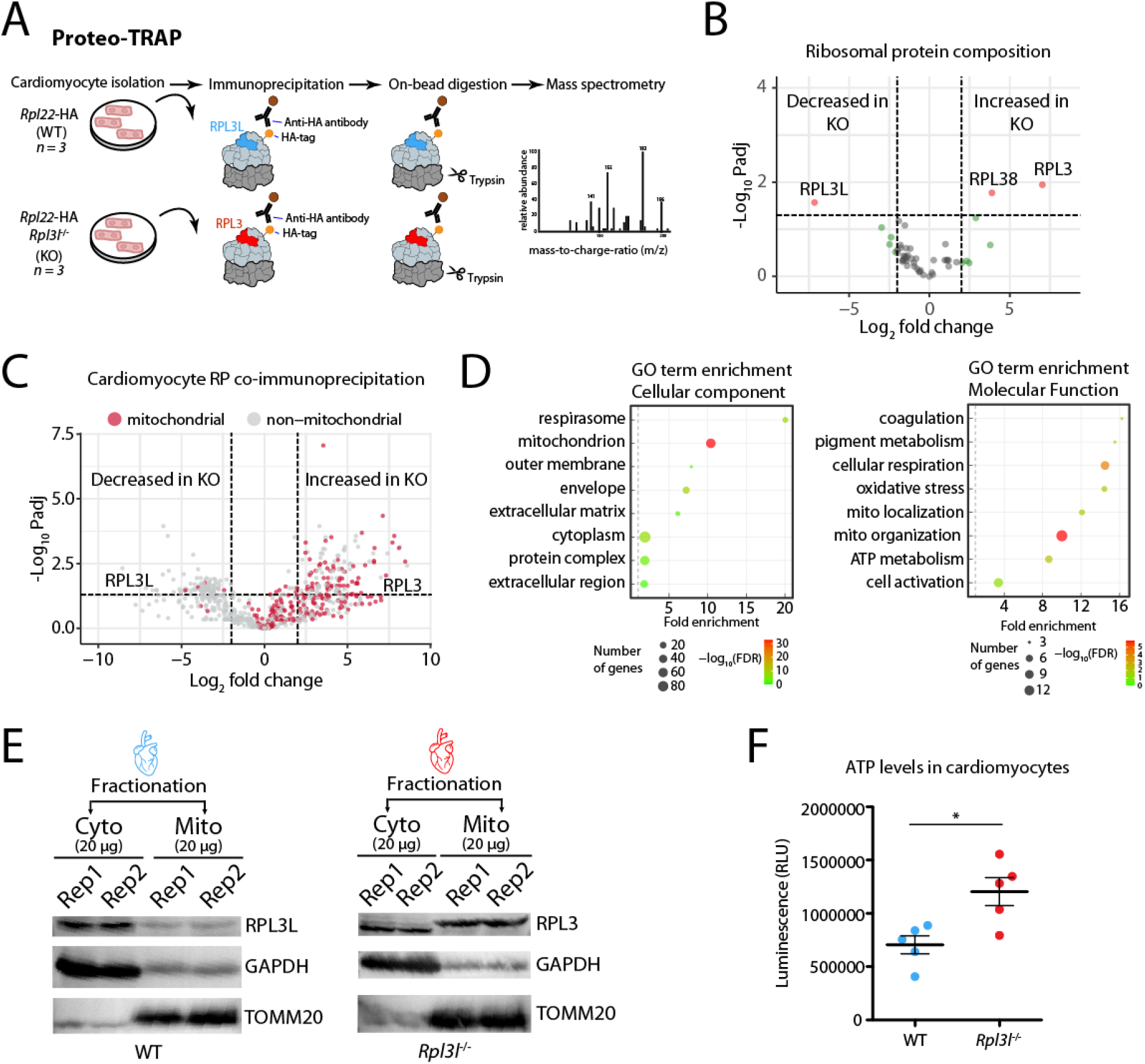
RPL3 (uL3)-containing ribosomes establish physical contact with mitochondria. **(A)** Schematic representation of the Proteo-TRAP method. **(B)** Analysis of differential ribosome composition in cardiomyocytes from WT and *Rpl3l*^-/-^ mice. See also Table S10. **(C)** Volcano plot showing mitochondrial (red) and non-mitochondrial (gray) proteins co-precipitating with ribosomes in WT and *Rpl3l*^-/-^ cardiomyocytes. See also Table S11 and Figure S17. **(D)** GO term enrichment plots showing top hits for cellular components (left) and molecular function (right). **(E)** Western blot analysis of cytosolic and mitochondrial fractions of WT (left) and *Rpl3l*^-/-^ (right) hearts. GAPDH and TOMM20 were used as cytosolic and mitochondrial fraction markers, respectively. See also Figure S17 for full membrane images with marker sizes. **(F)** Luminometric measurement of ATP levels in WT and *Rpl3l*^-/-^ cardiomyocytes. Statistical significance was assessed using unpaired t-test (* for p < 0.05).

We should note that our results show that our sNuc-seq results showed that both *Rpl3* and *Rpl3l* are being transcribed both in WT and *Rpl3l*^-/-^ mice hearts, as they can be detected at the mRNA level in the nucleus of both WT and *Rpl3l*^-/-^ hearts (**Figure S2A**), in agreement with the predicted chromatin states of these genes in heart samples (**Figure S3**). Altogether, our results suggest that the 13 bp deletion in *Rpl3l*^-/-^ mice leads to degradation of the *Rpl3l* mRNA at the cytosolic level (**Figure 2B**), and consequently lack of RPL3L (uL3L) protein (**Figure 2C,D**).

### Generation of *Rpl3l*^-/-^-RiboTag-E2a-Cre mice

RiboTag mice were purchased from The Jackson Laboratory (Stock No: 011029). This mouse line, generated by the McKnight lab (33), carries a floxed wild-type C-terminal exon in the *Rpl22* gene, followed by a mutant exon that has a triple hemagglutinin (HA) epitope inserted in front of the stop codon, leading to the generation of HA-tagged ribosomes when crossed with a Cre-expressing mouse strain. The RiboTag mice were first crossed with the E2a-Cre mouse line, also purchased from The Jackson Laboratory (Stock No: 003724) and bred to homozygous genotype, ensuring the excision of the wild-type C-terminal exon and the expression of HA-tagged *Rpl22* in all tissues. These mice were then crossed with the *Rpl3l*^−**/-**^ strain and bred to obtain a homozygous genotype.

### Mice tissue collection

All experiments were performed with male mice aged between 8 and 10 weeks, unless stated otherwise. All mice were euthanized using CO_2_, except for the Ribo-Seq, Nano-TRAP and single nuclei RNA-Seq experiments, for which mice were euthanized using cervical dislocation, and tissues were snap frozen in liquid nitrogen.

### Total RNA extraction from mice tissues

Tissues were homogenised in TRIzol (Life Technologies, 15596018) using the Polytron PT 1200 E hand homogenizer in pulses of 10 seconds at maximum speed until thoroughly homogenised. Total RNA was extracted using ethanol precipitation, and purity and concentration were measured using NanoDrop spectrophotometer.

### Body composition measurements using magnetic resonance imaging (MRI)

The body composition of 8 week old male and 55 week old female *Rpl3l*^*-/-*^ and *Rpl3l*^+/+^ mice was analysed using the EchoMRI^™^-100H Body Composition Analyser without anaesthetising. The EchoMRI^™^-100H delivers precise body composition measurements of fat, lean, free water and total water masses in live animals. Statistical significance of the results was assessed using unpaired t-test.

### Histopathological analyses of *Rpl3*^-/-^ knockout and control mice

A complete necropsy of 8 week old male *Rpl3l*^*-/-*^ and *Rpl3l*^+/+^ mice (n = 5) was performed to check for possible macroscopic findings. Mice were weighed before euthanasia with CO_2_. The heart and brain were weighed in order to obtain the body weight to organ and brain-organ ratios. The right quadriceps, right gastrocnemius and apex of the heart were embedded in optimal cutting temperature compound (OCT) and were frozen straight away. The left quadriceps, left gastrocnemius and the whole heart (excluding the apex) were collected and fixed in buffered 4% paraformaldehyde overnight and transferred to PBS. The tissues were then trimmed, paraffin embedded, sectioned (3-4 μm tissue sections) and stained with Haematoxylin/Eosin (H&E). All the hearts were trimmed transversally at the same level in order to obtain a biventricular short axis section (biventricular, transversal). The proximal area of the gastrocnemius and quadriceps were trimmed transversally, and the remaining tissue was sectioned longitudinally. At least three cross sections of each heart were evaluated in a blinded manner in order to detect histopathological changes of the myocardium such as disarray, myocardial degeneration, hypertrophy or atrophy and other possible alteration such as inflammation or fibrosis. Furthermore, quantitative analysis of the left ventricle, right ventricle and myocardial fibres was performed using the NDP.view 2 2.8.24 software (Hamamatsu). The measurements of the left and right ventricular free walls were performed as previously reported (34), with a single line from the endocardium to the pericardium oriented as perpendicular as possible to both surfaces, always at areas without papillary muscle. Five measurements were performed preferably in the same cross section, except in cases that it was not possible. The muscle samples were evaluated in a blinded manner in order to detect histopathological changes such as sarcoplasmic enlargement or thinning, vacuolization or nuclear centralisation (signs of degeneration or atrophy) and other possible alterations such as inflammation or fibrosis. A description of the main histological features present was done using the following semi-quantitative grading system: 1; minimal, 2; mild, 3; moderate, 4; marked, 5; severe.

### Echocardiography analyses of *Rpl3*^-/-^ knockout and control mice

Anaesthesia was induced in mice using 5% isoflurane then weighed and transferred to a warming pad (40 °C) with built in ECG for heart rate (HR) determination. The pad is integrated into the Vevo3100 (Fujifilm SonoSite, Inc., WA, USA) ultrasound system. An MX400 probe was used for all assessments. The chest was shaved, washed and ultrasound gel applied. Isoflurane was adjusted to keep HR as close to 500 BPM as possible while inhibiting the righting reflex. Systolic function was assessed from single high resolution ECG gated B-mode cine-loops (EKV). Endocardial and epicardial volumes were determined using the bullet formula Vol = 5/6 x L x A, where A is the area determined by planimetry at the mid-papillary level and L is long axis length from the junction of aortic leaflets to the apical dimple. stroke volume (SV) was calculated as the difference between endocardial volumes at end diastole (ED) and end systole (ES), EDV and ESV respectively. Ejection fraction (EF) was calculated as SV / EDV. ED was defined as the beginning of the ECG R-wave upstroke and ES as the closure of the aortic valve. Diastolic function was assessed using pulse-wave and tissue Doppler from an apical 4-chamber view of the mitral valve. Left atrial dimension was determined as the diameter along a line aligned to the inferior edge of the aortic annulus in a parasternal long axis B-mode image at end-systole. All recorded metrics, for both WT and *Rpl3l*^-/-^ mice, can be found in **Table S2**.

### Micromanometry analyses of *Rpl3*^-/-^ knockout and control mice

Micromanometry was performed 24-48 hours after echocardiography in a randomised blinded way. Following anaesthetic induction under 4-5% isoflurane in medical oxygen, mice were laid and secured supine on a pre-warmed mat with a nose cone delivering 1-2% isoflurane. To monitor anaesthesia, pedal reflex and respiration rate were observed. With the head extended, the neck was shaved and washed, then a 1-1.5cm longitudinal incision to the right of the anterior midline was made. Using blunt dissection the sternohyoid and sternomastoid muscles were separated to expose the internal right carotid artery posterior and lateral to the trachea. The vessel was isolated with three 4-0 Silk (J&J Inc. NJ, USA) ligatures. The distal was ligated while the proximal was looped under the artery, these two were then tensioned against each other while the middle was run loosely twice around the artery. The pre-warmed and calibrated 1.2F Scisence (Transonic Scisense Inc.,. NY, USA) catheter was dipped in lubricating gel and introduced into the artery via a puncture made using a 32G needle at the distal end, then the middle ligature was tensioned to secure the catheter. With the proximal ligature released the catheter was progressed into the aortic arch. A characteristic sawtooth signal was observed in the acquisition software display (Acqknowledge Ver 3.8.2. BioPac Systems Inc.,. CA, USA) and was recorded for several minutes. During this time, the anaesthesia was lightened to allow heart rate to increase to ∼500BPM, before the catheter tip was advanced into the LV and recorded for several minutes before removal. Analysis was performed using the same software, and results can be found in **Table S2**.

### RT-qPCR

Total RNA isolated from mouse tissues was used as starting material for qPCR reactions. Briefly, total RNA was first DNase-treated with TURBO™ DNase (Life Technologies, AM2239) for 30 minutes at 37 °C to remove possible DNA contamination. Total RNA was then re-extracted using acid phenol-chloroform (Life Technologies, AM9720) and quantified using NanoDrop spectrophotometer. 100 ng of total RNA was mixed with Oligo(dT) primers, random hexamers and 10mM dNTP mix (Invitrogen, 18080051). The reaction mixture was heated to 65 °C in a thermocycler, after which it was quickly chilled on ice. RNase inhibitors (Invitrogen™ RNaseOUT™), 0.1M DTT and the First-strand buffer (Invitrogen, 18080051) were added to the reaction mixture which was incubated at 42 °C for 2 minutes. SuperScript™ II reverse transcriptase (Life Technologies, 18064-014) was finally added, and the reaction mixture was incubated at 42 °C for 50 minutes, followed by a 15 minutes-long inactivation at 70 °C. The synthesized cDNA was diluted 1:6 and used for the qPCR reaction. The Power SYBR Green Master Mix (Life Technologies, A25741) was used as the fluorescent dye, and the primers were used as follows: *Rpl3_fw:* GGAAAGTGAAGAGCTTCCCTAAG, *Rpl3_rev*: CTGTCAACTTCCCGGACGA, *Rpl3l_fw*: GAAGGGCCGGGGTGTTAAAG, *Rpl3l_rev*: AGCTCTGTACGGTGGTGGTAA, *GAPDH_fw*: AGCCTCGTCCCGTAGACAAA, *GAPDH_rev:* AATCTCCACTTTGCCACTGC. All oligonucleotide sequences used in qPCR experiments can be found in **Table S15**. The LightCycler® 480 Instrument II was used for the qPCR reaction with the following programme: 90 °C, 2 min; 40 cycles of 90 °C, 5 s; 60 °C, 10 s; 72 °C, 20 s.

### Quantification of mitochondrial genome copy number

Mitochondrial DNA (mtDNA) quantification was performed according to Quiros et al. (35). Briefly, mice were euthanized and hearts were collected and cut into small pieces. 10-30 mg of tissue was then transferred to an ice cold 1.5 mL eppendorf tube and 600 µL of lysis buffer (100 mM NaCl, 10 mM EDTA, 0.5% SDS, 20 mM Tris-HCl pH 7.4) was added, followed by 0.2 mg/mL of proteinase K. The samples were incubated overnight at 55 °C. The next day, 100 µg/mL of RNase A (Qiagen, 158922) was added and the samples were incubated for 30 minutes at 37 °C. 600 µL of ultrapure phenol: chloroform: isoamyl alcohol (25:24:1, v/v) was added and mixed well. The samples were centrifuged at 12,000 x g for 5 minutes, and the aqueous phase was transferred to a new tube. 250 µL of 7.5 M ammonium acetate and 600 µL of isopropanol (0.7 v/v) were mixed well. The samples were then centrifuged at 15,000 x g for 10 minutes at 4 °C. The supernatant was discarded, and the pellet was washed with 500 µL of 70% ethanol. The pellet was air-dried and resuspended in TE buffer (10 mM Tris-HCl pH 7.4 and 1 mM EDTA). Concentration was measured using NanoDrop and the samples were diluted to 10 ng DNA/µL. qPCR was done as previously described. The primers were used as follows: *mt16S_fw*: CCGCAAGGGAAAGATGAAAGAC, *mt16S_rev*: TCGTTTGGTTTCGGGGTTTC, *Nd1_fw*: CTAGCAGAAACAAACCGGGC, *Nd1_rev*: CCGGCTGCGTATTCTACGTT, *Hk2_fw*: GCCAGCCTCTCCTGATTTTAGTGT, *Hk2_rev*: GGGAACACAAAAGACCTCTTCTGG.

### Polysome profiling

Whole hearts were cut into small pieces and homogenized in pre-chilled 2 mL eppendorf tubes using a bead beater in ice-cold polysome extraction buffer (PEB), which contained 20 mM Tris-HCl pH 7.4, 200 mM KCl, 10 mM MgCl2, 1 mM DTT, 0.1 mg/mL of cycloheximide, 10 U/mL of DNaseI (NEB, M0303S), 1x cOmplete^™^ protease inhibitor cocktail (Roche, 11873580001) and 100 U/mL of RNAse inhibitors (RNaseOUT, Invitrogen, no. 18080051) using the following program: 2 cycles of 15 seconds, 1 minute incubation on ice and 1 cycle of 30 seconds. The homogenates were incubated for 5 minutes on ice and then centrifuged at 20.000 x g for 12 minutes at 4 °C. Triton X-100 was added to the final concentration of 1%, and the homogenates were incubated for 30 minutes on an end-over-end rotator at 4 °C. The samples were then centrifuged in a benchtop centrifuge at maximum speed for 12 minutes and the supernatant was transferred to a new tube. Linear sucrose gradients of 10–50% were prepared using the Gradient Station (BioComp). Briefly, SW41 centrifugation tubes (Beckman Coulter, Ultra-ClearTM 344059) were filled with Gradient Solution 1 (GS1), which consisted of 20 mM Tris-HCl pH 8, 200 mM KCl, 10 mM MgCl2, 0.2 mg/mL of cycloheximide and 10% wt/vol RNAse-free sucrose. Solutions GS1 and Gradient Solution 2 (GS2) were prepared with RNase/DNase-free UltraPure water and filtered with a 0.22-µm filter. The tube was then filled with 6.3 mL of GS2 layered at the bottom of the tube, which consisted of 20 mM Tris-HCl pH 8, 200 mM KCl, 10 mM MgCl2, 0.2 mg/mL of cycloheximide and 50% wt/vol RNAse-free sucrose. The linear gradient was formed using the tilted methodology, with the Gradient Station Maker (BioComp). Once the gradients were formed, 700 µl of each lysate was carefully loaded on top of the gradients, and tubes were balanced in pairs, placed into pre-chilled SW41Ti buckets and centrifuged at 4 °C for 150 minutes at 35,000 r.p.m. Gradients were then immediately fractionated using the Gradient Station, and 20 × 500 µL fractions were collected in 1.5 mL Eppendorf tubes while absorbance was monitored at 260 nm continuously. Total proteins were extracted from individual fractions by adding trichloroacetic acid (TCA) to the final concentration of 10% v/v, followed by centrifugation at 4 °C at maximum speed for 10 minutes. The supernatant was discarded, the protein pellets were dissolved in NuPAGE™ LDS Sample Buffer supplemented with NuPAGE™ Sample Reducing Agent and later used for subsequent western blot analysis.

### Puromycin incorporation assay

The mice were euthanized by cervical dislocation, the aortae were clamped using hemostatic forceps and the hearts were quickly excised and placed onto a petri dish. 1.2 µmol of puromycin (Sigma-Aldrich, P8833) dissolved in PBS was injected per single heart though the left ventricle, and the buffer was reinjected for 20 minutes. The hearts were snap frozen and later used for western blot analysis. The anti-puromycin antibody (DSHB, PMY-2A4) was used at a concentration of 0.4 µg/mL.

### Isolation of the mitochondrial fraction

The mice were euthanized by cervical dislocation and the hearts were quickly excised and placed into ice cold BIOPS (10 mM Ca-EGTA, 0.1 µM free calcium, 20 mM imidazole, 20 mM taurine, 50 mM K-MES, 0.5 mM DTT, 6.56 mM MgCl_2_, 5.77 mM ATP, 15 mM phosphocreatine, pH 7.1). Blood clots were carefully removed and the hearts were cut into small pieces using cooled scissors. The tissue was transferred into a 2 mL Dounce homogenizer, and 0.5 mL of isolation buffer (225 mM mannitol, 75 mM sucrose, 1 mM EGTA) was added. The tissue was homogenized at medium speed in 6-8 strokes. The homogenate was transferred to a 1.5 mL tube and centrifuged at 800 g for 10 minutes at 4 °C. The supernatant was transferred to a new 1.5 mL falcon tube, and centrifuged at 10,000 g for 10 minutes at 4 °C. The supernatant was saved for subsequent western blot analysis of the cytoplasmic fraction. The mitochondrial pellet was carefully resuspended in 1 mL of isolation buffer and centrifuged at 10,000 g for 10 minutes at 4 °C. The supernatant was discarded and the mitochondrial pellet was lysed in 100 μL of ice-cold RIPA buffer. The lysate was used for subsequent western blot analysis.

### Protein extraction and Western Blot

Frozen tissues were chopped into smaller pieces and homogenized in ice-cold RIPA buffer using the Polytron PT 1200 E hand homogenizer in pulses of 10 seconds at maximum speed until thoroughly homogenized. The homogenates were then agitated on an end-over-end shaker at 4 °C for 2 hours. Finally, the homogenates were centrifuged for 20 minutes at 12,000 rpm at 4 °C in a microcentrifuge. The supernatant was placed in a new tube and kept on ice, and the pellet discarded. Protein concentration was measured using the Bradford assay, and ∼15 μg of total protein was mixed with the NuPAGE™ LDS Sample Buffer supplemented with NuPAGE™ Sample Reducing Agent, incubated at 70 °C for 10 minutes and loaded onto a 12% polyacrylamide gel. The electrophoresis was run at 80V until the samples reached the resolving gel, when the voltage was increased to 120V. The transfer was performed onto a PVDF membrane in a Bolt™ Mini Gel Tank, at a constant voltage of 20V for 50 minutes. The membrane was then washed three times for 5 minutes in TBST, after which it was blocked for 1 hour in 3% BSA in TBST (blocking buffer). The incubation with primary antibodies (RPL3 Rabbit Polyclonal Antibody (Proteintech, 11005-1-AP); RPL3L (uL3L) Rabbit Polyclonal Antibody (custom-made, kindly provided by Prof. John McCarthy), RPL7 Rabbit Polyclonal Antibody (Abcam, ab72550) and TOMM20 Rabbit Polyclonal Antibody (Abcam, ab78547)) was performed in blocking buffer overnight at 4 °C at a 1:1000 dilution. The next day, the membrane was washed three times for 5 minutes, and incubated in the secondary HRP-coupled antibody (Abcam, ab6721) at a 1:10,000 dilution in blocking buffer for 1 hour at room temperature. The membrane was washed three times and imaged using the SuperSignal™ West Pico PLUS Chemiluminescent Substrate. The membrane was then reprobed with an anti-GAPDH antibody (1:1000 in blocking buffer) for one hour, and reimaged the same way. Full images of all western blots shown in this work can be found in **Figures S4, S5** and **S6**. We should note that commercially available anti-RPL3L (uL3L) antibodies were also tested (Abcam, ab200646; Biorbyt, orb234958; Sigma-Aldrich, HPA049136) but none of them gave specific bands that were present in WT and absent in *Rpl3l*^-/-^ samples. The custom-made anti-RPL3L (uL3L) antibody used here also has some non-specific bands, but those bands are also present in tissues in which *Rpl3l* is not expressed, such as the liver (**Figure S4D**).

### Immunofluorescence assays

Freshly collected heart and muscle tissues were quickly frozen with OCT (Optimal cutting temperature compound) and cut in 12 µm thickness slices using a cryotome. The slices were circled with a hydrophobic marker and incubated with 4% formaldehyde in PBS for 10 minutes at room temperature. The slides were then washed in PBS in staining jars three times for 5 minutes. To ensure the permeabilization of the cells, the samples were incubated with 0.5% Triton-X-100 in PBS for 30 minutes at 4 °C. Three washes with PBS were repeated, and the tissues were blocked with 3% BSA in TBS-T for 30 minutes at room temperature. The slides were washed with TBS-T three times for 5 minutes, and incubated with the primary antibodies (RPL3 Rabbit Polyclonal Antibody (Proteintech, 11005-1-AP); RPL3L Rabbit Polyclonal Antibody (custom-made in the lab of Prof. John McCarthy); anti-Atp5a Mouse Monoclonal Antibody (Abcam, ab14748) in blocking solution at a dilution of 1:200 (5 μg/mL in the case of anti-Atp5a) overnight at 4 °C. The next day, the samples were washed with TBS-T three times for 5 minutes, and incubated in the secondary antibody mixture (Anti-Rabbit Secondary Antibody Alexa Fluor 555 (Thermo Fisher Scientific, A-21429) 1:400, Phalloidin-iFluor 488 (Abcam, ab176753), 1:1000 and Hoescht 33342 (Thermo Fisher Scientific, H3570) 1:1000 in blocking solution) for one hour at room temperature. The slides were then washed with TBS-T three times for 5 minutes, air-dried for a few minutes and mounted with coverslips. The immunofluorescence images were made using the Leica TCS SPE confocal microscope.

### Luminometry assays for ATP quantification

Cardiomyocytes isolated from WT and *Rpl3l*^-/-^ hearts were diluted to same concentrations and the cell suspensions were mixed with the same volume of CellTiter-Glo (Promega, G7570), as instructed by the manufacturer. The reaction mixtures were pipetted into wells of an opaque 96-well plate and mixed for 2 minutes on an orbital shaker. The plate was then incubated for 10 minutes at room temperature and the luminescence was read using the Berthold LB 960 Microplate Luminometer.

### Phylogenetic analysis

To build a phylogenetic tree of the *Rpl3*/*Rpl3l* family, full proteomes of representative species (36) were downloaded from Uniprot. A Hidden Markov Model (HMM) profile of RPL3 (PF00297) was downloaded from Pfam (37), and was used to query the selected proteomes using the hmmsearch function from the HMMER package (version 3.3) (38). Hits were then aligned to the HMM using the hmmalign function from HMMER, and a phylogenetic tree was built using the maximum likelihood method of IQ-TREE (version 1.6.11) (39, 40). The final tree was visualized using FigTree (version 1.4.4) (41). Code to reproduce the phylogenetic analyses is available in GitHub: https://github.com/novoalab/RPL3L/tree/main/Phylogeny.

### Analysis of ribosomal protein expression patterns across mouse tissues and developmental stages

Previously published RNA-seq data (42) was used for the expression analysis. The RPKM values for all tissues across the time-series were downloaded from the ArrayExpress EBI repository (https://www.ebi.ac.uk/arrayexpress/files/E-MTAB-6798/E-MTAB-6798.processed.1.zip). RPKM values corresponding to ribosomal proteins were extracted from the full list, and the median expression values were calculated from biological replicates. The *ComplexHeatmap* R package was used to construct the heatmaps shown in **Figure 1** and **Figure S7**, and the code to reproduce the heatmap figures can be found in GitHub: https://github.com/novoalab/RPL3L/tree/main/ComplexHeatmap.

### Ribosome pulldown using anti-HA antibodies (for Nano-TRAP)

The HA-pulldown on *Rpl3l*^-/-^-RiboTag-E2a-Cre and *Rpl3l*^+/+^-RiboTag-E2a-Cre mice hearts was done using a modified approach devised by Sanz et al. (43). The mice were euthanized using cervical dislocation, and the hearts were excised, removing the aorta and the atria. The hearts were placed in ice-cold PBS supplemented with 100 µg/mL cycloheximide (CHX) and the blood was removed by gently squeezing with forceps. While on ice, the hearts were chopped to smaller pieces and added to pre-chilled 2 mL tubes containing ∼100 μL of acid-washed glass beads (425 - 600 µm) and 1 mL of homogenization buffer (50 mM Tris-HCl, pH 7.5; 100 mM KCl; 12 mM MgCl2; 1% Nonidet P-40 substitute; 1 mM DTT; 200 U/mL RNasin; 1 mg/mL heparin; 100 μg/mL cycloheximide; 1× protease inhibitor mixture). The hearts were then homogenized using a Mini Beadbeater (BioSpec Products) in 2 cycles of 60 seconds and 1 cycle of 15 seconds, allowing the samples to cool down on ice for 1 minute between the cycles. The lysates were cleared by centrifuging for 10 minutes at 10,000 x g at 4 °C. The supernatants were transferred to pre-chilled DNA LoBind tubes, and small aliquots of 40-80 µL were transferred to separate tubes and kept at -80 °C for subsequent input analysis. 4 µL of anti-HA antibody (BioLegend, 901513) was added to the remaining cleared lysate and incubated for 4 hours at 4 °C on an end-over-end rotator. Pierce™ Protein A/G Magnetic Beads were resuspended by gentle vortexing and transferred into DNA LoBind tubes. The tubes were placed on a magnetic stand and the storage buffer was discarded. 400 µL of the homogenization buffer was added and the tubes were incubated for 5 minutes on an end-over-end rotator. The beads were collected with the magnetic stand and the buffer was discarded. The cleared lysate and with the antibody was added to the beads and incubated overnight at 4 °C in an end-over-end rotator. On the next day, the high-salt buffer was prepared freshly (50 mM Tris-HCl, pH 7.5; 300 mM KCl; 12 mM MgCl2; 1% Nonidet P-40 substitute; 0.5 mM DTT; 100 μg/mL cycloheximide). The samples were placed into the magnetic stand and the supernatant removed. 800 µL of high-salt buffer was added to tubes to remove nonspecific binding from the immunoprecipitates and the tubes were washed for 5 minutes on an end-over-end rotator at 4 °C. The tubes were then placed into the magnetic stand and the supernatant removed. The high-salt washes were repeated twice more, for a total of three washes. In the last washing step, the beads were transferred to a clean tube, and all high-salt buffer was carefully removed. 350 µL RLT buffer from the Qiagen RNeasy extraction kit supplemented with β-mercaptoethanol was added to the tubes, which were then vortexed for 30 seconds at room temperature. The tubes were placed into the magnetic stand, and total RNA was extracted from the immunoprecipitates following Qiagen’s RNeasy extraction kit directions. Total RNA was quantified using Nanodrop and Qubit, and the integrity was assessed using the Tapestation.

### Purification of cardiomyocytes from total hearts

The purification of cardiomyocytes from mouse hearts was done according to a protocol published by Acker-Johnson et al. (44). Briefly, the mice were anaesthetized with isoflurane and the chest was opened to expose the heart. The descending aorta was cut and the heart was perfused through the right ventricle with 7 mL of EDTA buffer (130 mM NaCl,5 mM KCl, 0.5 mM NaH_2_PO_4_, 10 mM HEPES, 10 mM glucose, 10 mM BDM, 10 mM taurine, 5 mM EDTA, pH adjusted to 7.8 with NaOH and sterile filtered). Ascending aorta was clamped using hemostatic forceps, and the heart was excised and submerged in fresh EDTA buffer in a 60 mm dish. The heart was then perfused through the left ventricle with 10 mL EDTA buffer, 3 mL perfusion buffer (130 mM NaCl,5 mM KCl, 0.5 mM NaH_2_PO_4_, 10 mM HEPES, 10 mM glucose, 10 mM BDM, 10 mM taurine, 1 mM MgCl_2_, pH adjusted to 7.8 with NaOH and sterile filtered) and 30-50 mL of collagenase buffer (0.5 mg/mL collagenase 2, 0.5 mg/mL collagenase 4, 0.05 mg/mL protease XIV in perfusion buffer, heated to 37 °C). Digested hearts were pulled apart into 1 mm pieces using forceps and gently triturated using a P1000 pipette. Digestion was stopped by adding stop buffer (perfusion buffer with 5% FBS), and the cell suspension was filtered through a 100 µm strainer. The cells were submitted to two rounds of gravitational sedimentation of 15-20 minutes each, and then used for subsequent analyses.

### Preparation of cardiomyocyte ribosomes (for Mass Spectrometry analysis)

After isolating cardiomyocytes from *Rpl3l*^-/-^-RiboTag-E2a-Cre and Rpl3l^+/+^-RiboTag-E2a-Cre mice, ribosomes were pulled down using anti-HA antibodies and magnetic beads, as explained above. Ribosomes bound to magnetic beads were washed three times using the high salt buffer, and then another six times with 200 mM ammonium bicarbonate (ABC, #09830-500G, Sigma, MI, US) to wash away the detergent. The beads were then resuspended in 6 M urea (#17-1319-01, GE Healthcare, UK), reduced with 30 nmols dithiothreitol (#D9163-25G, Sigma, MI, US) (37 °C, 60 min with shaking), alkylated in the dark with 60 nmols iodoacetamide (#D9163-25G, Sigma, MI, US), (25 °C, 30 min) and then diluted to 1M urea with 200 mM ABC for trypsin digestion (Sequence-grade, #V5111, Promega, WI, US) (1 µg, at 37 °C, overnight with shaking). All buffer were prepared in 200 mM ABC.The beads were separated from the supernatant on a magnet, and the supernatant was acidified with 100% formic acid (#1.00264.0100, Merck, DK). C18 stage tips (UltraMicroSpin Column, #SUM SS18V, The Nest Group, Inc., MA, US) were then conditioned by adding methanol (#14262, Sigma, MI, US) and centrifuged at 100 g for 5 min. They were then equilibrated by two additions of 5% formic acid and centrifuged as in the previous step. Acidified samples were then loaded onto the columns and centrifuged at 100 g for 10 min. The samples were reapplied and the centrifugation step repeated. Three washing steps were performed with 5% formic acid, and the peptides were eluted using 50% acetonitrile (#34967, Sigma, MI, US) 5% formic acid. The eluate was vacuum dried and used for subsequent analyses.

### Digestion and analysis of cardiomyocyte protein samples using Mass Spectrometry

Samples were analysed using a Orbitrap Eclipse mass spectrometer (Thermo Fisher Scientific, San Jose, CA, USA) coupled to an EASY-nLC 1200 (Thermo Fisher Scientific (Proxeon), Odense, Denmark). Peptides were loaded directly onto the analytical column and were separated by reversed-phase chromatography using a 50-cm column with an inner diameter of 75 μm, packed with 2 μm C18 particles spectrometer (Thermo Scientific, San Jose, CA, USA). Chromatographic gradients started at 95% buffer A and 5% buffer B with a flow rate of 300 nL/minutes for 5 minutes and gradually increased to 25% buffer B and 75% A in 79 minutes and then to 40% buffer B and 60% A in 11 min. After each analysis, the column was washed for 10 minutes with 10% buffer A and 90% buffer B. Buffer A: 0.1% formic acid in water. Buffer B: 0.1% formic acid in 80% acetonitrile.The mass spectrometer was operated in positive ionisation mode with nanospray voltage set at 2.4 kV and source temperature at 305°C. The acquisition was performed in data-dependent acquisition (DDA) mode and full MS scans with 1 micro scans at resolution of 120,000 were used over a mass range of m/z 350-1400 with detection in the Orbitrap mass analyzer. Auto gain control (AGC) was set to ‘auto’ and charge state filtering disqualifying singly charged peptides was activated. In each cycle of data-dependent acquisition analysis, following each survey scan, the most intense ions above a threshold ion count of 10000 were selected for fragmentation. The number of selected precursor ions for fragmentation was determined by the “Top Speed” acquisition algorithm and a dynamic exclusion of 60 seconds. Fragment ion spectra were produced via high-energy collision dissociation (HCD) at normalised collision energy of 28% and they were acquired in the ion trap mass analyzer. AGC was set to 2E4, and an isolation window of 0.7 m/z and a maximum injection time of 12 ms were used. Digested bovine serum albumin (NEB, P8108S) was analysed between each sample to avoid sample carry-over and to assure stability of the instrument and QCloud (45) has been used to control instrument longitudinal performance during the project. The MaxQuant software suite (v1.6.0.16) was used for peptide identification and quantification. The data were searched against a Swiss-Prot mouse database (as in June 2020, 17056 entries) plus Q9CQD0, E9PWZ3, Q3V1Z5 isoforms and a list of common contaminants and all the corresponding decoy entries (46). A precursor ion mass tolerance of 4.5 ppm at the MS1 level was used, and up to two missed cleavages for trypsin were allowed. The fragment ion mass tolerance was set to 0.5 Da. Oxidation of methionine, and protein acetylation at the N-terminal were defined as variable modification; whereas carbamidomethylation on cysteines was set as a fixed modification. Identified peptides and dependent peptides have been filtered using a 5% and 1% FDR respectively. Intensities were normalized by the levels of Atp5a1, a mitochondrial protein with similar intensities across all samples, and used to calculate fold change, p-value and adjusted p-value (FDR) to compare WT vs KO. We manually reviewed the peptides identified “by matching” corresponding to protein RPL3 in WT samples (FQTMEEK, IGQGYLIKDGK and VAFSVAR). Skyline-daily software (v21.1.1.233) (47) was used to extract the area of these peptides and to confirm that their intensity was much less abundant or absent in WT compared to *Rpl3l*^-/-^ samples, thus most likely representing false positive matches. Thus, the intensity of the aforementioned peptides assigned “by matching” was corrected for the analyses and final results. The raw proteomics data have been deposited in the Proteomics Identification Database (PRIDE) (48) repository with the dataset identifier PXD026985.

### PolyA(+) RNA selection from mouse hearts

Poly(A) selection was performed using Dynabeads Oligo(dT)25 beads (Invitrogen, 61002) starting from either HA-pulldown immunoprecipitated RNA (5 to 10 µg of input) or from total RNA from whole hearts (15 µg of input). The beads were first pelleted on a magnetic stand and resuspended in binding buffer (20 mM Tris-HCl, pH 7.5, 1.0 M LiCl, 2 mM EDTA). The beads were repelleted, the buffer was removed, and the beads were resuspended in binding buffer. Total RNA (5 to 10 µg) in DEPC-treated water was mixed 1:1 with binding buffer, and the mixture was immediately heated at 65 °C for 2 minutes. The mixture was then added to the beads, mixed thoroughly by pipetting and incubated on the end-over-end rotator for 10 minutes at room temperature. The samples were placed on the magnet and the supernatant was transferred to a clean tube (for the second round of the poly(A) selection). The beads were washed with washing buffer B (10 mM Tris-HCl, pH 7.5; 0.15 M LiCl; 1 mM EDTA ;10 mM Tris-HCl, pH 7.5) three times, and the supernatant was removed. The first round of poly(A)-containing RNA was eluted with cold 10 mM Tris-HCl, pH 7.5 by heating the mixture at 75 °C for 2 minutes. The supernatant for the second poly(A) selection was denatured by heating at 65 °C for 3 minutes and immediately placed on ice. The beads were resuspended in lysis/binding buffer (100 mM Tris-HCl, pH 7.5; 500 mM LiCl; 10 mM EDTA; 1% LiDS; 5 mM DTT), placed on the magnet and the supernatant was removed. The denatured RNA was added to the beads, mixed thoroughly by pipetting and incubated on an end-over-end rotator for 10 minutes at room temperature. The tube was placed on the magnet and the supernatant was discarded. The washing steps and the elution were performed as previously described, and the elute was mixed with the elute from the first round. In the case of poly(A) selection from whole hearts, the two elutes were mixed with the beads for a third round.

### Nanopore direct cDNA sequencing library preparation

All Nanopore sequencing runs were performed using the MinION sequencer (flow cell type: FLO-MIN106, sequencing kit: SQK-DCS109, barcoding expansion kit: EXP-NBD104). Standard Oxford Nanopore direct cDNA sequencing protocol (version DCB_9091_v109_revC_04Feb2019) was used to sequence mouse total RNA heart samples (input) as well as translated fractions (HA-bound). 100 ng of Poly(A) selected RNA per sample was used for the first strand synthesis reaction, mixed with 2.5 µL of VNP (ONT cDNA sequencing kit), 1 µL of 10 mM dNTPs and filled to 7.5 µL with RNase-free water. The mixture was incubated at 65 °C for 5 minutes and then snap cooled on ice. The following reagents were mixed in a separate tube: 4 µL of 5x RT buffer, 1 µL RNaseOUT (Invitrogen™), 1 µL of RNase-free water and 2 µL of Strand-Switching Primer (SSP, ONT cDNA sequencing kit). The tubes were gently mixed by flicking and incubated at 65 °C for 2 minutes. 1 µL of Maxima H Minus Reverse Transcriptase (Life Technologies, EP0751) was added to the reaction mixture, which was mixed by flicking and incubated for 90 minutes at 42 °C, followed by heat inactivation at 85 °C for 5 minutes. RNA was degraded by adding 1 µL of RNase Cocktail Enzyme Mix (ThermoFisher, AM2286) followed by incubation for 10 minutes at 37 °C. DNA cleanup was performed using AMPure XP beads and quantity and quality were assessed using Qubit™ and Tapestation™. The second strand was synthesized by mixing the following reagents: 25 µL of 2x LongAmp Taq Master Mix (NEB, 174M0287S), 2 µL PR2 primer (ONT cDNA sequencing kit), 20 µL reverse-transcribed sample and 3 µL RNase-free water. The reaction mixture was incubated using the following protocol: 94 °C, 1 minute; 50 °C, 1 minute; 65 °C, 15 minutes; 4 °C, hold. Another AMPure XP beads cleanup step was performed, proceeding to the end-prep step by mixing the following reagents: 20 µL cDNA sample, 30 µL RNase-free water, 7 µL Ultra II End-prep reaction buffer (NEB, E7647A), 3 µL Ultra II End-prep enzyme mix (NEB, E76468). The mixture was incubated at 20 °C for 5 minutes and 65 °C for 5 minutes. After another AMPure XP beads cleanup step, the samples were barcoded by mixing the following reagents: 22.5 µL End-prepped cDNA, 2.5 µL native barcode (NB01-NB12, ONT barcode extension kit EXP-NBD104), 25 µL Blunt/TA ligase master mix. The reaction mixture was incubated for 10 minutes at room temperature, and the barcoded samples were cleaned up using AMPure XP beads. The cDNA amounts were measured using Qubit™, and the samples were pooled together in equal ratios, not exceeding 120 ng (200 fmol) as the maximum total amount of barcoded cDNA. The adapter ligation was performed by mixing together 65 µL of the pooled barcoded sample, 5 µL Adapter Mix II (AMII, ONT cDNA sequencing kit), 20 µL NEBNext Quick Ligation Reaction Buffer 5X (NEB, B6058S) and 10 µL Quick T4 DNA Ligase (NEB, M2200L). The reaction mixture was incubated for 10 minutes at room temperature, after which the cDNA was cleaned up using AMPure XP beads and eluted in 13 μL of Elution Buffer (EB, ONT cDNA sequencing kit). The final amount was ∼50 ng of cDNA, which was mixed with 37.5 µL Sequencing Buffer (SQB) and 2.5 µL Loading Beads (LB, ONT cDNA sequencing kit) and loaded onto a previously primed MinION R9.4.1 flowcell.

### Analysis of cDNA nanopore sequencing data (Nano-TRAP)

Basecalling and demultiplexing of the raw fast5 files was done using Guppy (version 3.6.1) through the MasterOfPores pipeline (version 1.1) (49). Fastq files were then mapped to the GRCm38 mouse genome, in which all non-protein coding sequences were previously masked. Mapping was performed using minimap2 (version 2.14) (50). The bam files were used for the subsequent differential expression analysis, and Gencode’s M25 mouse release was used as the annotation file. Differential expression analysis of nanopore cDNA sequencing runs (Nano-TRAP input and IP) was performed using bambu (version 0.3.0) (doi: 10.18129/B9.bioc.bambu, pre-publication release). DeltaTE (initial release) was used to compute the translation efficiency (51). All code for analysis of Nano-TRAP data is available at : https://github.com/novoalab/RPL3L/tree/main/bambu and https://github.com/novoalab/RPL3L/tree/main/Nano-TRAP_TE.

### Ribosome profiling library preparation

For heart homogenates, mice were sacrificed, and their hearts were quickly excised, washed in PBS containing 100 μg/mL cycloheximide (CHX), and snap frozen in liquid nitrogen. Left ventricular tissue was homogenised using a tissue homogenizer in 5 volumes of ice-cold polysome buffer (20 mM Tris pH 7.4, 10 mM MgCl, 200 mM KCl, 2 mM DTT, 1% Triton X-100, 1U DNAse/μl) containing 100 μg/mL CHX and further homogenised using a 25G needle. For complete lysis, the samples were kept on ice for 10 minutes and subsequently centrifuged at 20,000 g to precipitate cell debris and the supernatant was immediately used in the further steps. From the lysate, 100 μL was used as input, from which RNA was extracted using Trizol. The remaining lysate was used to generate RPFs by treatment with RNAse I for 45 minutes at room temperature. After 45 minutes the reaction was stopped by adding SUPERase RNAse Inhibitor. RPFs were purified using MicroSpin S-400 columns (Cytiva). Purified RPFs were used for the generation of Ribosome profiling libraries using the NEXTFLEX small-RNAseq V3 kit (Perkin-Elmer) according to the user guide. Input RNA libraries were prepared using NEBNext® Poly(A) mRNA Magnetic Isolation Module (ref. e7490) and NEBNext® Ultra II Directional RNA Library Prep Kit for Illumina (24 reactions ref. e7760 or 96 reactions ref. e7765) according to the manufacturer’s protocol, to convert total RNA into a library of template molecules of known strand origin and suitable for subsequent cluster generation and DNA sequencing. For all samples, ribosome profiling library size distributions were checked on the Bioanalyzer 2100 using a high sensitivity DNA assay (Agilent) and input libraries were analysed using Bioanalyzer DNA 1000 or Fragment Analyzer Standard Sensitivity (ref: 5067-1504 or ref: DNF-473, Agilent) to estimate the quantity and validate the size distribution, and were then quantified by qPCR using the KAPA Library Quantification Kit KK4835 (REF. 07960204001, Roche) prior to the amplification with Illumina’s cBot. Libraries were sequenced 1 * 50+8 bp on Illumina’s HiSeq2500.

### Ribosome profiling data analysis

Fastq files were processed and analysed using the RiboToolkit (52) and RiboFlow (initial release) followed by RiboR (53) pipelines. RiboToolkit was used to calculate the codon occupancy at A, E and P sites and the translation efficiency. Briefly, cutadapt (version 1.18) (54) was used to trim 5’ and 3’ adapters and to filter out low quality reads. Trimmed fastq files were uploaded to the RiboToolkit website and the analysis was done using default parameters. RiboFlow was used to process the fastq files (trimming, mapping, filtering reads mapping to rRNA and tRNA sequences, aligning to the transcriptome), and the created ribo objects were loaded into RiboR, which was used to produce the 3nt periodicity plots. Metagene plots were built using the R package ribosomeprofilingQC (https://bioconductor.org/packages/release/bioc/html/ribosomeProfilingQC.html).

### Single nuclei RNAseq library preparation

Single nuclei were obtained from flash-frozen tissues that were dissociated following a previously described method (55). Tissue homogenization was performed using a 7 mL glass Dounce tissue grinder set (Merck) with 8 strokes of a loose and a tight pestle in homogenization buffer (250 mM sucrose, 25 mM KCl, 5 mM MgCl2, 10 mM Tris-HCl, 1 mM dithiothreitol (DTT), 1X protease inhibitor, 0.4 U μL-1 RNaseIn, 0.2 U μL-1 SUPERaseIn, 0.1% Triton X-100 in nuclease-free water). Homogenate was filtered into a 50 mL tube through a 40-μm cell strainer (Corning) and centrifuged (500 g, 5 min, 4 °C) to resuspend the pellet in 500 μL of storage buffer (1× PBS, 4% bovine serum albumin (BSA), 0.2 U μl-1 Protector RNaseIn). Nuclei were stained with NucBlue Live ReadyProbes Reagents (ThermoFisher) and single nuclei were sorted by fluorescent activated cell sorting (FACS). The obtained tube was centrifuged (500 g, 5 min, 4 °C) to get nuclei pellet. Next, 5 µL of single nuclei suspension were mixed with 5µL of Trypan Blue and applied to a Countess II to determine the nuclei concentration. The suspension was adjusted to 800–1,400 cells per µL and loaded to the Chip, targeting the recovery of 5,000 nuclei per sample. The Chip was processed using the Chromium Controller protocol (10X Genomics). Libraries were prepared following the Chromium Single Cell 3’ Reagent Kits User Guide (10X Genomics) protocol. Libraries were sequenced using NovaSeq 6000 flowcell (Illumina).

### Single nuclei RNAseq data analysis

Sequenced single nuclei samples were demultiplexed using bcl2fastq (Illumina) and aligned to the annotated mouse reference genome (mm38, Ensembl v98) with CellRanger (v5.0.0) using default parameters and including introns to globally capture expression of pre-mRNA transcripts. Genomic regions corresponding to annotated ribosomal protein pseudogenes were masked out. Mapped samples were grouped into AnnData objects and analysed by Scanpy v 1.6.0 (56). Doublets were predicted and removed using Solo (score < 0.25) (57) and only nuclei with 300-5,000 expressed genes were retained. A gene was considered as expressed if at least one unique molecular identifier (UMI) was detected in 3 nuclei. Nuclei with less than 200 UMI or an abnormally high proportion of mitochondrial RNAs (≥ 10%) or ribosomal RNAs (≥ 40%), were further removed. Next, UMI were normalised so that every nuclei has the same total UMI count, and nuclei were clustered and visually displayed using the UMAP method (58). Possible sample batches were corrected using Harmony (59). We should note that sNuc-seq captured *Rpl3l* transcripts in nuclei from *Rpl3l*^-/-^ mice (**Figure S1**). However, the 13 bp-long deletion in *Rpl3l*^-/-^ cells was confirmed, since no reads were found to span the CRISPR target sequence (**Figure S1**). *Rpl3l* transcripts are not present in cytosolic extracts of *Rpl3l*^-/-^ hearts (**Figure 2B**), suggesting that they are readily degraded in the cytosol, most likely via nonsense-mediated decay (NMD) mechanisms. The jupyter notebook used for this analysis is available at: https://github.com/novoalab/RPL3L/tree/main/single_nuclei_analysis.

### Imputation of mRNA expression levels in sNuc-seq

MAGIC (v. 0.1.0) (60) was used to impute gene expression corrected for dropout and to recover ribosome paralog pair relationships. MAGIC was run using the table of single nuclei normalized UMI and with default parameters (number pca components = 20, ka = 10, t = 6).

### Animal Ethics

All experimental procedures were approved by the Garvan/St Vincent’s Hospital Animal Ethics Committee, in accordance with the guidelines of the Australian Code of Practice for the Care and Use of Animals for Scientific Purposes (Project No. 16/14 and 16/26). All animals were entered into the study in a randomized order and operators were blinded to genotype and treatments.

## RESULTS

### RPL3L (uL3L) is a vertebrate-specific RP paralog that is expressed postnatally in cardiomyocytes

Previous works have shown that RP paralogs are not equally expressed across mammalian tissues (10). However, their expression patterns across developmental stages have been much less studied. Here, we examined the dynamics of RP paralog expression patterns across both tissues and developmental stages using publicly available RNA-seq datasets (42), which included 7 major organs (brain, cerebellum, heart, kidney, liver, ovary, testis) from embryos (E10.5 to E18.5) and postnatal mice (P0, P3, P14, P28, P63). We found that most RPs were constitutively expressed in all tissues, in agreement with previous observations, and found that these RPs typically showed relatively stable expression levels across developmental stages (**Figure 1A**, see also **Figure S7**). By contrast, the expression patterns of RP paralogs fell into one of three possible behaviours: i) one of the two paralogs is expressed in all tissues, while the other is only expressed postnatally in a single tissue (e.g., *Rpl3*-*Rpl3l, Rpl10*-*Rpl10l, Rpl39*-*Rpl39l*); ii) both paralogs are expressed in similar levels across tissues and developmental stages (e.g., *Rpl22*-*Rpl22l1*) or iii) one of the paralogs is dominantly expressed across tissues and developmental stages, while the other one is either expressed at lower levels or not expressed at all (e.g., *Rpl7*-*Rpl7l1, Rpl36a*-*Rpl36al, Rps27*-*Rps27l*) (**Figure 1B)**.

We observed that RP paralog genes (*Rpl3l, Rpl10l* and *Rpl39l*) that displayed tissue-specific expression patterns also followed similar temporal expression patterns across developmental stages, with their expression levels increasing steeply from embryo to postnatal developmental stages. Moreover, in those tissues in which the tissue-specific RP paralog was expressed, the expression levels of their counterparts (*Rpl3, Rpl10* and *Rpl39*) decreased postnatally, coinciding with increased expression levels of their paralog (**Figure 1A**), and suggesting a possible regulatory interplay between RP paralog pairs.

The *Rpl3l* paralog gene emerged in early vertebrates (**Figure S8A**), and its sequence is relatively similar to that of its paralog gene *Rpl3* (75% sequence identity), with the C-terminus being the most distinct region between the two paralogs (**Figure S8B**). A superimposition of the RPL3L (uL3L) homology model in the ribosome structure shows that the differential C-terminal region is located on the surface of the ribosome (**Figure 1C**), suggesting that RPL3L (uL3L)-containing ribosomes could potentially alter the ability of certain accessory proteins to bind to the ribosome, in a similar fashion to what has been previously observed with RPL36 and RPS17-containing ribosomes (64).

To examine with further detail the expression patterns of *Rpl3l*, we used publicly available single-cell RNA-seq data from mouse hearts (63), revealing that the expression of *Rpl3l* in heart tissues is in fact restricted to cardiomyocyte (CM) cells (**Figure 1D**). Thus, *Rpl3l* expression is not only restricted to specific developmental stages and tissues, but also, its expression is limited to myocyte cell types.

### *Rpl3l* knockout mice show upregulated RPL3 expression in heart and muscle and decreased lean body mass

To reveal the biological function of *Rpl3l*, we generated constitutive knockout mouse models for both *Rpl3* and *Rpl3l* using the CRISPR-Cas9 system (**Figure 2A**, see also **Figure S1**). Depletion of *Rpl3l* in mice led to viable homozygous knockout mice (*Rpl3l*^-/-^), with offspring following mendelian proportions. Knockout of *Rpl3l* was validated both at the mRNA level (**Figure 2B**) as well as at the protein level (**Figure 2C,D**, see also **Figure S4 and S5**). By contrast, depletion of *Rpl3* led to an embryonic-lethal phenotype, and only heterozygous knockout mice (*Rpl3*^+/-^) could be obtained (see *Methods)*.

Histopathological analysis of heart and skeletal muscle tissues (gastrocnemius and quadriceps) from *Rpl3l*^-/-^ mice did not reveal pathological traits nor significant morphological changes compared to control *Rpl3l*^+/+^ mice (**Figure 2E**, see also **Figure S9A** and *Methods*). No statistically significant differences were found between *Rpl3l*^-/-^ and WT mice in heart weight, muscle weight, heart-body weight ratio nor in heart-brain weight ratio (**Figure S9B**, see also **Table S1**). The left and right ventricular free wall thickness was also found to be non-significant, albeit with a moderate but not significant increase in *Rpl3l*^-/-^ left ventricular free wall thickness (p-value = 0.0615). At the physiological level, echocardiographic profiles of *Rpl3l*^-/-^ mice showed no significant differences to those of control *Rpl3l*^+/+^ mice (**Figure S10**, see also **Table S2 and S3**), except a moderate increase in the rate of pressure generation (dP/dtmax, p-value = 0.03). On the other hand, EchoMRI^™^ analysis of body composition of live mice showed a significant increase in total lean mass in aged *Rpl3l*^-/-^ knockout mice when compared to age-matched WT mice (n = 5 vs 5; p-value = 0.02), but not in younger 8 week old mice (n = 5 vs 5; p-value = 0.47) (**Figure 2F**, see also **Figure S9C** and *Methods*).

At the molecular level, we observed that depletion of *Rpl3l* led to a significant increase in *Rpl3* expression levels in heart and skeletal muscle, both at the mRNA (**Figure 2B**) and protein level (**Figure 2C,D**). Thus, a compensatory mechanism upon the absence of *Rpl3l* seems to be present in cardiomyocytes, leading to the production of RPL3-containing ribosomes in cardiomyocytes, a cell type where RPL3-containing ribosomes are normally not present postnatally. This biological compensation provides a biological setup that allows for a functional comparative study of the ribosomal activity of RP paralogs in cardiomyocytes, i.e., by comparing the translational activity of cardiomyocytes with RPL3L-containing ribosomes (in WT mice) or RPL3-containing ribosomes (in *Rpl3l*^-/-^ mice), respectively.

### RPL3L and RPL3 are incorporated into translating ribosomes

RPL3 (uL3) is a highly conserved core ribosomal protein, and it is well-documented that the incorporation of RPL3 (uL3) into ribosomes is of paramount importance for their function (65) (**Figure 1D**). Even though RPL3L has been described as a tissue-specific paralog of RPL3, whether it is actually incorporated into translating ribosomes remains elusive.

To examine the incorporation of RPL3L and RPL3 into translating ribosomes in WT and *Rpl3l*^-/-^ cardiomyocytes, respectively, we performed sucrose gradient fractionation of polysome extracts from WT and *Rpl3l*^-/-^ mice hearts coupled to western blotting (**Figure 3**). Liver was used as a positive control, due to its characteristic and reproducible polysome profiles (**Figure 3A**, top). Protein content was extracted from each polysome fraction, and the fractions were analysed using western blot, by probing them with anti-RPL3L (uL3L), anti-RPL3 (uL3), and anti-RPS6 (eS6) antibodies. Our results showed that RPL3L (uL3L) is found in the monosome and polysome fractions, demonstrating that RPL3L (uL3L) is incorporated into translating ribosomes (**Figure 3A**, middle). Moreover, we observed that upon *Rpl3l* depletion, RPL3 (uL3) compensates for the missing paralog by replacing it in the 60S subunit, and is incorporated into translating ribosomes from *Rpl3l*^-/-^ hearts (**Figure 3A**, bottom). As additional control, WT and *Rpl3l*^-/-^ heart polysome profile fractions were inversely probed with anti-RPL3 and anti-RPL3L (uL3L) antibodies, respectively, showing that RPL3 (uL3) is also incorporated into monosomes isolated from a whole WT heart, albeit at lower levels when compared to monosomes of a *Rpl3l*^-/-^ heart (**Figure 3B**, top). RPL3 signal in WT hearts most likely arises from ribosomes present in non-cardiomyocyte cell types present in the heart, as shown in expression profiles obtained using single-cell RNA-seq (**Figure 1D)**. Conversely, RPL3L (uL3L) was not detected in any of the *Rpl3l*^-/-^ polysome profile fractions (**Figure 3B**, bottom).

Finally, we wondered whether the incorporation of RPL3 (uL3) and RPL3L (uL3L) into actively translating ribosomes might lead to global changes in protein translation. To explore this, we performed puromycin incorporation assays (66) on hearts from WT and *Rpl3l*^-/-^ mice. Analyses of puromycin incorporation levels did not show significant differences, suggesting that the distinct use of RP paralogues, namely RPL3L (uL3L) in WT mice and RPL3 in *Rpl3l*^*-/-*^ mice, does not globally affect protein translation rates of heart tissues (**Figure 3C**).

### The use of RPL3L does not lead to preferential translation or enhancement of translation efficiency

Previous works have shown that heterogeneity in RP composition endows ribosomes with differential selectivity for translating subpools of transcripts including those controlling metabolism, cell cycle, and development (7). Specifically, it was found that RPL10A and RPS25-bearing ribosomes showed preferential translation of subsets of transcripts when compared to the whole ribosome population. However, it is unclear whether the use of RP paralogs would lead to preferential translation of specific subsets of transcripts.

Here we examined this question by testing whether RPL3L (uL3L)-containing ribosomes showed preferential translation of subsets of transcripts, relative to RPL3 (uL3)-containing ribosomes. To this end, we devised a novel method, which we termed Nano-TRAP: HA-tag mediated Translating Ribosome Affinity Purification (TRAP) (67) coupled to nanopore cDNA sequencing (**Figure 4A**, see also *Methods*). Nano-TRAP captures full-length mRNAs that are associated with at least one 80S ribosome. We applied Nano-TRAP in biological triplicates to mouse hearts that were isolated from either *Rpl22*-HA^+/+^/*Rpl3l*^-/-^ or control *Rpl22*-HA^+/+^/*Rpl3l*^+/+^ hearts corresponding to mRNA populations that are bound to RPL3 (uL3)- and RPL3L (uL3L)-containing ribosomes, respectively. We first examined the replicability of Nano-TRAP, finding that per-gene log2-counts across biological replicates were highly replicable (pearson r^2^ = 0.886-0.939) (**Figure S11A**,**B)**. We then performed differential expression (DE) analysis (see *Methods*) to examine whether the ribosome-bound mRNA populations would be significantly distinct upon knock-out of *Rpl3l*, finding only 5 differentially-bound transcripts (**Figure 4B**, see also **Table S4**). Our results identified *Rpl3* as one of the differentially-bound mRNAs, in agreement with our previous results that showed upregulation of *Rpl3* upon *Rpl3l* depletion (**Figure 2B**). In addition to increased binding to *Rpl3* transcripts, RPL3L (uL3L)-depleted ribosomes showed significantly decreased binding towards 2 genes: *Rbp7* and *Ankrd1*, the latter being a transcription factor whose dysregulation has been associated with several types of cardiomyopathies (68, 69) including hypertrophic cardiomyopathy.

Finally, we examined whether translation efficiency (TE) would be significantly altered upon *Rpl3l* depletion. TE was calculated by dividing normalised counts from the IP experiment by those from the input experiment. We then compared the TE values between WT and *Rpl3l*^-/-^ strains; however, most transcripts did not show significant changes in their TE values (**Figure 4C**, see also **Table S4**), and the few genes that showed significant TE changes were not enriched for any GO term (**Table S5)**. Altogether, our analyses using Nano-TRAP suggest that the use of RPL3L (uL3L) instead of RPL3 (uL3) in translating ribosomes does not globally lead to preferential translation of subsets of transcripts in mouse hearts.

### *Rpl3l* depletion does not affect the distribution of ribosomes along transcripts

It has been reported that RPL3, which is located close to the peptidyl-transferase centre in the ribosome, plays a role in preventing ribosomes from piling up in the mRNA 5’ region early during translation elongation (65). Therefore, we wondered whether the presence of RPL3 (uL3) or RPL3L (uL3L) in ribosomes might lead to changes in the distribution of ribosomes along specific transcripts.

To examine this, we performed ribosome profiling (Ribo-Seq) (70) in WT and *Rpl3l*^-/-^ mouse hearts (**Figure 4D**, see also **Figure S12)**, in biological triplicates, finding a consistent 3-nucleotide periodicity in ribosome-protected (RPF) libraries in all samples (**Figure S13A**). We first examined whether the depletion of *Rpl3l* led to accumulation of reads in specific genic regions. To this end, we analysed the distribution of reads of ribosome-protected footprints (RPFs) at the metagene level; however, we did not observe statistically significant differences between WT and *Rpl3l*^-/-^ RPF reads, finding that reads mapped in similar proportions to the 3’ UTR, CDS and 5’ UTR (**Figure 4E**). Similarly, metagene analyses showed that RPFs did not accumulate in specific regions of the transcript (**Figure 4F**, see also **Figure S14**).

We then assessed whether Ribo-Seq, when integrated with RNA-seq, would be able to identify transcripts with differential mRNA expression levels (input mRNA), ribosomal occupancy (RFP), or translational efficiency (TE). This suggests that *Rpl3l* depletion did not globally lead to changes in transcription, translation or translation efficiency (**Figure 4G**), in agreement with our previous observations using Nano-TRAP. Finally, we examined whether RPL3 (uL3) and RPL3L (uL3L)-containing ribosomes would show differences in ribosome codon occupancies; however, we did not find significant differences between WT and *Rpl3l*^-/-^ hearts (**Figure 4H**, see also **Figure S13B**).

### *Rpl3l* expression is mutually exclusive with *Rpl3* expression in cardiomyocytes

Our results show that *Rpl3l* depletion leads to upregulation of *Rpl3* expression levels, which is concordantly detected both at the mRNA (**Figure 2B** and **4B**, see also **Figure S15**) and protein levels (**Figure 2C,D**). Intrigued by this compensatory mechanism, we performed single nuclei RNA sequencing (sNuc-seq) (71–73) on left ventricles from *Rpl3l*^+/+^ and *Rpl3l*^-/-^ mice, in biological triplicates. Compared to single cell RNA-seq, sNuc-seq improves the profiling of gene expression in cells which are difficult to dissociate from tissues (73).

Analyses of sNuc-seq datasets confirmed that depletion of *Rpl3l* showed no major transcriptomic changes in cardiomyocytes (**Table S7**), in agreement with our previous observations (**Figure 4B** and **S10C**). Moreover, we confirmed that *Rpl3l* is cardiomyocyte-specific, whereas *Rpl3* is mainly expressed in non-cardiomyocyte cells (**Figure S16A**). To examine the interplay between *Rpl3* and *Rpl3l* expression in cardiomyocytes, we imputed gene expression levels across all the identified single nuclei (see *Methods*), finding that *Rpl3* and *Rpl3l* are expressed in a mutually exclusive manner (**Figure S16B, Table S8**). By contrast, this mutual exclusivity was not observed in any of the other ribosomal protein paralog pairs expressed in the heart (**Figure S16C and S14**). Altogether, our results suggest that the presence of RPL3L (uL3L) in cardiomyocytes is directly responsible for the lack of RPL3 (uL3) in a given cell.

### RPL3-bearing ribosomes establish physical contact with mitochondria

Our analyses show that *Rpl3* and *Rpl3l* protein-coding sequences significantly differ in their C-terminal region (**Figure S8A**), which protrudes towards the outer region of the ribosome (**Figure 1C**). We therefore reasoned that the incorporation of RPL3L (uL3L) or RPL3 (uL3) in ribosomes could lead to the recruitment of different ribosome-associated proteins (RAPs) (64), which could in turn alter the function or fate of the ribosome.

In order to study how *Rpl3l* depletion affects ribosomal protein composition, we employed antibody immunoprecipitation (pulldown of RPL22-(eL22)-HA-tagged ribosomes from WT or *Rpl3l*^-/-^ cardiomyocytes, respectively) coupled to mass spectrometry, which we termed Proteo-TRAP (**Figure 5A**). Our results confirmed that RPL3L (uL3L) was depleted from *Rpl3l*^-/-^ cardiomyocytes at the protein level, while RPL3 was upregulated (**Figure 5B**, see also **Table S9**). The only other RP that was significantly overrepresented in *Rpl3l*^-/-^ cardiomyocytes was RPL38, suggesting that the *Rpl3l* knockout does not globally lead to substantial alterations in terms of ribosomal protein composition (**Table S10**).

To our surprise, ribosomes isolated from *Rpl3l*^-/-^ cardiomyocytes, which have RPL3-containing ribosomes, showed a dramatic enrichment in interactions with mitochondrial proteins. By contrast, this enrichment was absent in WT cardiomyocytes, which have RPL3L (uL3L)-containing ribosomes (**Figure 5C**, see also **Figure S17**). GO term enrichment analysis (74) confirmed that interacting proteins were significantly enriched for mitochondria-related molecular functions and biological processes (**Figure 5D**). We reasoned that the presence of mitochondrial proteins could be caused by increased physical contact between RPL3 (uL3)-containing ribosomes and mitochondria, relative to RPL3L (uL3L)-containing ribosomes. In this regard, it has been shown that in yeast, ribosomes can localise to the outer mitochondrial membrane, supporting co-translational transport of mitochondrial proteins into the mitochondria (75), with electron cryo-tomography images supporting this hypothesis (76). Moreover, it is well documented that mammalian mitochondria also contain ribosome receptors on their outer membrane (77). To examine whether RPL3 (uL3) and RPL3L (uL3L)-containing ribosomes might show distinct subcellular localizations, we performed western blot analyses of cytosolic and mitochondrial fractions of WT and *Rpl3l*^-/-^ hearts, and found that RPL3 localised to both the cytosolic and the mitochondrial fraction, while RPL3L (uL3L) was almost exclusively detected in the cytosolic fraction (**Figure 5E**). Thus, we conclude that RPL3-containing ribosomes show altered subcellular localization relative to RPL3L (uL3L)-containing ribosomes, causing an increased presence of mitochondrial proteins upon ribosome pulldowns.

Previous works have shown that RPs, including RPL3 (uL3), can be SUMOylated post-translationally (78, 79). In this regard, we observed that the RPL3 (uL3) protein that is located in the mitochondrial fraction was of slightly larger size than the one found in the mitochondrial fraction (∼46kDa) (**Figure 5E**, see also **Figure S6**). However, the addition of SUMO would account for a ∼10 kDa difference, which is larger than ∼3 kDa that we observed (**Figure S6B**). Thus, RPL3 (uL3) could be a target of yet uncharacterized post-translational modifications, which could account for the observed size difference between the cytosolic and mitochondrial forms. Alternatively, distinct isoforms could be present in mitochondrial-bound RPL3 (uL3), relative to cytosolic RPL3 (uL3).

Finally, we examined whether the change in subcellular localization towards the mitochondria would be accompanied by altered mitochondrial function. To this end, we performed luminometric measurement of ATP levels in WT and *Rpl3l*^-/-^ cardiomyocytes, finding a significant increase in ATP production upon RPL3L (uL3L) depletion (**Figure 5F**). We should note that this global increase in ATP levels was not caused by increased abundance of mitochondria in RPL3L (uL3L)-depleted cardiomyocytes (**Figure S18**). Future work will be needed to better comprehend how the increased presence of ribosomes bound to the mitochondria leads to increased mitochondrial function.

### Hypertrophy leads to increased *Rpl3* and decreased *Rpl3l* levels in the heart

Cardiac hypertrophy is an adaptive response to pressure or volume stress, mutations of certain proteins, or loss of contractile cardiac mass from prior infarction (80). Hypertrophy can occur as a compensatory consequence of pressure overload; however, recent studies raise the prospect of modulating hypertrophy to afford clinical benefit without provoking hemodynamic compromise (81, 82). To accomplish this goal, it is essential to identify the molecular events that distinguish pathological hypertrophy versus physiological hypertrophy.

Previous works have shown that, in skeletal muscle, hypertrophy leads to increased *Rpl3* and decreased *Rpl3l* mRNA levels (15). Thus, we wondered whether a similar phenomenon may also occur in the hypertrophic heart. To this end, we examined RP mRNA expression levels upon hypertrophic stimuli using publicly available RNA-seq datasets (63, 83, 84). Analysis of scRNA-Seq datasets from resting and hypertrophic hearts (2, 5, 8 and 11 weeks after transverse aortic constriction (TAC) surgery) (63) revealed that *Rpl3l* levels in cardiomyocytes steadily decrease upon hypertrophic stimulus, being *Rpl3l* almost undetectable at week 11 (**Figure 6A**, see also **Figure S18**). Similar results were observed when examining RP expression patterns in bulk RNA-Seq data from TAC-induced hypertrophic and control hearts (83). Specifically, *Rpl3/Rpl3l* mRNA ratios were consistently altered in all timepoints after surgery (2, 4, 7 and 21 days) (**Figure 6B)**, while this trend was not seen in other RP paralog pairs (**Figure S20** and **S21**). Finally, we examined whether the interplay between *Rpl3/Rpl3l* might be altered upon knockout of *Lin28a (84)*, an RNA-binding protein that directly binds and increases mitochondrial phosphoenolpyruvate carboxykinase 2 (*Pck2*) mRNA levels, which has been shown to play an important role in maintaining cardiac hypertrophy (84). Indeed, we found that upon *Lin28a* knockout, the interplay between *Rpl3* and *Rpl3l* expression patterns upon hypertrophic stimulus was impaired (**Figure 6C**, see also **Figure S22)**.

**Figure 6.**
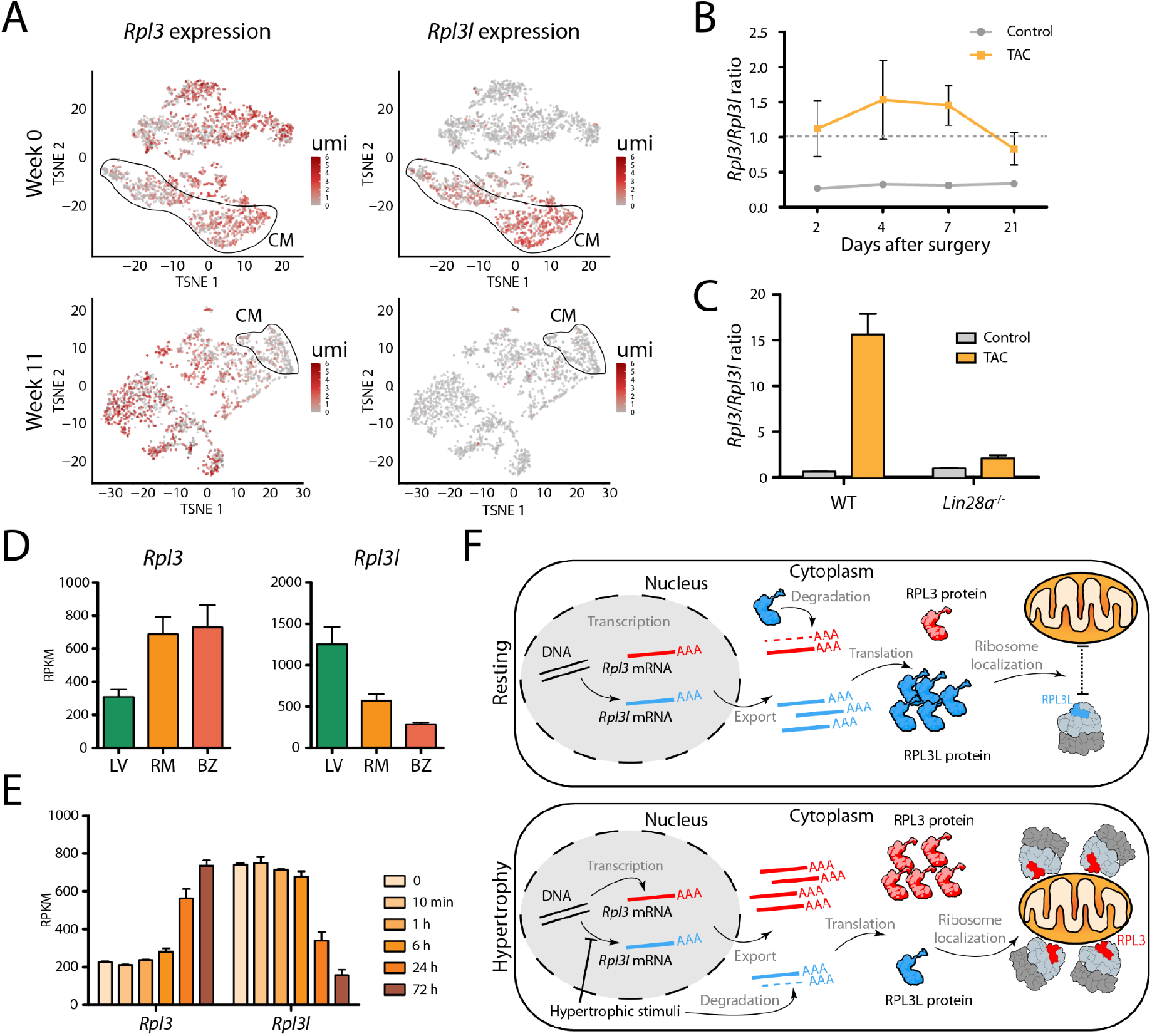
Pressure overload leads to an increase in *Rpl3* expression and a decrease in *Rpl3l* expression in the heart. **(A)** The expression of *Rpl3* and *Rpl3l* across cell types in mouse hearts before (week 0) and after hypertrophy (week 8) induced by transverse aortic constriction (TAC). Cardiomyocytes (CM) are circled. Expression levels are shown as umi (unique molecular identifiers). **(B)** TAC leads to heart hypertrophy that correlates with increased *Rpl3* expression and decreased *Rpl3l* expression. The *Rpl3*/*Rpl3l* ratio is significantly increased in all timepoints (2, 4, 7 and 21 days after surgery) when compared to control hearts. See also Figure S19 and S20. **(C)** Effects of TAC-induced hypertrophy on *Rpl3*-*Rpl3l* interplay are impaired in *Lin28a*^-/-^ mice. See also Figure S22. **(D)** *Rpl3* (left) and *Rpl3l* (right) nuclear mRNA levels in cardiomyocytes from the left ventricle (LV), remote myocardium (RM) and border zone (BZ) after myocardial infarction. Processed data (RPKM) was obtained from Günthel et al. (85). **(E)** *Rpl3* and *Rpl3l* mRNA expression levels at different time points (0 min, 10 min, 1 h, 6 h, 24 h, 72 h) after myocardial infarction. Processed data (RPKM) was obtained from Liu et al. (86). **(F)** Model showing the *Rpl3*-*Rpl3l* interplay in resting (top) and hypertrophic (bottom) conditions. In the resting heart, *Rpl3l* is predominantly expressed in cardiomyocytes, and the *Rpl3l* protein negatively regulates *Rpl3* expression, while RPL3L (uL3L)-containing ribosomes do not establish close contact with mitochondria. Upon hypertrophic stimuli, *Rpl3l* expression is impaired and *Rpl3l* mRNA is degraded in the cytoplasm, leading to an increased expression of *Rpl3*. RPL3 (uL3)-containing ribosomes establish close contact with mitochondria.

We then wondered whether other heart-related conditions, such as myocardial infarction, may be accompanied by a switch in *Rpl3*/*Rpl3l* expression patterns. To this end, we analyzed RNA-seq data performed on cardiomyocyte nuclei isolated from hearts after myocardial infarction (85), finding that *Rpl3* levels were increased both in the ‘border zone’ (the region most affected by the infarction) and ‘remote myocardium’ (mildly affected by the infarction), relative to the left ventricle (unaffected heart region) **(Figure 6D)**. The increase in *Rpl3* levels was accompanied by a decrease in *Rpl3l* levels in the infarcted areas, suggesting that the *Rpl3-Rpl3l* transcriptional switch occurs not only upon hypertrophy, but also infarction. Analysis of a second RNAseq dataset consisting of a time-course of mouse hearts induced with myocardial infarction(86) revealed that the *Rpl3-Rpl3l* switch is already detectable after 6 hours post-infarction, while maximal change in *Rpl3-Rpl3l* levels was observed 72 hours post-infarction (**Figure 6E**). Altogether, our results point to a molecular interplay between *Rpl3* and *Rpl3l* in cardiomyocytes upon hypertrophic stimuli (**Figure 6F**) and infarction, suggesting that *Rpl3l* holds promise as potential target for therapeutic intervention.

## DISCUSSION

In recent years, compelling evidence has accumulated supporting ribosome composition heterogeneity (2, 7, 9, 10, 87–89). At the level of protein composition, ribosomes lacking specific ribosomal proteins have been shown to exist within mouse embryonic stem cells (7, 9), yeast (3, 17, 18, 90) and bacteria (19), though questions remain as to the functionality of such ribosomes (11). At the level of rRNAs, several studies have shown that these can be expressed from separate genomic loci, with the resulting rRNA molecules differing significantly in their sequences. For example, zebrafish express rRNAs from two separate loci: one locus serves for maternal rRNA transcription, and the other for zygotic rRNA (91). Similarly, in *Plasmodium falciparum*, different diverging copies of rRNA are encoded in the genome, one of which is utilised during the mosquito stage and another during the human stage of the infection (92, 93).

Ribosomal protein (RP) paralogs are present in a wide range of eukaryotic species, but their use as a source of ribosome heterogeneity, as well as their functional relevance, remains unclear. In vertebrates, RP paralogs are believed to have appeared via duplication of the ancestral RP gene (94), can show constitutive or restricted tissue expression profiles (10, 94), and their depletion or mutation can lead to tissue-specific phenotypes (95, 96). In the case of mice, we find 7 RP paralogs, which display 39-99% sequence identity to their ancestral gene (**Table S13**). Here, we examined the functional relevance of RP paralogs in mammals by focusing on the biological role of RPL3L (uL3L) in mice, to unveil why striated muscles in vertebrates evolved to express a distinct version of RPL3, a core RP essential for ribosomal function. Our findings point to a highly specialized function of RPL3L (uL3L), with its expression being tightly regulated at the temporal level (postnatal expression), restricted to few cell types (myocytes) and in limited subsets of tissues (striated muscles) (**Figure 1A,D**). Moreover, we find that the C-terminus of RPL3L (uL3L) is the most distinct region when compared to RPL3 (**Figure 1B**, see also **Figure S8A**), which is well-conserved among mammalian RPL3L (uL3L) proteins (**Figure S8C**), suggesting its importance to fulfil its specialized function.

By generating *Rpl3* and *Rpl3l* knockout mouse models, we found that ubiquitously expressed *Rpl3* is of paramount importance for mouse development, with homozygous *Rpl3* knockout being embryonic lethal. We should note that this is not the case of other constitutively expressed RP proteins, such as *Rpl22* (97). *By contrast, we successfully generated Rpl3l*^-/-^ homozygous knockout mice, and identified a rescue mechanism in which RPL3 expression levels are upregulated in striated muscles upon *Rpl3l* depletion (**Figure 2B-D**, see also **Figure S5**). *Rpl3l*^-/-^ mice showed no apparent histological or physiological aberrations (**Figure S9**), with no significant differences in their echocardiogram profiles when compared to age-matched wild type mice (**Table S3** and **Figure S10**). By contrast, 55 week old *Rpl3l*^-/-^ mice had significantly lower lean mass when compared to age matched wild type mice (**Figure 2F**), suggesting that RPL3L (uL3L) might be important for muscle maintenance with ageing.

Previous work had already reported an inverse correlation of *Rpl3* and *Rpl3l* expression upon muscle hypertrophy (15), which we also observed upon *Rpl3l* knockout (**Figure 2B-D**, see also **Figure S15**); however, it remains unclear whether RPL3L (uL3L) is incorporated into translating ribosomes, as its putative role could be exclusively extraribosomal. To test this, we performed polysome profiling coupled to western blot analysis on WT and *Rpl3l*^-/-^ hearts, showing that RPL3L (uL3L) is incorporated into polysomes in WT heart, while it is being substituted by RPL3 (uL3) in *Rpl3l*^-/-^ hearts (**Figure 3**). Therefore, *Rpl3l* knockout mice offer a model system in which cardiomyocyte ribosomes incorporate RPL3 (uL3) instead of RPL3L (uL3L), thus allowing us to study how the use of RPL3 (uL3) or RPL3L (uL3L) in the ribosome influences translation in i*n vivo* mouse models. We should note that neither iPSC-derived myocytes nor neonatal myocytes can be used as a model system to study the RPL3 (uL3)-RPL3L (uL3L) interplay that occurs upon hypertrophy as RPL3L (uL3L) is not expressed in embryonic or neonatal myocytes (42).

To examine the molecular function of RPL3L (uL3L) *in vivo*, we first performed Ribo-Seq experiments on both WT and *Rpl3l*^-/-^ hearts, finding that the use of RPL3- or RPL3L-containing ribosomes did not lead to altered translation efficiency of subsets of transcripts (**Figure 4G**). To confirm these observations, we devised a novel method to study preferential translation, consisting of ribosome pulldown (using HA-tagged ribosomes) coupled to Nanopore direct cDNA sequencing, which we termed Nano-TRAP. Ribo-Seq is a well-established method that relies on sequencing of ribosome-protected fragments (RPFs) using short read sequencing technologies; ∼30 nucleotide-long RPFs allow for analyses at the subcodon resolution, but such short fragments also lead to multi-mapping issues, as well as loss of isoform-specific information. On the other hand, Nano-TRAP relies on long-read sequencing, allowing for the analysis of full transcripts and per-isoform analyses, alleviating the mappability problems of RPFs. Furthermore, Nano-TRAP could also be customized for direct-RNA sequencing instead of direct cDNA sequencing, allowing for translatome-wide analysis of mRNA modifications.

Our Nano-TRAP analyses confirmed that *Rpl3l* depletion does not lead to preferential translation of transcripts (**Figure 4C**), in agreement with our observations using Ribo-Seq. Therefore, both orthogonal methods show that *Rpl3l* depletion does not lead to global accumulation of ribosome footprints along the transcript nor preferential translation of subsets of transcripts, compared to WT samples. We should note that our datasets were not significantly different in terms of coverage than previously published datasets (98), for which we successfully identified differential translation of subsets of transcripts when using the same bioinformatic pipeline (**Table S14**, see also **Figure S23**), suggesting that sequencing coverage was not a limiting factor in the identification of differentially translated transcripts in our experiments.

We then investigated whether the stoichiometry of ribosomal proteins (RPs) or ribosome-associated proteins (RAPs) might be altered upon *Rpl3l* depletion. To this end, we performed proteomic analyses of ribosome immunoprecipitates (Proteo-TRAP), in a similar fashion to previous works employing FLAG-tagged ribosomes coupled to mass spectrometry analyses (7). Unexpectedly, we found a drastic enrichment of genome-encoded mitochondrial proteins in RPL3 (uL3)-containing ribosome immunoprecipitate fractions, compared to RPL3L (uL3L)-containing fractions (**Figure 5C**). The fact that such a large number of mitochondrial proteins were found in ribosome immunoprecipitates in RPL3 (uL3)-containing ribosomes (*Rpl3l*^-/-^/*Rpl22HA*^+/+^) but not in RPL3L (uL3L)-containing ones (*Rpl3l*^+/+^/*Rpl22HA*^+/+^) despite stringent washing steps (see *Methods*) suggests a strong physical contact between RPL3-containing ribosomes and mitochondria.

It is well-documented that ribosomes can bind to the outer mitochondrial membrane, allowing for co-translational transport of mitochondrial proteins encoded by the nuclear genome (77). Such findings have been further corroborated by visualisation of cytosolic ribosomes on the mitochondrial surface by electron cryo-tomography (76). In yeast, OM14 has been identified as the mitochondrial receptor of the nascent chain-associated complex, allowing for the binding of ribosomes on the mitochondrial surface (75). In agreement with these observations, previous studies in yeast have shown that certain strains deficient in specific RP paralogs (*Rpl1b, Rpl2b* and *Rps26b*) display altered mitochondrial morphology and function (24). Moreover, RPL3 (uL3), in its ribosome-free form, has been shown to have the ability to localise to mitochondria (99). Based on these previous observations, we speculated that the incorporation of RPL3 into cardiomyocyte ribosomes might lead to tighter ribosome-mitochondria interactions, compared to RPL3L (uL3L)-containing ribosomes, potentially affecting mitochondrial function. To test this, we performed western blot analyses of mitochondrial and cytosolic fractions of WT and *Rpl3l*^-/-^ hearts, finding that RPL3L (uL3L) is only detected in the cytosolic fraction, whereas RPL3 localises both to the cytosolic and the mitochondrial fractions (**Figure 5E**). We then tested whether the localization of RPL3-containing ribosomes to mitochondria affected their function, finding that the use of RPL3 in ribosomes (present in *Rpl3l*^*-/-*^ cardiomyocytes) was accompanied by a significant increase in ATP levels (**Figure 5F**), compared to WT cardiomyocytes with RPL3L (uL3L)-containing ribosomes. Future work will be needed to further dissect how the *Rpl3*-*Rpl3l* interplay regulates mitochondrial activity.

Previous studies focusing on hypertrophy have shown that RPL3L (uL3L) levels decrease and RPL3 (uL3) levels increase in muscle upon hypertrophic stimulus (15) as well as in Duchenne Muscular Dystrophy (100), supporting its importance in muscle function. Similarly, our analyses using publicly available RNAseq datasets showed that RPL3L (uL3L) is downregulated in cardiomyocytes upon hypertrophic stimuli (**Figure 6A-C** see also **Figure S18**) as well as upon myocardial infarction (**Figure 6D,E)**. Therefore, our *Rpl3l* knockout mouse models may constitute a promising *in vivo* model to study hypertrophy and myocardial infarction. Notably, a recent study showed that knockdown of RPL3L (uL3L) improved muscle function (100), but the underlying molecular mechanism remains unclear. We propose that the improved muscle function observed in these studies (15, 100) might be a direct consequence of altered mitochondrial activity upon RPL3L (uL3L) downregulation, in a similar fashion to what we observe upon *Rpl3l* depletion (**Figure 5F**).

The dynamic interplay between mammalian RP paralogs is not an exclusive feature of the *Rpl3-Rpl3l* pair, but has also been reported to occur in other paralog pairs, such as the *Rpl22-Rpl22l1* pair. Specifically, homozygous depletion of *Rpl22* in mice is compensated by an upregulation of its paralog RPL22L1 (eL22L1) (97), which is otherwise largely absent in its protein form in most adult tissues (**Figure 1A**). Intrigued by this molecular interplay, we examined previously published genome-wide epigenetic chromatin states (101) of distinct RP paralog pairs across a wide variety of tissues, finding that both *Rpl3* and *Rpl3l* show active transcription chromatin states in adult striated muscle tissues but not in other tissues (**Figure S3**), suggesting that both genes are actively transcribed, and therefore, that the interplay between *Rpl3* and *Rpl3l* most likely occurs at the mRNA level. In agreement with these observations, a recent study in yeast showed that the interplay between *Rpl22a* and *Rpl22b* occurred at the mRNA level, where the RPL22A (eL22) protein was shown to bind the intron of the unspliced *Rpl22b* RNA transcript, causing its degradation (102). We speculate that a similar regulatory mechanism could be present in vertebrates, and could potentially explain how the *Rpl3-Rpl3l* interplay that occurs upon hypertrophic stimuli is achieved (**Figures S19** and **S21**).

RPL3 (uL3) is a highly conserved core ribosomal protein that plays an important role in ribosome function, and mutation of key residues of yeast RPL3 (uL3) has been shown to cause altered peptidyltransferase activity and frameshifting efficiency (103). On the other hand, methylation of H243 in *S. cerevisiae* has been shown to affect translation elongation and translational fidelity (104). More recently, studies in yeast have shown that the mutation of W255, the residue that is closest to the peptidyl-transferase centre, affected diverse aspects of ribosome biogenesis and function (65). Thus, we speculate that while the selective use of RPL3 (uL3) and RPL3L (uL3L) in cardiomyocytes does not affect translation efficiency, it might be affecting translational fidelity. Future work will be needed to decipher whether the incorporation of RPL3 (uL3) or RPL3L (uL3L) into ribosomes might be significantly affecting translational fidelity in mammals.

Altogether, our work shows that *Rpl3l* is a cell-type specific paralog of *Rpl3* that is restricted to myocyte cells in adult striated muscle tissues, which is downregulated upon hypertrophic stimuli. We show that although RPL3L (uL3L) is incorporated into actively translating ribosomes, it does not lead to preferential translation nor to altered translation efficiency, as demonstrated by Ribo-Seq and Nano-TRAP. By contrast, we show that the use of RPL3L (uL3L) or RPL3 (uL3) modulates the subcellular location of ribosomes and mitochondrial activity in cardiomyocytes. Our work expands our understanding of mammalian RP paralogs and their functional relevance *in vivo*, supporting the view that ribosome heterogeneity is accompanied by functional specialization.

## DATA AND CODE AVAILABILITY

Nano-TRAP (IP and Input) and Ribo-Seq (RPF and mRNA) FASTQ files have been deposited in the Gene Expression Omnibus (GEO), under accession codes GSE189872 and GSE189854, respectively. Single nuclei RNA-Seq data has been deposited in the European Nucleotide Archive (ENA) at EMBL-EBI under accession number PRJEB49255. Proteomics data has been deposited in the Proteomics Identification Database (PRIDE), under accession code PXD026985. A complete list of samples and corresponding codes can be found in **Table S14**. Raw immunofluorescence images have been deposited in Figshare (https://doi.org/10.6084/m9.figshare.19617165.v1). All scripts used in this work, including those used to perform single cell data analysis, Nano-TRAP analyses, Ribo-Seq analyses, phylogenetic trees and heatmaps of RP expression patterns, have been made publicly available in GitHub (https://github.com/novoalab/RPL3L).

## ACKNOWLEDGEMENTS

We thank all the members of the Novoa lab for their valuable insights and discussion. We thank Dr. Elisenda Sanz Iglesias for her invaluable insights regarding the RiboTag mice. We thank Dr. Rob Brink for all his help and discussions on diverse strategies to obtain the CRISPR knockout mice strains used in this work. We thank Prof. John McCarthy for providing us with an aliquot of the RPL3L (uL3L) antibody, as well as for insightful discussions on the role of RPL3L (uL3L). We thank Prof. Dr. Antonio Zorzano and Dr. David Sebastián for the discussions and insights regarding mitochondrial biology. We thank Dr. Jorge Ferrer for providing us with a cre mouse line. We thank Oliver Hummel for his help uploading the generated sNuc-seq datasets to the European Nucleotide Archive (ENA) repository. We thank Ana Arsenijevic for her help with the PyMOL structures. IM is supported by “la Caixa” InPhINIT PhD fellowship (LCF/BQ/DI18/11660028). This project has received funding from the European Union’s Horizon 2020 research and innovation programme under the Marie Skodowska-Curie grant agreement No. 713673. This work was supported by the Australian Research Council (DE170100506 to EMN) and the Spanish Ministry of Economy, Industry and Competitiveness (MEIC) (PGC2018-098152-A-100 to EMN). NH is the recipient of an ERC advanced grant under the European Union Horizon 2020 Research and Innovation Program (AdG788970). NH is supported by a grant from the Leducq Foundation (16CVD03) and a grant by the Chan Zuckerberg Foundation (2019-20266). We acknowledge the support of the MEIC to the EMBL partnership, Centro de Excelencia Severo Ochoa and CERCA Programme / Generalitat de Catalunya. The CRG/UPF Proteomics Unit is part of the Spanish Infrastructure for Omics Technologies (ICTS OmicsTech) and it is a member of the ProteoRed PRB3 consortium which is supported by grant PT17/0019 of the PE I+D+i 2013-2016 from the Instituto de Salud Carlos III (ISCIII) and ERDF.

## AUTHOR CONTRIBUTIONS

IM performed most of the experiments and bioinformatic analyses described in this work, including mouse tissue collection and colony management, mice genotyping, cardiomyocyte preparations, total RNA and protein extractions, western blotting, immunofluorescence assays, polysome profiling, immunoprecipitation, nanopore sequencing, Nano-TRAP and RiboSeq data analyses. HGSV performed sucrose gradients, characterised RP antibodies across tissues, and contributed to the set up of mouse tissue polysome profiles. MCL contributed to mouse tissue collections and cardiomyocyte isolations. GP built the sc-nucRNA libraries, and JR performed the sc-nucRNAseq data analyses, which were supervised by NH and SvH. SK and JW performed echocardiography experiments, which were supervised by MF. GE and ES performed mass spectrometry experiments. MV built the Ribo-Seq libraries. EMN conceived and supervised the work. IM and EMN wrote the manuscript, with the contribution from all authors.

