## Supplementary figures for "Dynamic interplay between RPL3- and RPL3L-containing ribosomes modulates mitochondrial activity in the mammalian heart"

**Figure S1. Validation of *Rpl3l* knockout in mice using sNuc-seq datasets and Sanger sequencing.** **(A)** Schematic overview of the GENCODE version M23 (Ensembl 98) annotated transcripts in the *Rpl3l* locus in *M. musculus*. In the zoomed area of the figure, the 13 bp deletion generated via CRISPR-Cas9 in *Rpl3l* knockout mice is underlined in red and boxed in black. **(B)** Sanger sequencing of WT and *Rpl3l*$^{-/-}$ mice, in biological triplicates. The aligned sequences are showing the region in which the 13 bp long deletion was introduced (boxed in red).

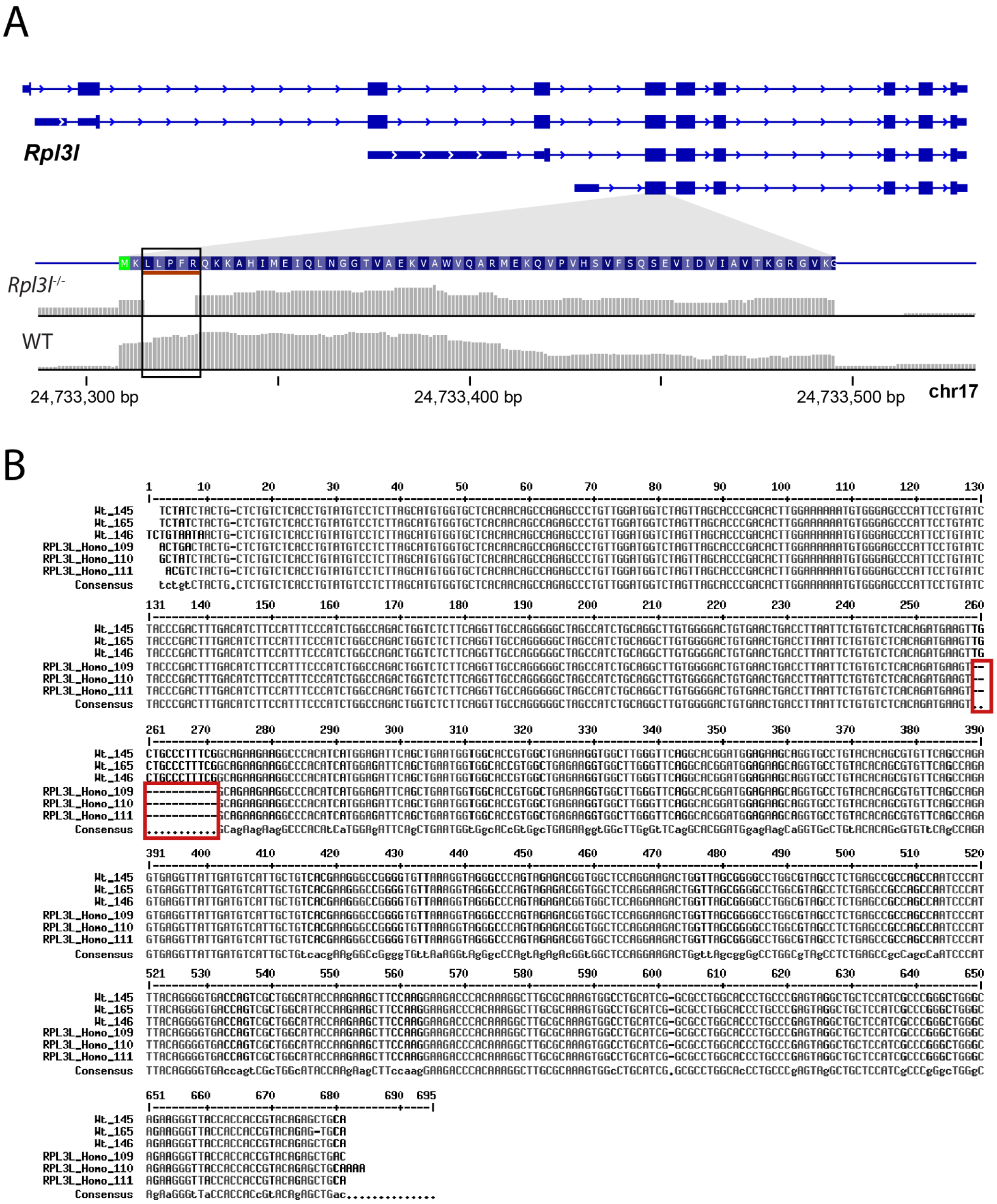

**Figure S2. Imputed expression of ribosomal protein paralogue pairs in heart cells. (A)** *Rpl3* and *Rpl3l* expression profiles in heart cells identified in WT (left) and *Rpl3l*$^{-/-}$ (right) hearts. **(B)** *Rps27* and *Rps27l*. **(C)** *Rpl7* and *Rpl7l1*. **(D)** *Rpl36a* and *Rpl36al*. **(E)** *Rpl22* and *Rpl22l1*. **(F)** *Rpl39* and *Rpl39l*. Expression levels are shown as umi (unique molecular identifiers).

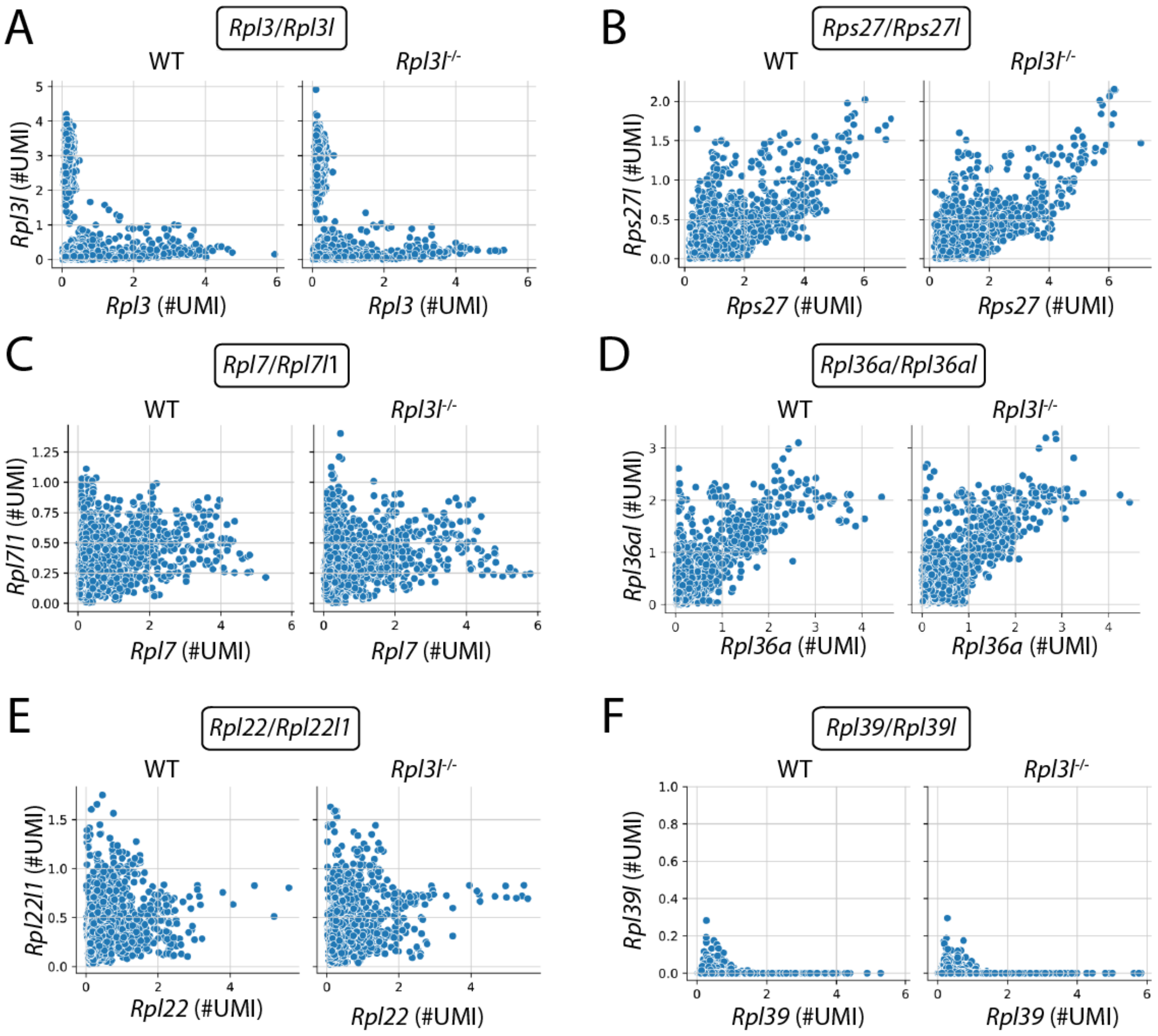

**Figure S3. Chromatin states of the human *RPL3* and *RPL3L* paralogue gene pair (upper panels) and *RPL22* and *RPL22L1* paralogue gene pair (lower panels).** Data was extracted from the WashU Epigenome Browser (1). The vertical colors annotate tissue groups: green - fetal striated muscle, pink - adult striated muscle, blue - adult smooth muscle, purple - other tissues.

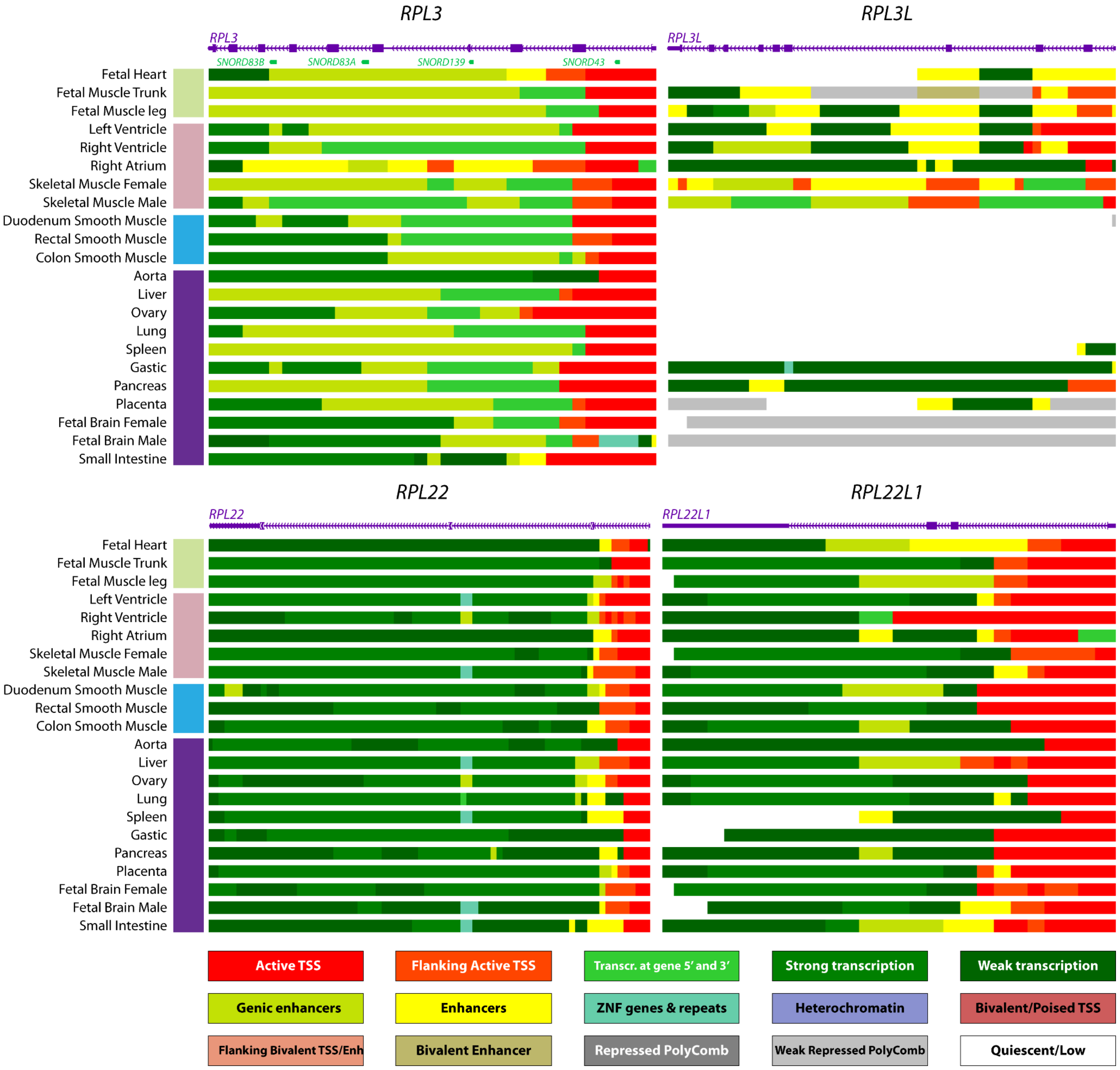

**Figure S4. Western blot analysis of WT and *Rpl3l*$^{-/-}$ cardiomyocytes. (A)** The membrane was probed with an anti-RPL3 (uL3) antibody (left) and reprobed with anti-GAPDH as a loading control (right). **(B)** The membrane was probed with an anti-RPL3L (uL3L) antibody (left) and reprobed with anti-GAPDH as a loading control (right). **(C)** Additional membrane probed with RPL3 (uL3) and GAPDH. **(D)** WT heart and liver protein extracts obtained with RIPA or PEB buffers were probed with the anti-RPL3L (uL3L) antibody. The RPL3L (uL3L)-specific band is only observed in the heart, while it is absent in the liver. Other non-specific bands are present in both tissues.

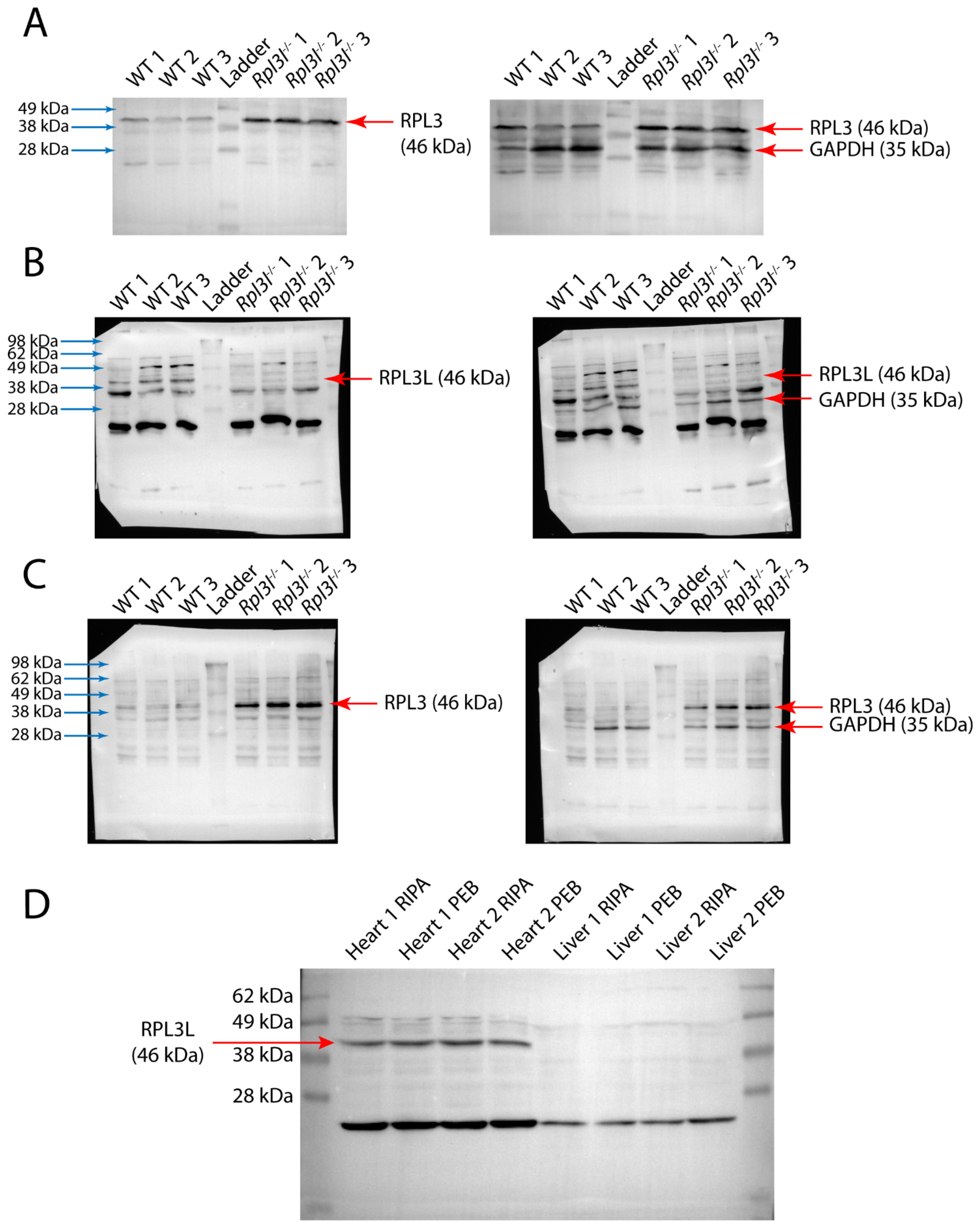

**Figure S5. Western blot analysis of WT and *Rpl3l*$^{-/-}$ hearts.** The membrane was probed with an anti-RPL3 (uL3) antibody and reprobed with anti-GAPDH as a loading control. RPL3 (uL3) levels in WT and *Rpl3l*$^{-/-}$ hearts (n = 3) were assessed densitometrically using ImageJ. Statistical significance was assessed using unpaired t-test (* for $p < 0.05$).

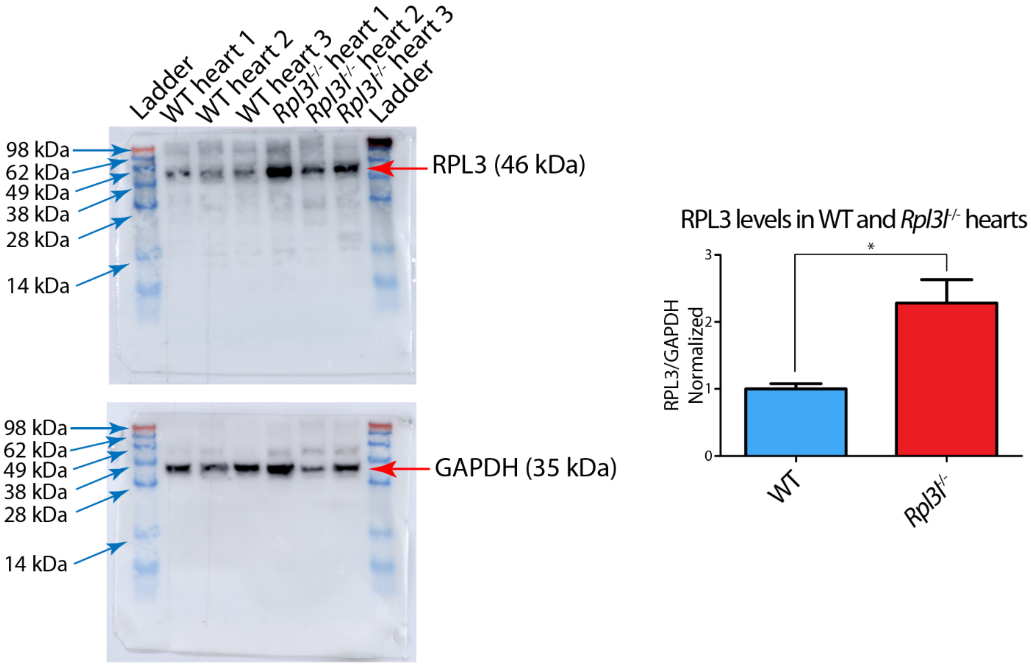

**Figure S6. Western blot analysis of cytosolic and mitochondrial fractions of WT and *Rpl3l*$^{-/-}$ heart lysates. (A)** The membrane was cut and probed with an anti-RPL3L (uL3L) antibody (top), anti-GAPDH as a cytosolic marker (middle) and anti-TOMM20 as a mitochondrial marker (bottom). **(B)** The membrane was cut and probed with an anti-RPL3 (uL3) antibody (top), anti-GAPDH as a cytosolic marker (middle) and anti-TOMM20 as a mitochondrial marker (bottom).

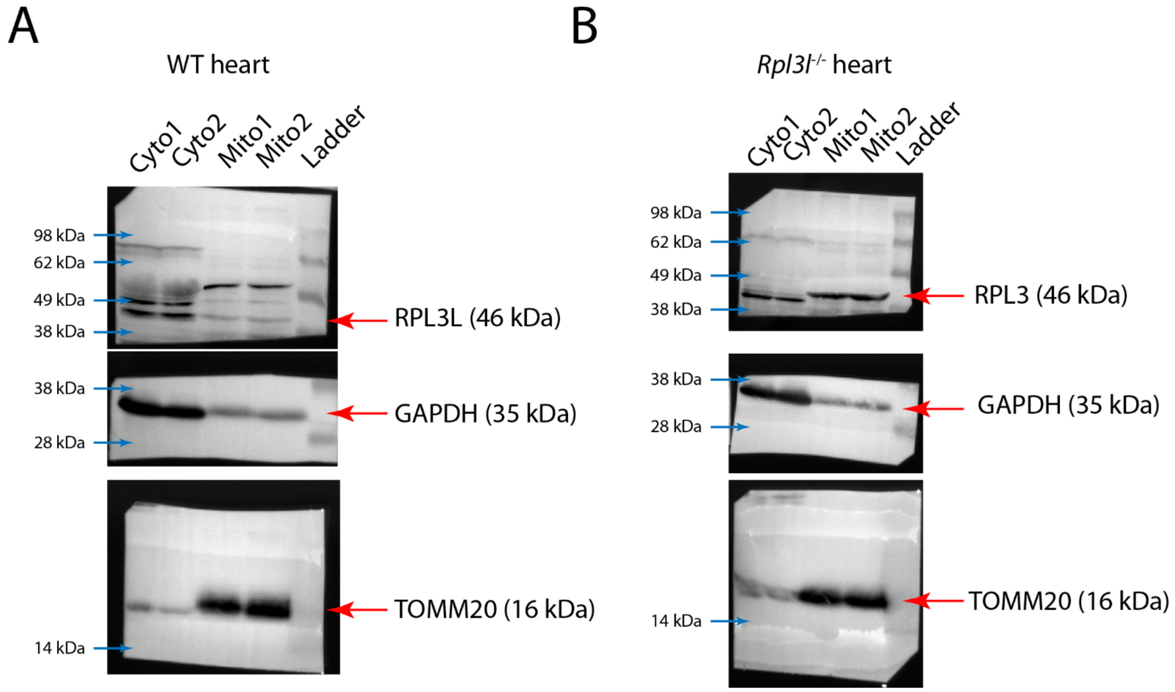

**Figure S7. mRNA expression levels of RPs across tissues and developmental stages in mice.** Embryonic stages are depicted in green (E10.5, E11.5, E12.5, E13.5, E14.5, E15.5, E16.5, E17.5, E18.5), whereas postnatal mice stages are shown in pink (P0, P3, P14, P28, P63). mRNA expression data (RPKM) was obtained from Cardoso-Moreira et al (2).

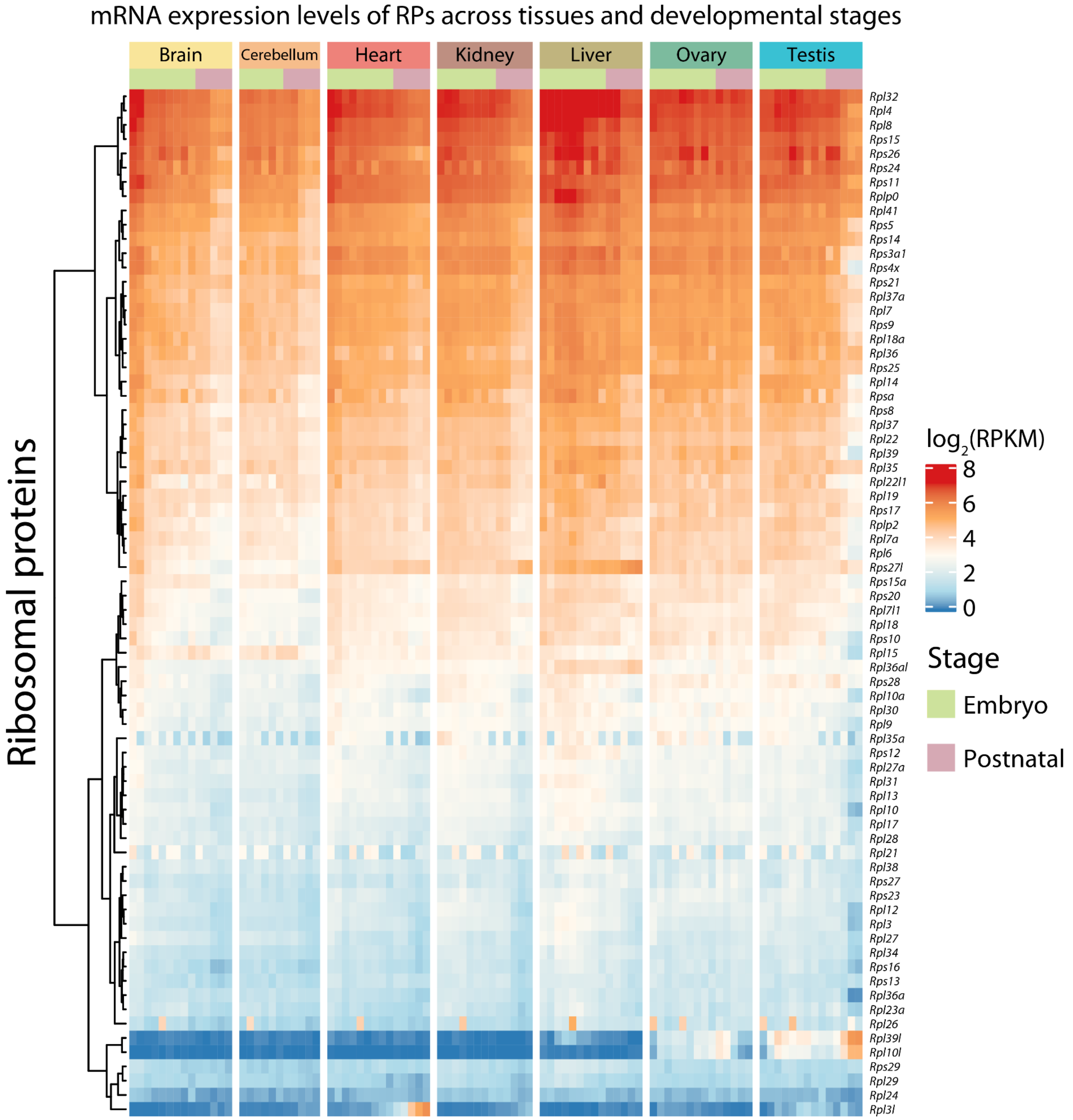

**Figure S8. (A)** Phylogenetic tree of the RPL3 (uL3)/RPL3L (uL3L) family. **(B)** Sequence alignment of human RPL3 (uL3) and RPL3L (uL3L) amino acid sequences. Conserved residues are shadowed in gray, whereas non-conserved ones are shown in white background. The C-terminus region, which is the most distinct between RPL3 (uL3) and RPL3L (uL3L), is boxed. **(C)** Multiple sequence alignment of the C-terminus region from distinct vertebrate RPL3L (uL3L) proteins. The protein length of RPL3L (uL3L) is 407 amino acids in all species.

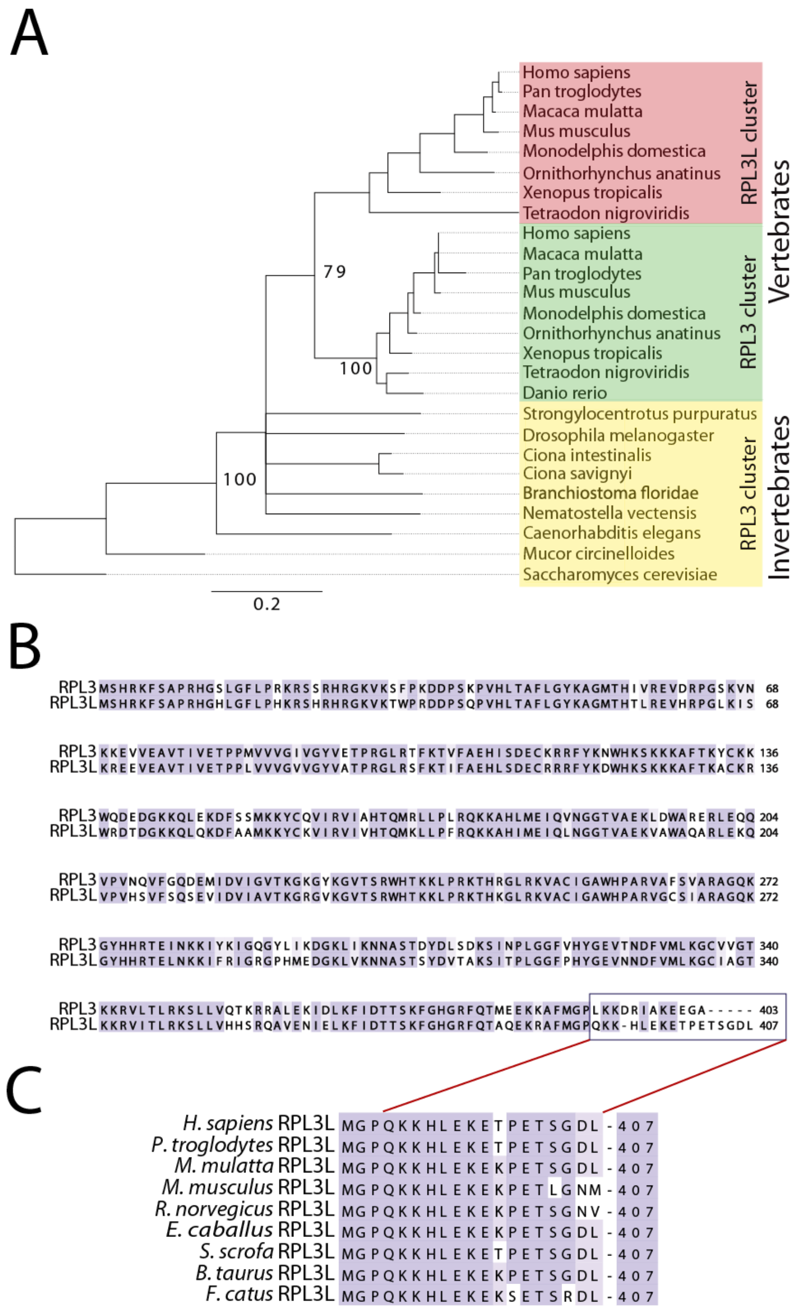

**Figure S9. Histopathological study of heart and skeletal muscle tissues from 8 week old WT and *Rpl3l*$^{-/-}$ male mice. (A)** Hematoxylin and eosin staining of *Rpl3l*$^{-/-}$ gastrocnemius muscles. **(B)** From left to right: heart to body mass ratio, heart to brain mass ratio, left ventricle thickness (mm) and right ventricle thickness (mm), measured on WT and *Rpl3*$^{-/-}$ male mice (n = 5). Statistical significance was assessed using unpaired t-test. **(C)** EchoMRI analysis of 8 week old WT and *Rpl3l*$^{-/-}$ mice in which fat, lean and total weight were measured for each animal (n = 5). Statistical significance was assessed using unpaired t-test.

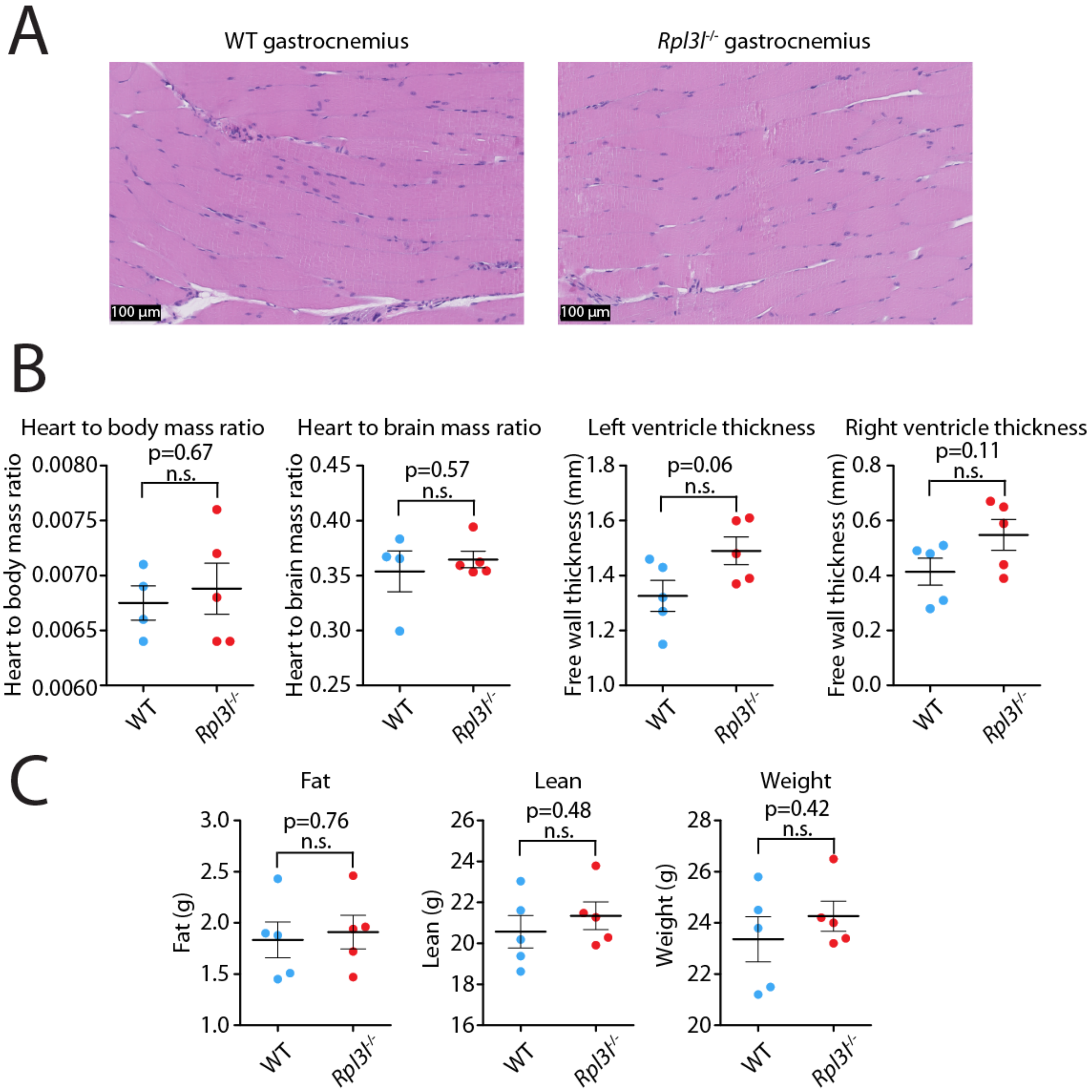

**Figure S10. Echocardiogram and micromanometry profiles of WT and *Rpl3l*$^{-/-}$ mice. (A) Echocardiogram parameters.** Abbreviations: HR - Heart rate; EDV - Left ventricle (LV) chamber volume at end diastole; ESV - LV chamber volume at end systole; SV - volume ejected during systole; CO - cardiac output; EF - ejection fraction; Vcf - fiber shortening velocity; h - Mean LV tissue wall thickness at the mid-papillary level at end diastole; r - Mean LV chamber radius at mid-papillary level at end diastole; h/r - ratio of wall thickness to chamber radius et end diastole; LVd wall mass - mass of the LV wall; LA - diameter of the left atria at end systole; E/A - early to atrial mitral Doppler velocity ratio; E/e' - early pulse wave velocity to mitral annulus velocity; IVCT - time interval from the mitral valve closure; IVRT - isovolumic relaxation time; ET - ejection time; AV peak vel - aortic valve peak velocity. **(B) Micromanometry parameters.** Abbreviations: AoPs - aortic systolic pressure; AoP mean - aortic pressure mean; AoPd - aortic diastolic pressure; LVPs - left ventricular systolic pressure; dP/dt max - maximal rate of rise of LV pressure; dP/dt min - minimal rate of rise of LV pressure; EDP - end diastolic pressure. Statistical significance was assessed using unpaired t-test (* for $p < 0.05$).

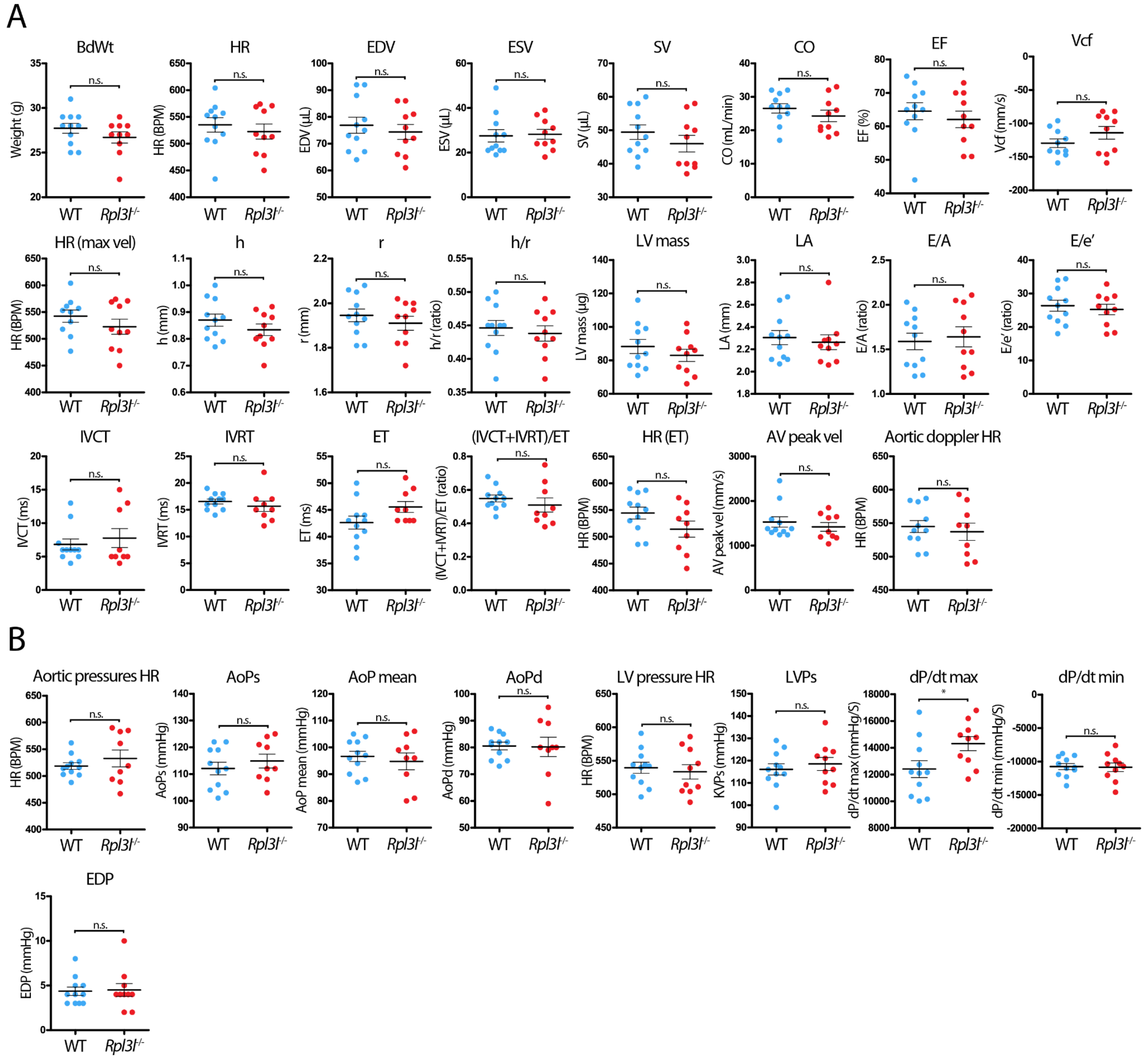

**Figure S11. Replicability of the Nano-TRAP sequencing.** Scatterplots of logarithm of raw counts obtained from Nano-TRAP sequencing of hearts from biological replicates of WT and *Rpl3l*$^{-/-}$ mice (n = 3). (**A**) Reads from the IP experiment (Ribo-bound). (**B**) Reads from the input experiment. Hemoglobin genes are coloured in gray. (**C**) Correlation of mean counts from WT and *Rpl3l*$^{-/-}$ samples from the input RNA (left), the immunoprecipitated ribosome-bound RNA (middle) and translation efficiency (right).

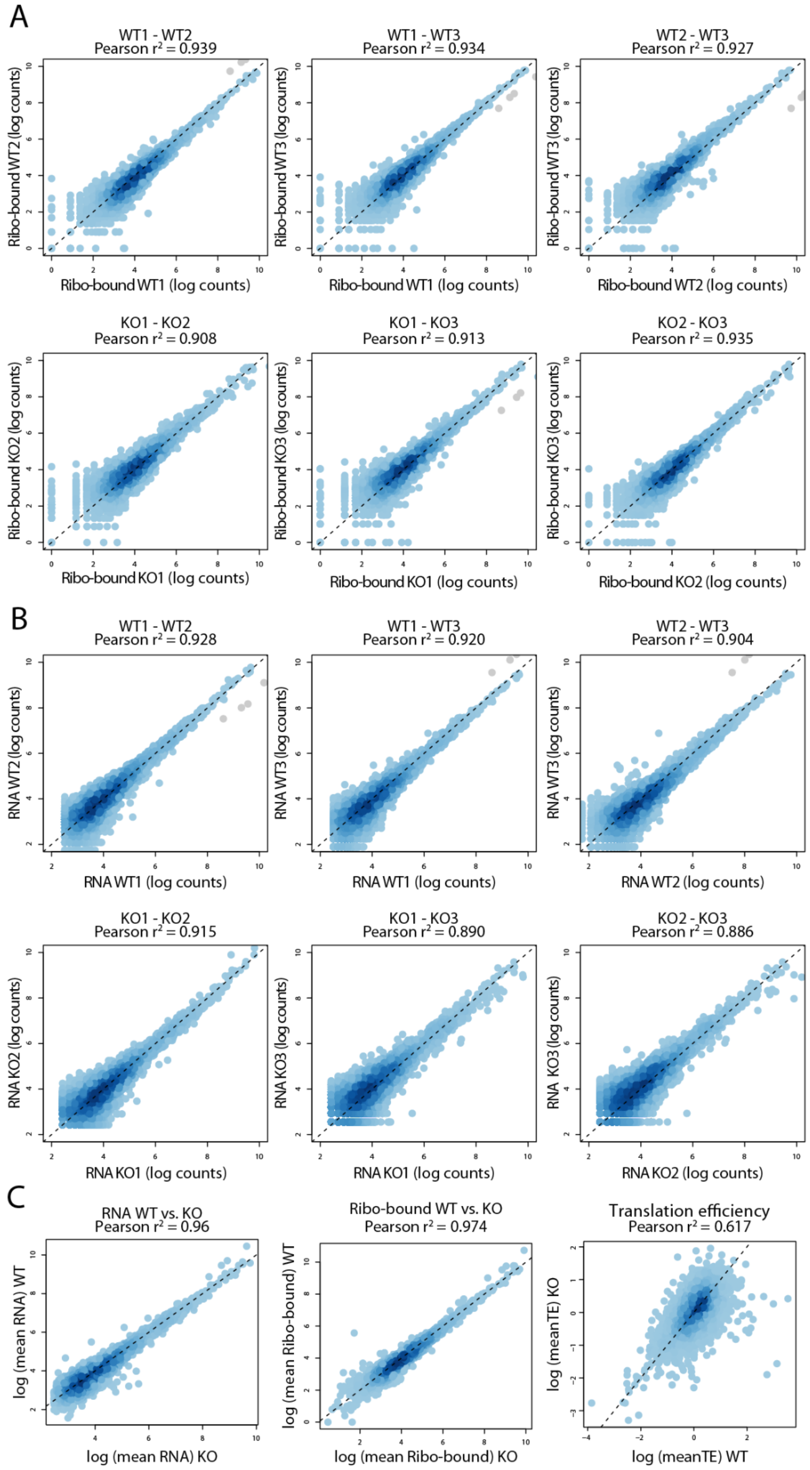

**Figure S12. Replicability of the Ribosome profiling experiments.** Scatterplots of logarithm of raw counts obtained from Ribo-Seq sequencing of hearts from biological replicates of WT and *Rpl3l*$^{-/-}$ mice (n = 3). (**A**) Reads from the Ribo-Seq experiment (RPFs). (**B**) Reads from the input (RNA-Seq) experiment.

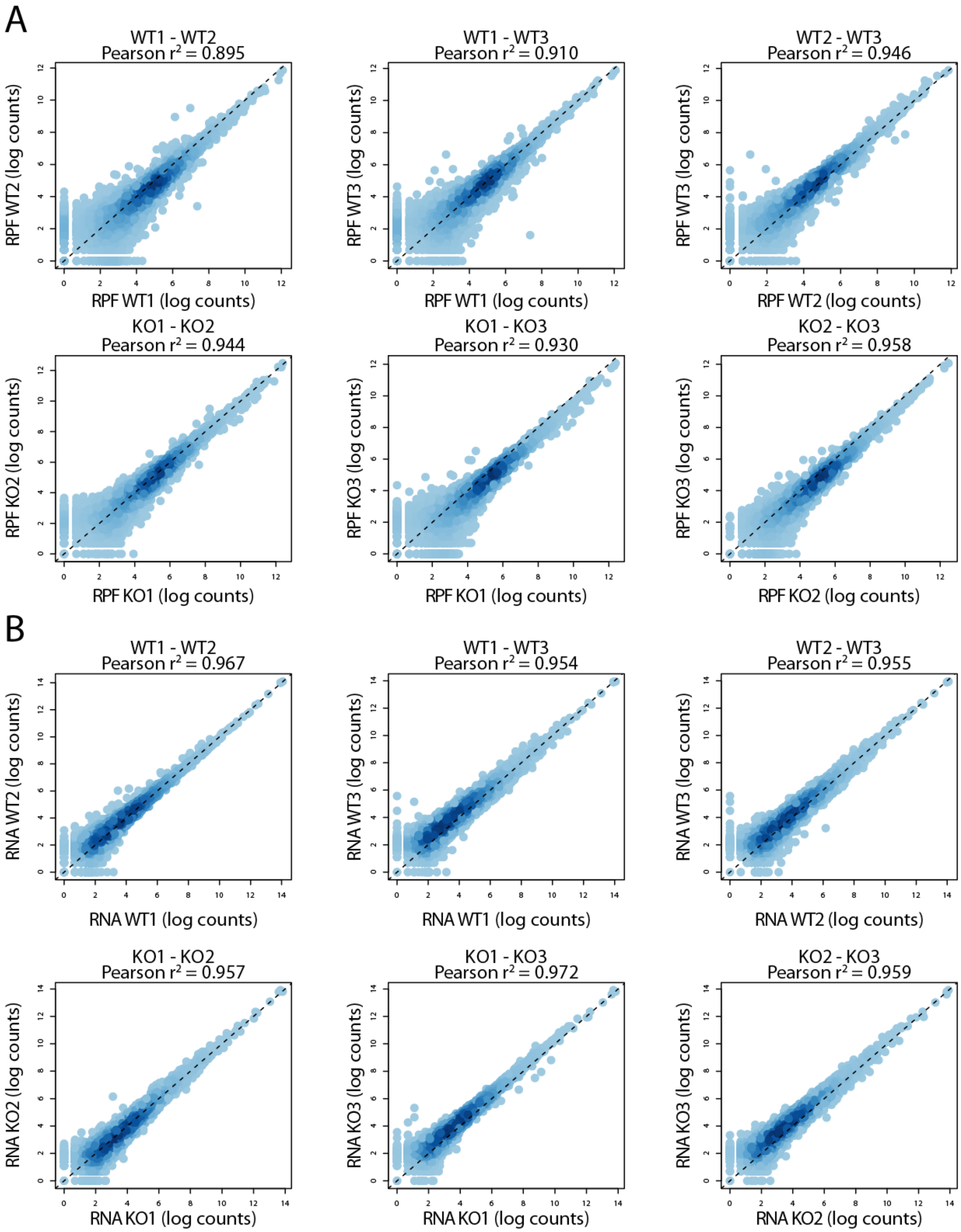

**Figure S13. Ribosome profiling quality metrics. (A)** Start site (left) and stop site (right) coverage showing 3-nucleotide periodicity in RiboSeq reads. **(B)** Codon occupancy differences between WT and *Rpl3l*$^{-/-}$ ribosomes centered at A-sites (top) and E-sites (bottom). **(C)** Volcano plot representing genes with differential abundance of genes (left), and genes with differential abundance of ribosome footprints (RPF) (right). Red dots indicate genes whose log2 fold change in normalized RPF counts is greater than 1 and FDR adjusted p-value lower than 0.05. **(D)** TE analysis showing the correlation of TE in WT and *Rpl3l*$^{-/-}$ hearts.

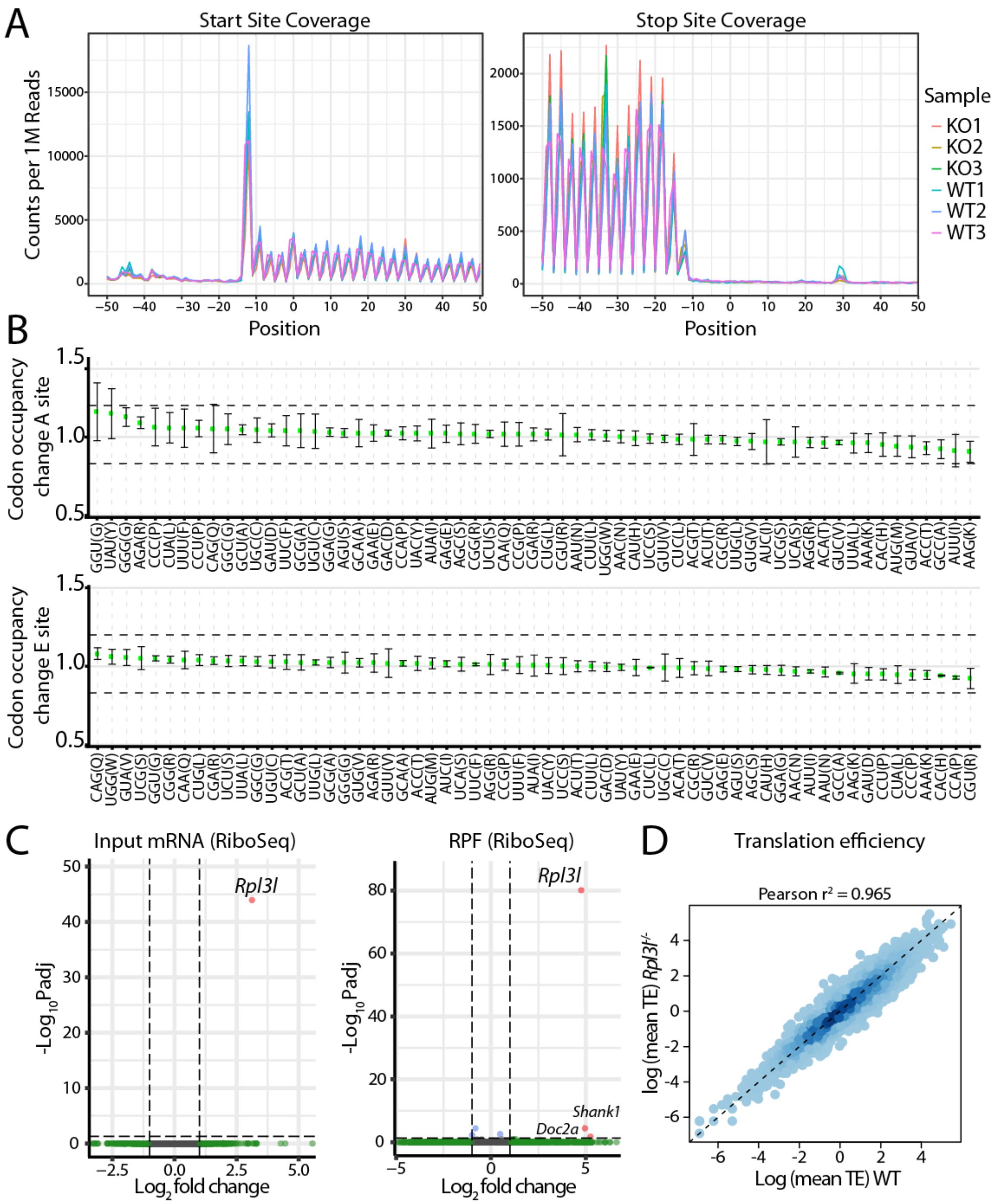

**Figure S14. Ribosome profiling metagene plots.** Metagene plots representing the RPF reads distribution across the 5' UTR, CDS and 3' UTR regions.

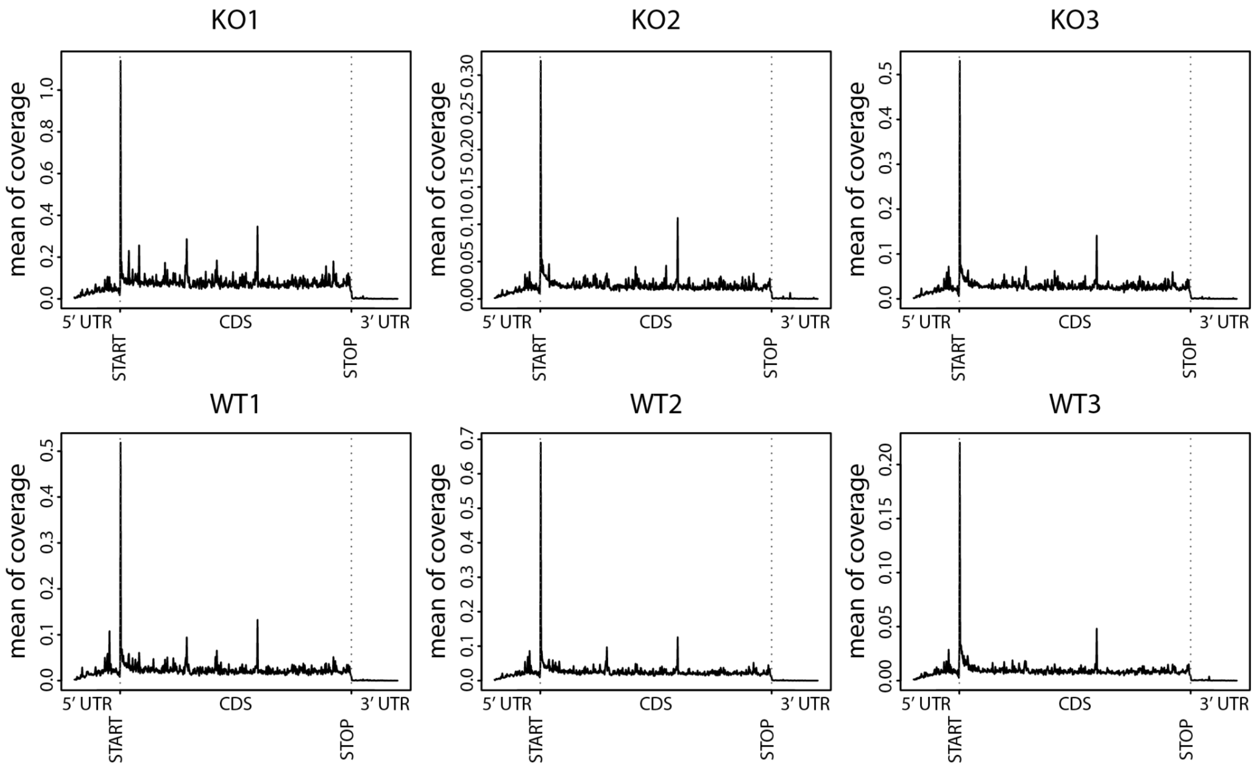

**Figure S15. *Rpl3l* and *Rpl3* counts in RiboSeq and Nano-TRAP experiments. (A)** *Rpl3l* and *Rpl3* counts detected in the Ribo-Seq and RNA-Seq (input) experiments (n = 3). **(B)** *Rpl3l* and *Rpl3* counts detected in the Nano-TRAP (IP and Input) experiments (n = 3).

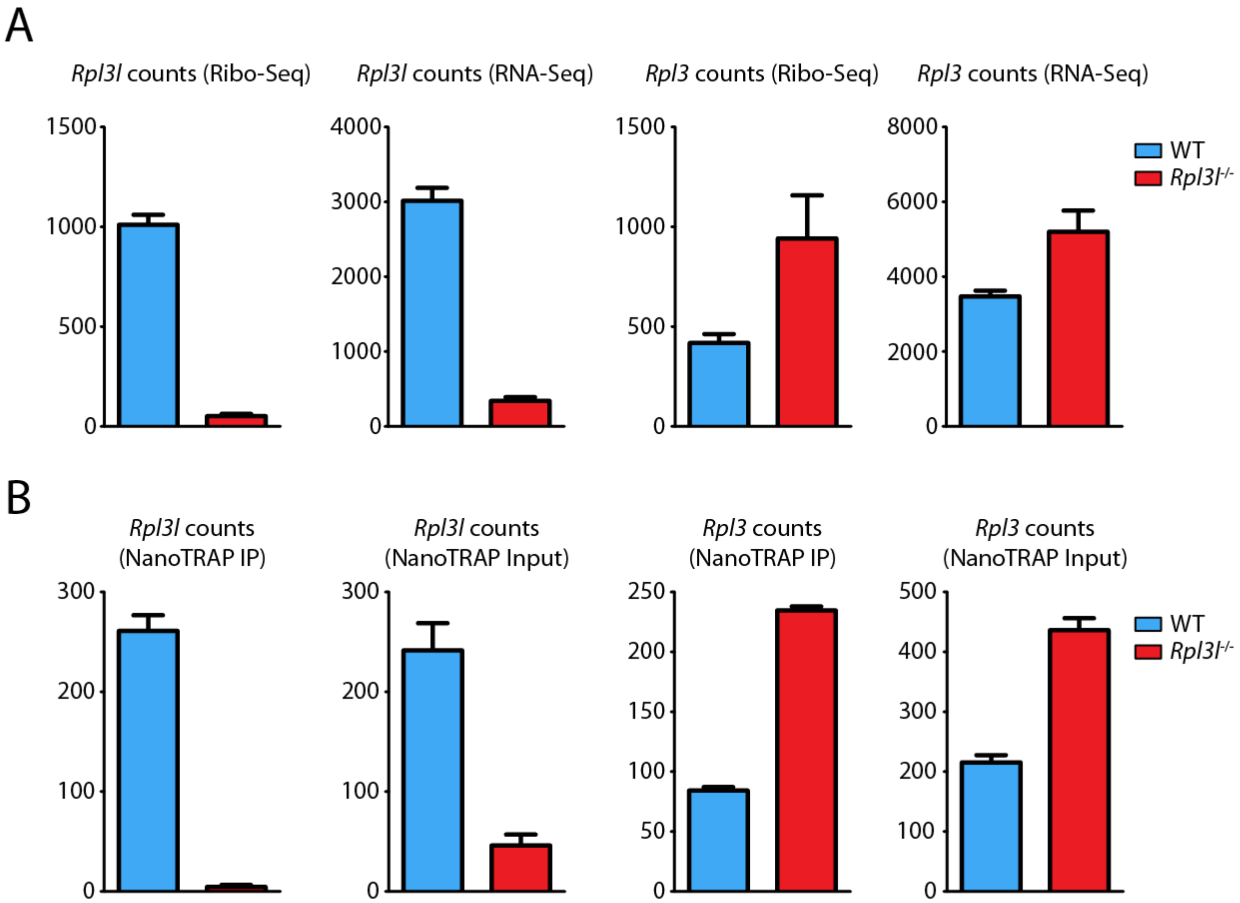

**Figure S16. *Rpl3l* is cardiomyocyte-specific and its expression is mutually exclusive with *Rpl3*. (A)** Overview of the UMAP clusters of cell types identified in heart left ventricles using single nuclei RNA-Seq (sNuc-seq) (left). In the middle and right panels, the expression levels of *Rpl3* (middle) and *Rpl3l* (right) for each of the cells in the clusters. Expression values represent umi (unique molecular identifiers). **(B)** Analysis of the *Rpl3-Rpl3l* expression in single cells from a WT heart, coloured by cell type, showing that *Rpl3* and *Rpl3l* expression is mutually exclusive. **(C)** *Rpl7-Rpl7l1* expression in single cells from a WT heart, coloured by cell type. Expression levels shown in panels B and C have been imputed using the MAGIC algorithm. Abbreviations: CM (cardiomyocytes), EC (endothelial cells), FB (fibroblasts), MP (macrophages), PC (pericytes), SMC (smooth muscle cells).

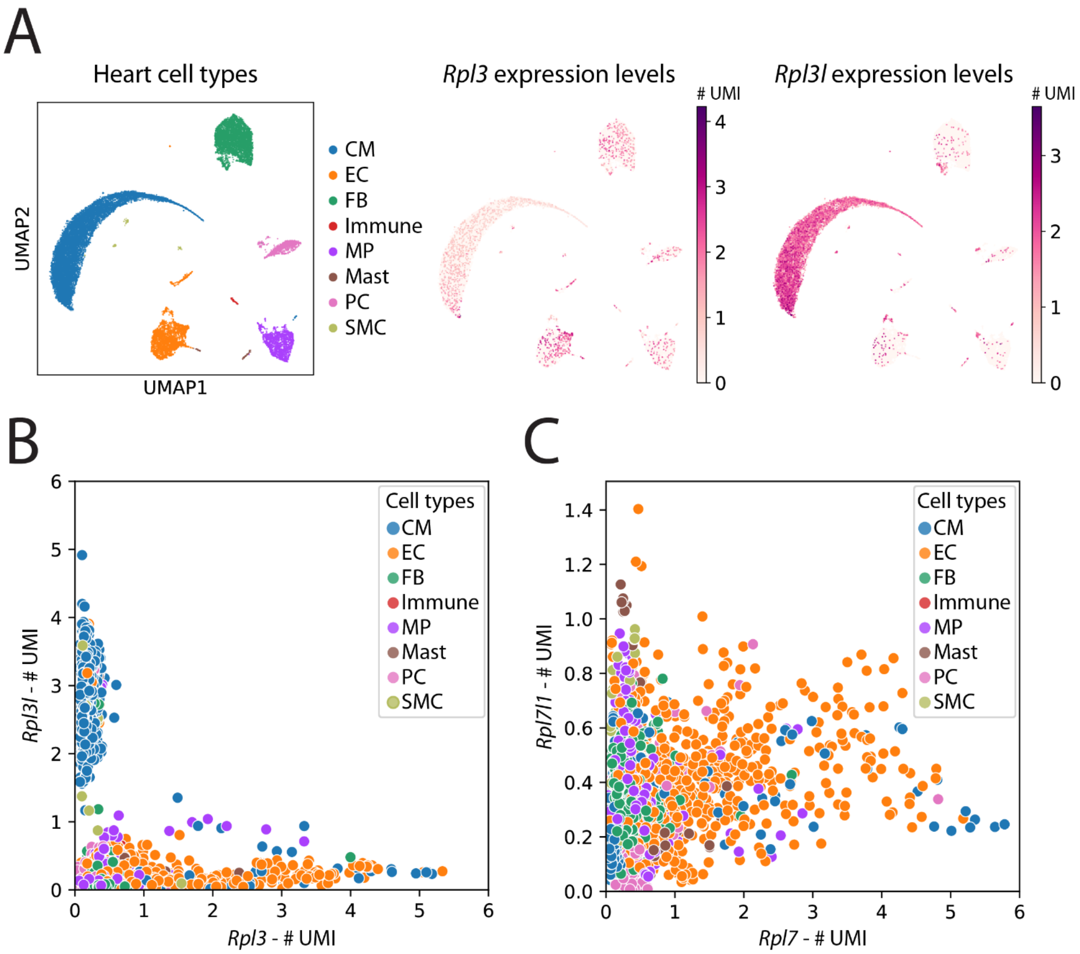

**Figure S17. Proteo-TRAP replicability among biological replicates.** Scatterplots showing the replicability of the Proteo-TRAP experiment among biological replicates and across WT and *Rpl3l*$^{-/-}$ samples. Every dot represents a protein. Mitochondrial proteins are colored in red, and non-mitochondrial proteins in gray.

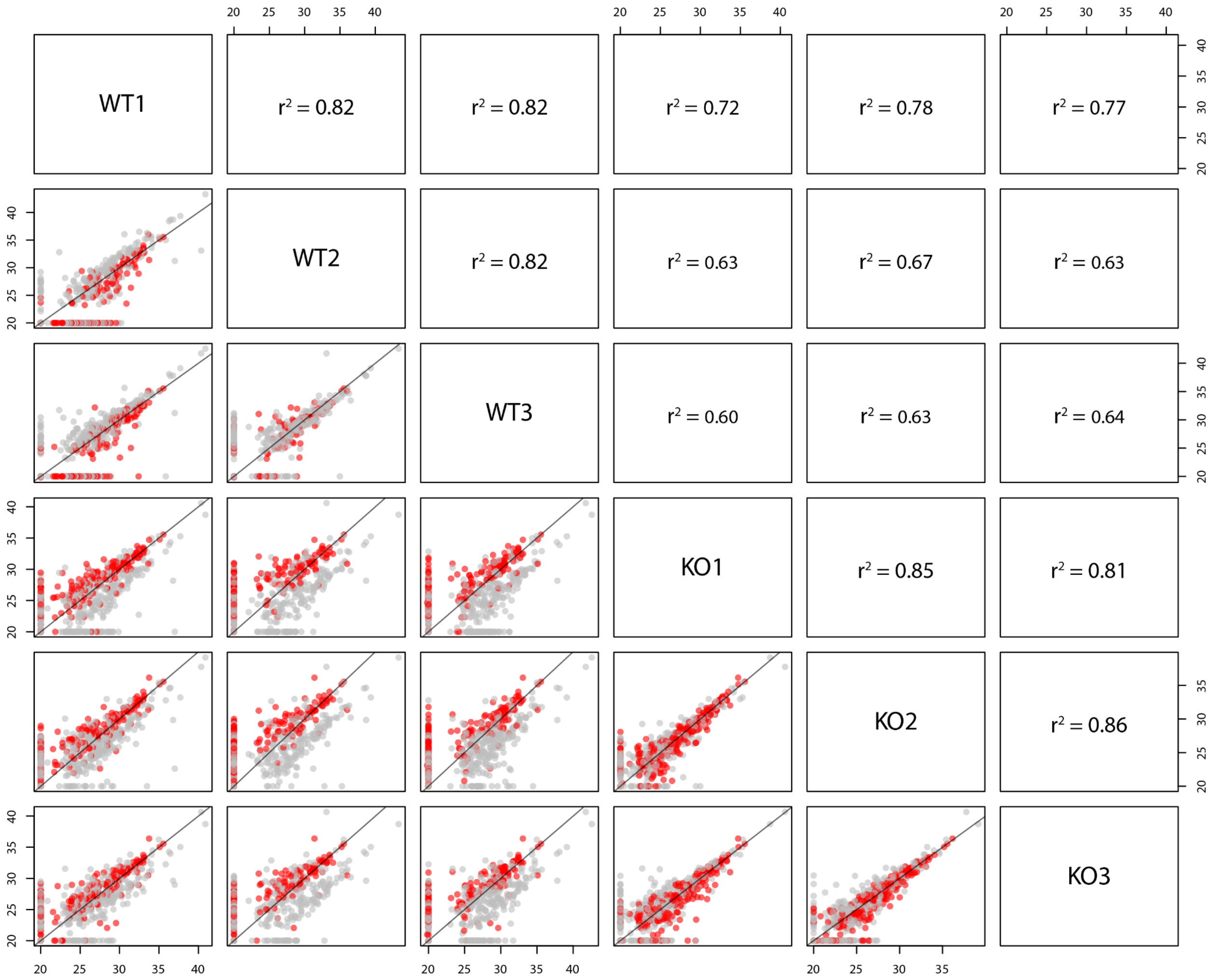

**Figure S18. *Rpl3l* depletion does not affect the abundance of mitochondria. (A)** Quantification of mitochondrial DNA by qPCR, normalized by genomic DNA. Mitochondrial copy number was assessed by the qPCR quantification of 16S mitochondrial ribosomal RNA and the NADH-ubiquinone oxidoreductase chain 1 (*Nd1*) encoded by the mitochondrial genome, both normalized by hexokinase-2 (*Hk2*), encoded by the nuclear genome. Left: 16S/*Hk2* ratio, right: *Nd1*/*Hk2* quantification. **(B)** Immunofluorescence of WT and *Rpl3l*$^{-/-}$ heart tissues, stained with Phalloidin (green, showing actin), anti-ATP5A (red, showing mitochondria) and DAPI (blue, showing nuclei).

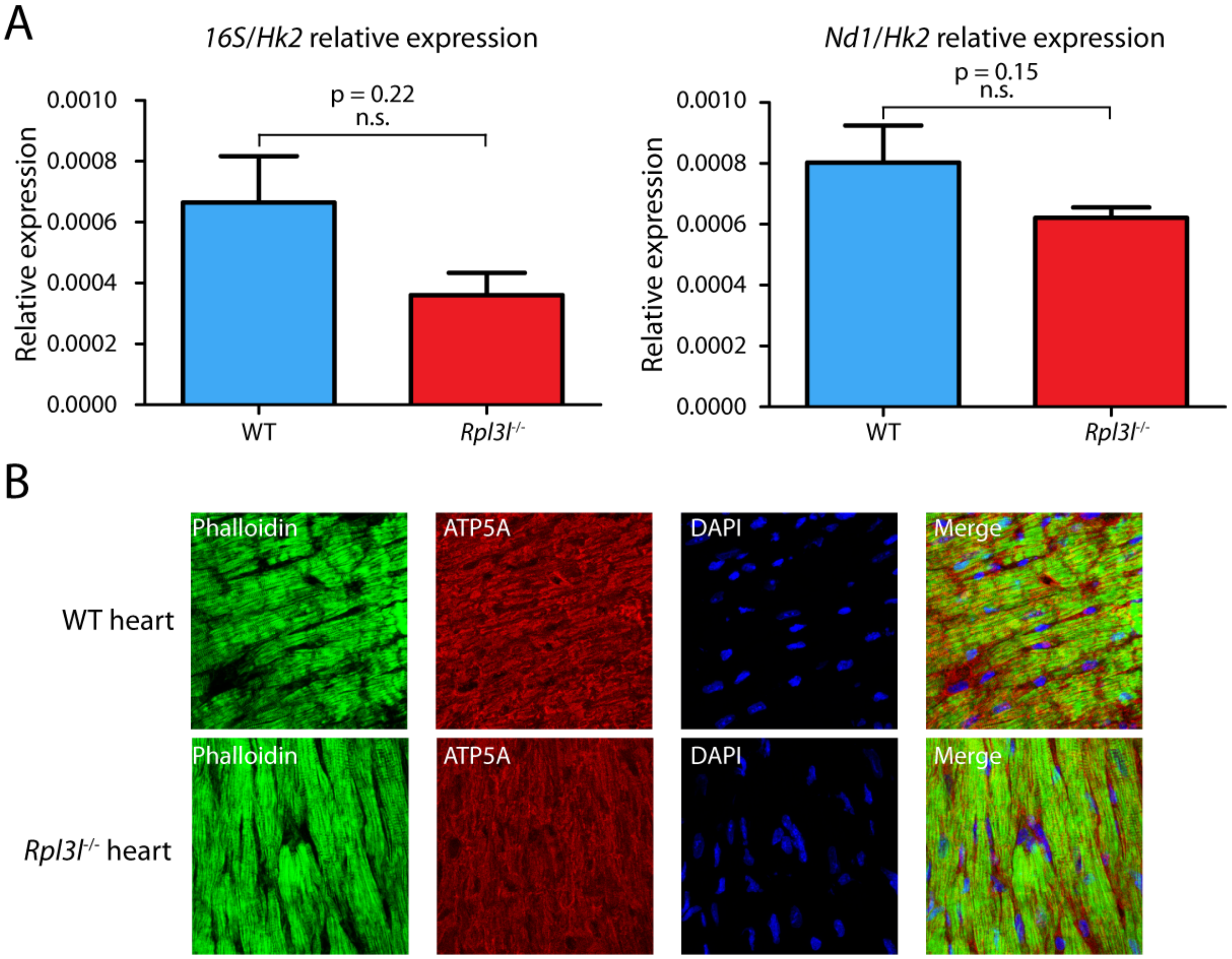

**Figure S19. Single cell RNA-Seq of a time-series of TAC experiments.** The expression of *Rpl3* and *Rpl3l* across cell types in mouse hearts before (week 0) and after hypertrophy induced by transverse aortic constriction (week 2, 5, 8, 11). Expression levels are shown as umi (unique molecular identifiers). Abbreviations: CM (cardiomyocytes), EC (endothelial cells), FB (fibroblasts), GN (granulocytes), MP (macrophages), T (T lymphocytes). Data for generating plots has been taken from Ren et al., 2020 (3). Cardiomyocyte clusters are circled.

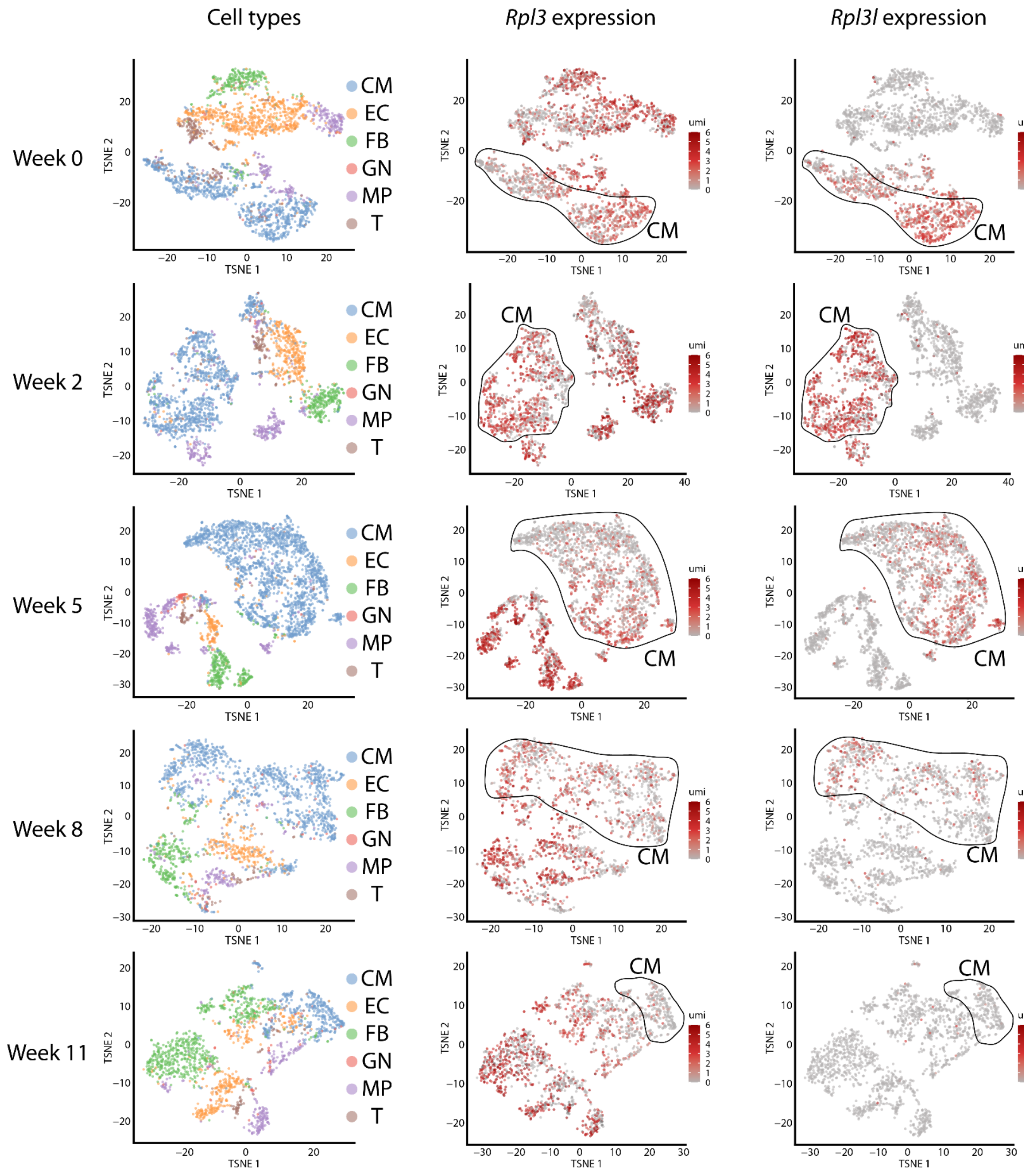

**Figure S20. *Rpl3-Rpl3l* interplay in hypertrophy. (A)** RNA-seq RPKM values of *Rpl3* and *Rpl3l* in hypertrophic (TAC) and control (sham) hearts 2, 4, 7 and 21 days after surgery. **(B)** Ratios of RPKM values of all RPs in TAC and sham conditions across the time-series. *Rpl3l* and *Rpl3* are coloured in blue and red, respectively. RNAseq data was taken from Wang et al.(4).

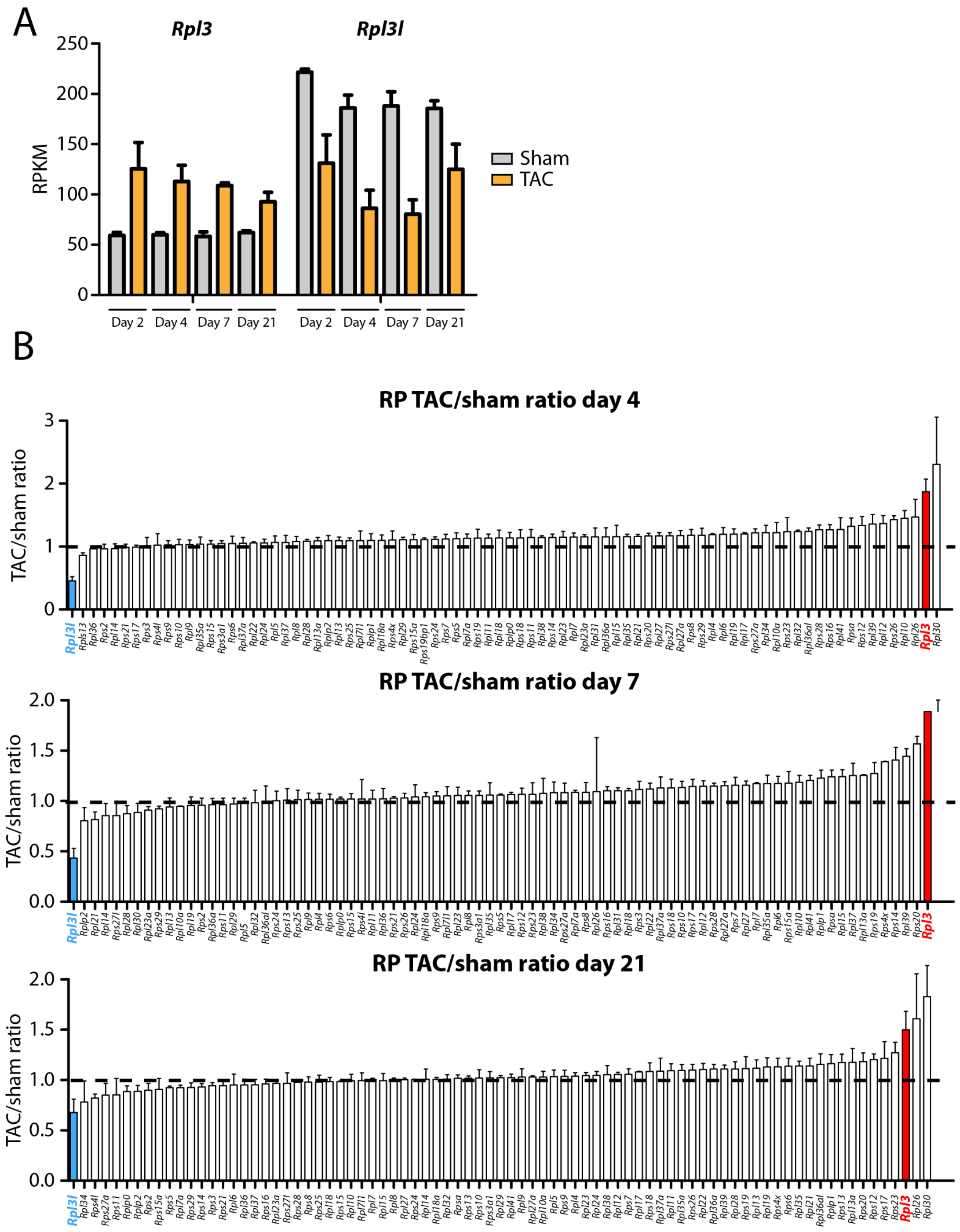

**Figure S21. Analysis of the dynamics of mRNA expression levels (RPKM) of mouse RP paralog pairs upon cardiac hypertrophy.** Ratio of RNA-seq RPKM values of RP paralog pairs (*Rpl7/Rpl7l1, Rps27/Rps27l, Rpl36a/Rpl36al* and *Rpl22/Rpl22l1)* in hypertrophic (TAC) and control (sham) hearts. *Rpl10/Rpl10l* and *Rpl39/Rpl39l* pairs were not included in the analysis because one of the paralog proteins was absent in all the conditions. See Figure 6B for *Rpl3/Rpl3l* pair. RNAseq data was taken from Wang et al.(4).

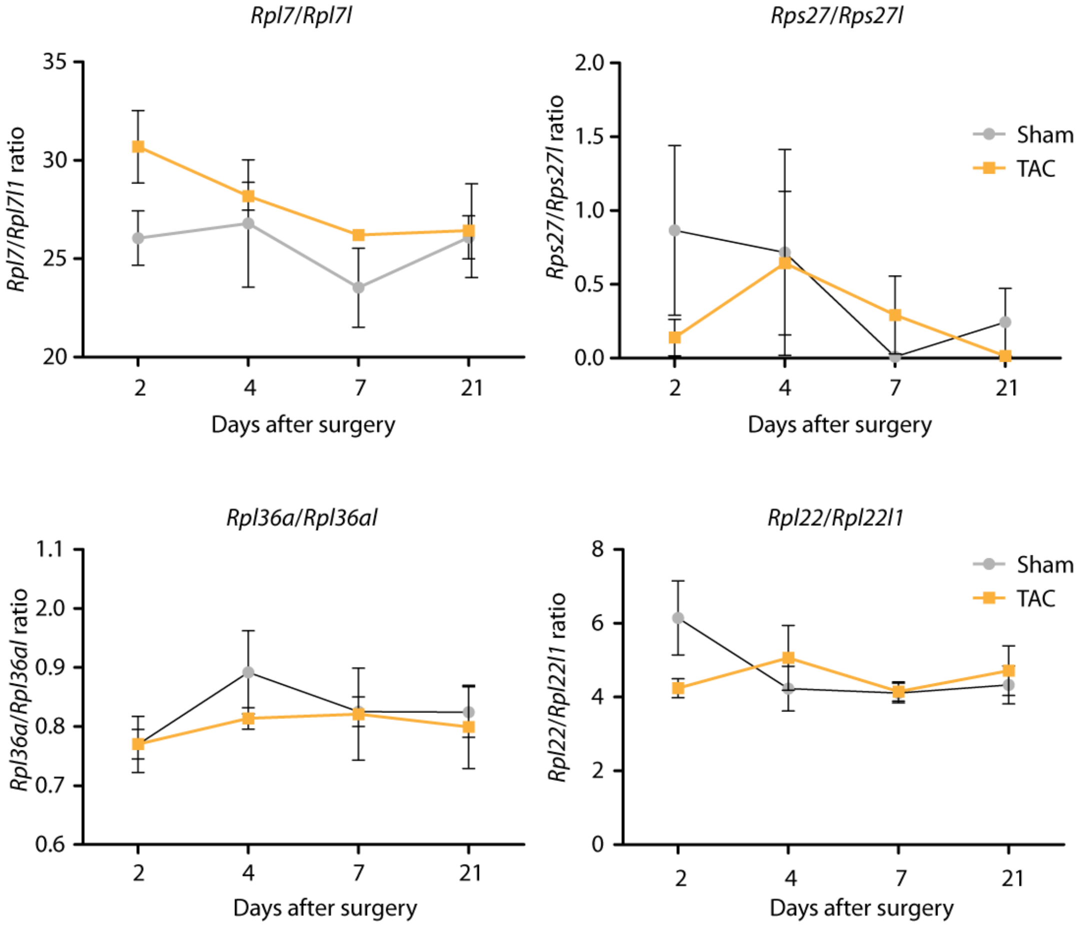

**Figure S22. *Rpl3-Rpl3l* interplay in hypertrophy is impaired in *Lin28a*$^{-/-}$ mice. (A)** RNA-seq RPKM values of *Rpl3* and *Rpl3l* in hypertrophic (TAC) and control (sham) hearts of WT and *Lin28a*$^{-/-}$ mice. **(B)** Ratios of RPKM values of all RPs in TAC and sham conditions for WT (top) and *Lin28a*$^{-/-}$ (bottom) mice. *Rpl3l* and *Rpl3* are coloured in blue and red, respectively. RNAseq data was taken from Ma et al.(5)

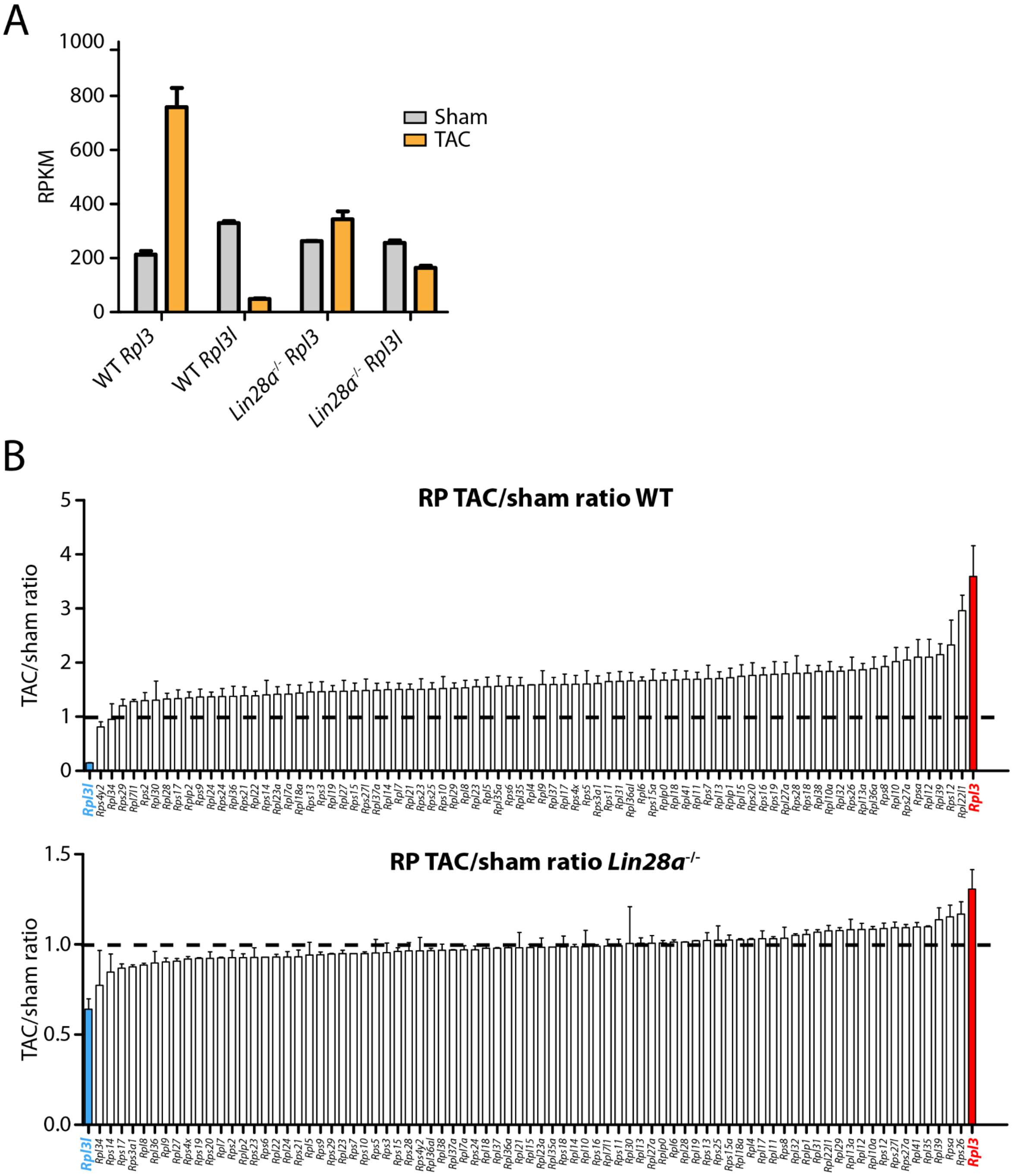

**Figure S23. Translation efficiency calculated using deltaTE. (A)** Volcano plot showing the translation efficiency calculated using deltaTE (6) in WT vs *Rpl3l*$^{-/-}$ samples. **(B)** Volcano plot showing the translation efficiency using deltaTE in control and PP242-treated PC3 cells (data for generating this plot was taken from (7)). Each dot represents a gene, and they have been coloured depending on: i) adjusted p-value < 0.05 and fold change > 1 (red), ii) only fold change > 1 (green) or only adjusted p-value < 0.05 (blue) , iii) neither fold change > 1 nor adjusted p-value < 0.05 (gray).

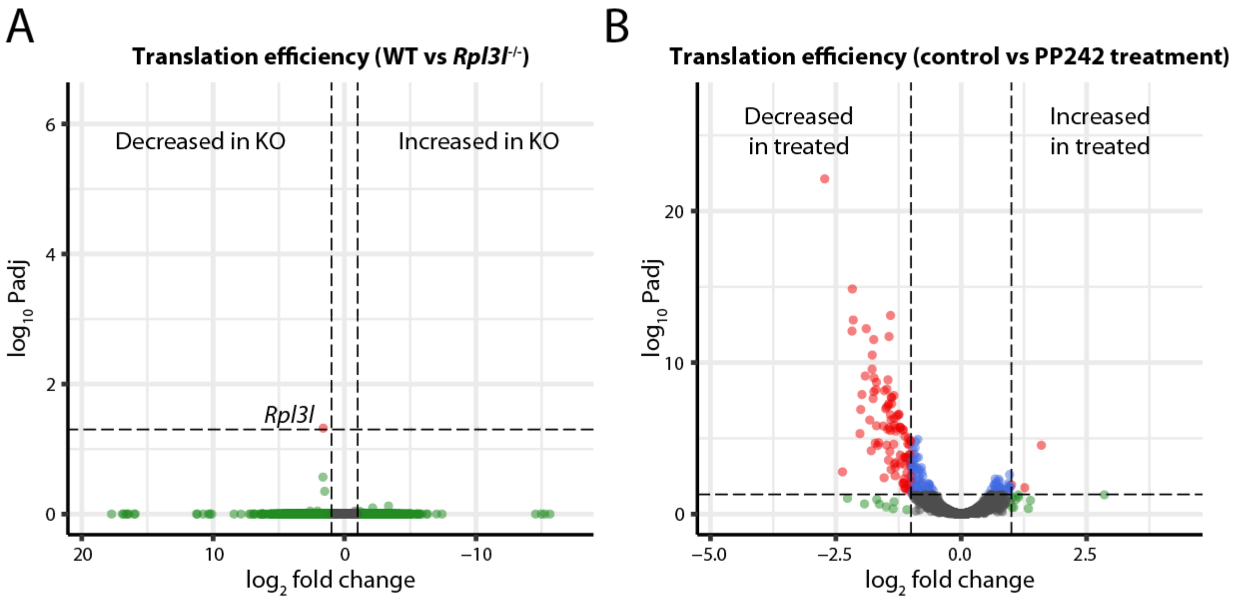
